# Fifteen-year microbiome survey of endangered killer whales (*Orcinus orca*) reveals declining diversity and population differences

**DOI:** 10.64898/2025.12.27.696689

**Authors:** Alexandra D. Switzer, Kim M. Parsons, M. Bradley Hanson, Candice Emmons, Linda Park, Jennifer Hempleman, Todd Robeck, Kelsey Herrick, Steven Osborn, Alexander L. Jaffe, Dan Olsen, Craig Matkin, Elisabeth M. Bik, Anna Robaczewska, Abigail Wells, Darran May, Samuel Wasser, Sheila J. Thornton, David A. Relman

## Abstract

The endangered Southern Resident killer whale (*Orcinus orca*) (SRKW) population is burdened by multiple anthropogenic stressors with limited non-invasive approaches for health surveillance. The gut microbiome is interconnected with host physiology and can be characterized using remotely collected fecal samples, creating unique opportunities to integrate environmental and individualized host-associated features of health. In this study, we used fecal samples collected over a 15-year period (2005-2019) from 77% of the living, wild SRKW population (56 individuals, 1-17 time points per individual), as well as fecal samples from the conspecific Northern and Alaska Resident killer whale populations, to characterize distal gut microbiota structure and genomic composition. SRKW microbiotas were individualized and distinct from those of the nearby Northern and Alaska Resident populations, both of which exhibit consistently higher fecundity and survivorship. During the study period, SRKW fecal microbiota species richness declined, despite stability over periods of time less than or equal to 1 year. Several potential bacterial pathogens, such as *Fusobacterium* spp., achieved dominance in the fecal microbiotas of SRKW individuals sampled within 6 months of death. These findings demonstrate the feasibility and value of harnessing non-invasively collected fecal samples and microbiome profiles for longitudinal killer whale health surveillance.

---

The critically endangered Southern Resident killer whale (SRKW) population faces numerous anthropogenic stressors. Accordingly, there is a need to improve non-invasive health surveillance approaches^1^. After being heavily impacted by the marine park live-capture industry from 1961-1972^2^, the SRKW population, located in waters off the Pacific northwest coast of the United States, experienced nearly two decades of population growth and recovery followed by a subsequent period of high mortality and low fecundity in the 1990s^3^ resulting in a depleted and vulnerable population. As a result, the SRKW population was federally listed as endangered under the U.S. Endangered Species Act in 2005^4^ and under the Species at Risk Act (SARA) in Canada in 2003^3^. As of the 2024 census, only 73 individuals remained^5^. Despite concerted efforts aimed at habitat recovery, minimizing anthropogenic disturbance, and amplifying available prey species, the SRKW population has failed to grow consistently^1,6,7^. The diminished SRKW population size has resulted in reduced fitness due to inbreeding depression, the effects of which are expected to worsen should their numbers continue to decline^8^. In contrast, two other Northeastern Pacific resident killer whale (KW) populations, the Northern (NRKW, >300 individuals^9^) and Alaska (ARKW, >700 individuals^10^) residents, consistently exhibit higher fecundity^11^, survivorship^12,13^, and genetic diversity^8^, despite sharing similar life histories and salmonid prey preferences. The sympatric NRKW population is an especially useful comparator for SRKWs given overlapping range and diet^7,14^. Both populations face similar anthropogenic pressures, although SRKWs may be disproportionately affected due to their accessibility to humans and proximity to heavily urbanized waterways^15^.

Four primary threats are thought to contribute to the decline and lack of recovery of the SRKW population: (1) scarcity of their preferred prey, Chinook salmon^16,17^; (2) acoustic and physical disturbance from vessels in the Salish Sea^18,19^; (3) environmental contaminants^20–22^; and (4) inbreeding^8^. SRKWs have one of the highest burdens of polychlorinated biphenyls (PCBs) among marine mammals^20,23^. Despite a global ban, PCBs are expected to continue to have toxic effects in the SRKW population through 2063. Primary threats to the SRKW population often interact cumulatively and synergistically. For example, anthropogenic noise may reduce foraging success^24^, which exacerbates nutritional stress, resulting in body fat metabolism and mobilization of lipophilic PCBs^19,23,25^. Nutritional stress may be a significant cause of SRKW low reproductive success, and has been identified as a potential proximate cause of late-term spontaneous abortions^11,17^. Additionally, the energetic demands of lactation accelerate the release of PCBs from maternal fat into circulation, potentially contributing to perinatal or neonatal mortality^11,21^. The preponderance of SRKW conservation research, including population viability analyses, has focused on this triad of primary extrinsic threats-but the interplay between these external stressors and host-associated microbial communities remains unexplored.

Host health is intrinsically tied to that of the gut microbiome, and while gut microbial communities generally demonstrate temporal stability, compositional shifts can cause, or reflect, illness^26^. Marine mammal microbiomes are relatively underexplored, although prior studies demonstrate host species specificity and an effect of provenance, i.e., life in the sea^27^. A few studies have indicated temporal stability of the distal gut microbiota of managed cetaceans (dolphins^27–29^, belugas^30^ and pilot whales^31^) and individuality. Toxin exposure, stress, and malnutrition have all been correlated with changes in the gut microbiome in terrestrial mammals, sometimes in a bidirectional manner. For example, exposure of mice to PCBs can alter gut community structure, and in turn these communities can mediate PCB-induced hepatic toxicity^32^. Prolonged strenuous exercise in humans has been associated with reduction of normal dominant taxa in fecal communities and an increase in intestinal permeability^33^. Malnutrition has been correlated with changes in the fecal microbiomes of children, such as loss of diversity and enrichment for enteric pathogens^34^. It follows that the SRKW gut microbiome may be similarly affected by these extrinsic stressors. However, the lack of pre-existing knowledge about normal baseline KW gut microbial community composition hinders the identification of features reflective of health and disease.

The development of non-invasive SRKW health surveillance methods is a priority since wild KWs are sensitive to vessel disturbance^11^ and direct handling can exacerbate stress^35,36^. Fortunately, long-term directed research has yielded methods to retrieve wild KW fecal samples remotely from the sea within minutes of defecation^11,37^. Since the entire SRKW population has been surveyed annually for more than 40 years^5^ and host DNA has been archived for most members^38^, fecal samples can be genetically assigned to individual SRKWs based on host DNA sequences from these samples. Fecal samples provide a non-invasive method for characterizing the SRKW distal gut microbiome and establishing a reference baseline for the population and individuals over time and ultimately could allow the identification of ‘normal’ versus ‘abnormal’ signatures or trends associated with host health status or survival outcome, as well as emerging pathogens.

Here, we examined the relationship between the gut microbiota and SRKW health through both a population-level and individual animal-based approach, leveraging longitudinal fecal sample collection over a 15-year period. We hypothesized that: 1) fecal microbiotas from the endangered SRKW population would differ from those of the relatively fit NRKWs and ARKWs and that this comparison could reveal population health biomarkers; and 2) that SRKW fecal microbiotas would demonstrate relative stability, allowing for the characterization of baseline structure in wild individuals over time and the identification of aberrant trends. To provide additional context for interpreting changes in microbial community composition, we also characterized temporal changes in the fecal microbiotas of aquarium-housed KWs. Finally, we linked fine-scale individual life history data and health status for individual SRKWs to their distal gut microbial community dynamics; in particular, we examined the functional potential of taxa detected in individuals immediately preceding death using metagenomics. This large-scale sampling effort offers important insights into the potential use of fecal microbiota structure as the basis for SRKW health surveillance.

## Methods

### Wild killer whale sample collection

We collected fecal samples from surface waters in the wake of killer whales (KWs) from three conspecific resident populations: Southern (SRKW), Northern (NRKW) and Alaska (ARKW) (**Fig. 1a, b**). Samples were collected during both dedicated and opportunistic field efforts between 2005 and 2019, using previously described field methods^11,37,39^. Opportunistically, samples were also collected from transient KWs (TKWs). Sample collection was conducted from small research vessels using long-handled pool nets or tricorner beakers and transferred immediately to single use sample bags or vials. Fecal samples were collected in U.S. waters (NMFS General Authorization No. 781-1725 and NMFS Scientific Research Permits 781-1824, 16163 and 21348) and Canadian waters (Marine Mammal License numbers MML 2006-02/SARA-24, 2007-03/SARA-64, 2008-03/SARA-84 and 2012-03 SARA-84, license numbers XMMS 8 2014 and 8 2014-Amendment 1, and file numbers 2014–22 SARA-355 and 2014–22 SARA-355). Sample collection methods were approved by the NWFSC/AFSC Institutional Animal Care and Use Committee (protocols A/NW2014-1 and A/NW 2015-2). Nearby surface seawater was collected concurrently with fecal samples during sampling of the SRKW and NRK populations in 2018 and 2019 using sterile 50mL conical vials. See **Supplementary Information** for more details.

**Fig. 1.**
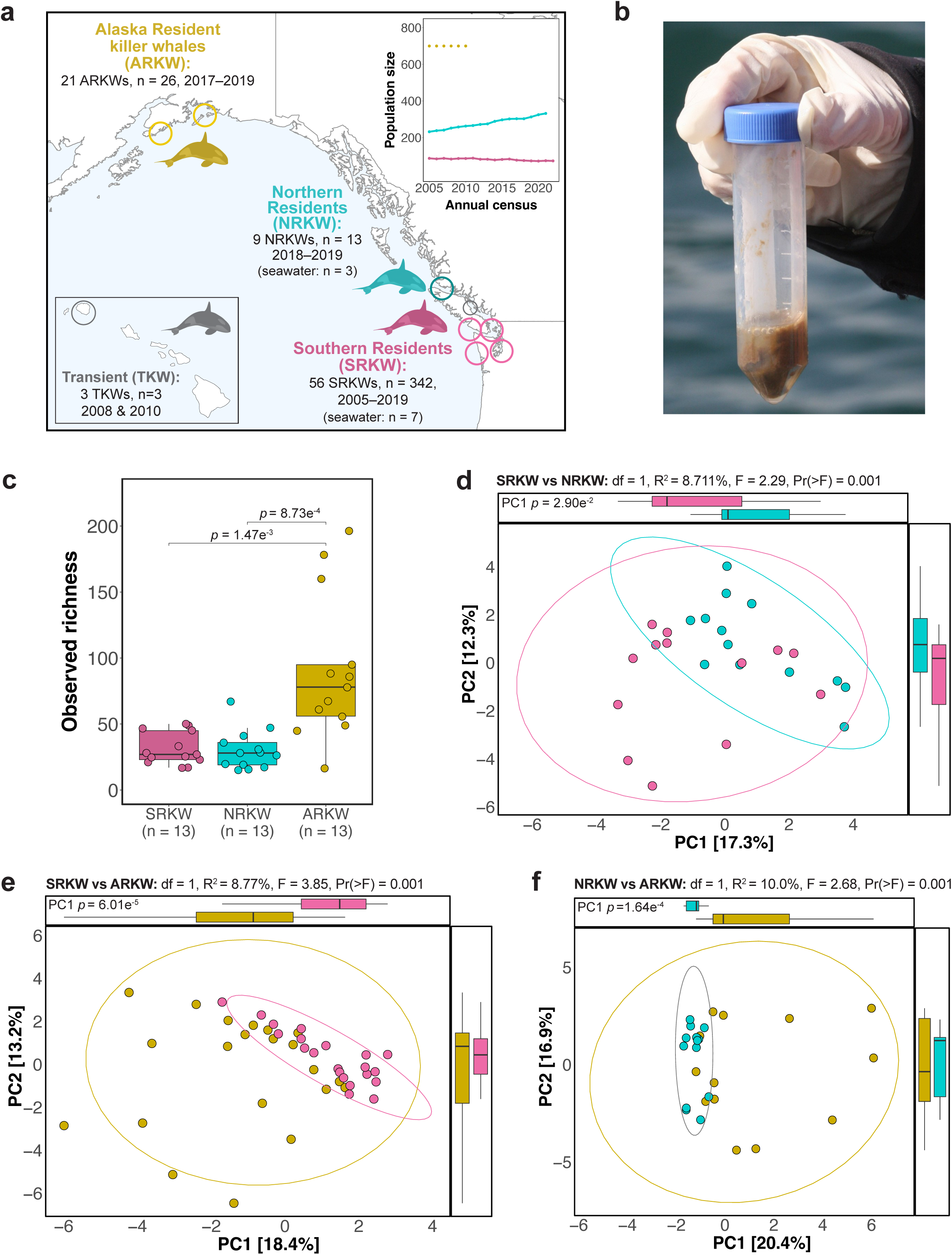
North Pacific Resident killer whale populations have distinct fecal microbiotas. (**a**) Map of the wild killer whale (KW) sampling locations. The inset graph shows annual population growth in the Northern Residents (NRKW; DFO 2020 census) versus the endangered Southern Residents (SRKW; https://www.mmc.gov) as solid lines. The dotted line shows the estimated population size (2005-2010) of the Alaska Residents (ARKW; Matkin *et al*.^10^). **(b)** Photograph of a KW fecal sample collected remotely from the seawater surface. **(c)** Boxplot comparison of the observed richness in each population’s fecal microbiotas. The dataset was restricted to fecal samples collected from 2017-2019, normalized (≤13 samples/population), and rarefied (39,824 reads/sample) prior to calculation of richness. Annotations denote significant differences identified by a pairwise Dunn test with Benjamini-Hochberg p-value adjustment. **(d-f)** Unconstrained principal component (PC) analysis ordinations of Aitchison distances between the (**d**) SRKW versus NRKW (13 each), (**e**) SRKW versus ARKW (21 each) and (**f**) NRKW versus ARKW (13 each) fecal microbiotas from 2017-2019. Titles denote the multivariate analysis of variance (’adonis2’) significance levels and R^2^ values (the variation explained by “Population”). Margins show univariable distribution boxplots along each PC; significant differences were identified via Dunn tests. Points, ellipses (95% CIs) and boxplots are colored by population.

### Aquarium-housed killer whale sample collection

Fecal samples were collected from 12 aquarium-housed KWs (AqKWs) by animal care staff as supervised by onsite veterinarians at SeaWorld facilities in the U.S. from April-July 2019. Samples were collected 3 times per week for the first week of each month, then once weekly. Fecal samples were collected using routine husbandry rectal catheter protocols. Samples of pool water were collected concurrently.

### Genomic DNA extraction and individual whale assignment

Fecal and water samples were stored frozen at -80°C without preservative until processing^40^. Total genomic DNA was extracted from fecal samples using a Qiagen QiaCube robot and the QIAamp DNA Stool Mini Kit or QIAamp Fast Stool DNA Mini kit (Qiagen, Valencia, CA) in a polymerase chain reaction (PCR)-free laboratory. Fecal sample aliquots were transferred into 2 mL vials for lysis as described previously^41^, including negative extraction controls with each set of extractions. Each fecal sample was genotyped at either a suite of 26 microsatellite loci^42^ or 68 single nucleotide polymorphism (SNP) loci^43^. Host identity was assigned with reference to existing genotypes from individually identified KWs, allowing stratification of data by age, sex and individual.

### 16S rRNA gene amplicon sequencing

The hypervariable V4 region of the 16S rRNA gene was amplified by PCR using barcoded primers 515 forward (5′-GTGYCAGCMGCCGCGGTAA-3′) and 806 reverse (5′-GGACTACNVGGGTWTCTAAT-3′). The 25 μL PCR reactions were carried out in triplicate in 96-well plates using 0.4 μM concentrations of each commercially synthesized primer, 2.5x Hot MasterMix (5 PRIME) and 3 μL of each DNA template. As a negative amplification control, at least one PCR reaction per 96 well plate was also run without DNA template. Thermal cycling conditions were as follows: 94°C for 3 min, followed by 30 cycles of 94°C for 45s, 50°C for 60s, and 72°C for 90s with a final extension step of 72°C for 10 min. Triplicates were pooled, purified and DNA was quantified as previously described^27^. Quantified DNA from all samples (including extraction and amplification controls) was included in two sequencing pools. The final DNA concentrations for purified amplicon pools were 51.90 ng/uL (Pool 1) and 74.23 ng/uL (Pool 2). Average amplicon sizes were 378 bp (Pool 1) and 396 bp (Pool 2). Library pools were sequenced (paired-end, 2×250 nucleotides) on separate lanes of an Illumina HiSeq 2500 in the same run (Rapid Run mode) with a 20% phiX spike-in at the Genetic Resources Core Facility at Johns Hopkins Genomics (Baltimore, MD).

### Amplicon sequence variant inference and taxonomic assignment

Demultiplexed reads were processed using the open-source package DADA2 and the parameters recommended in the “Big Data: Paired-end” workflow in R (see **Supplementary Code 1, Supplementary Code 2** and **Supplementary Data 1** for software versions, and citations). Read quality was assessed for each lane individually. Sequences from both lanes were merged prior to chimera removal. Taxonomy was assigned using a nailve Bayesian classifier (‘assignTaxonomy’, DADA2) and the SILVA v138 reference database. DADA2 was used to make species-level assignments for amplicon sequence variants (ASVs) with exact matches to sequences in the reference database (‘addSpecies’). A rooted phylogenetic tree was created in QIIME2 using the SILVA reference alignment.

### Quality filtering

Subsequent quality filtering and statistical analyses were performed using phyloseq (ps; **Supplementary Code 2**). Eukaryotic, chloroplast and mitochondrial reads and ASVs with unclassified phylum-level taxonomy were removed, leaving 10,480 unique ASVs. Examination of extraction and amplification controls identified sequences in 31/42 controls (median = 207.5 reads/control, range = 0–7750) representing 699 ASVs (median = 22 reads/ASV, range = 1–3269). Of these, 254 ASVs (2.42% of bacterial ASVs detected) were identified as contaminants via decontam (‘method = ‘prevalence’) and were pruned from the ps object. Samples with small library size (<4000 reads) were removed.

For alpha diversity analyses, permutation filtering (PERFect) was used to identify and remove potentially spurious ASVs while preserving richness. This yielded the “ps_alpha” object with 9,813 ASVs (72,639,633 reads) in 671 samples. For all other analyses, only ASVs present in ≥2 samples with ≥100 reads/sample were retained to emphasize biologically relevant taxa and to reduce computation time. This yielded “ps_filt” with 780 ASVs (71,869,905 reads) in 669 samples.

### Overall 16S rRNA gene amplicon sequence dataset

The wild KW dataset (considering ps_alpha) consisted of 384 fecal samples (resident: n=381, transient: n=3) with 6,460 ASVs (**Supplementary Code 2**), and 48,700,808 reads (mean = 126,825 ± 58,969 reads per sample), and 10 seawater samples with 863 ASVs and 575,578 reads (mean = 57,555 ± 17,636). The AqKW dataset consisted of 275 fecal samples with 3,309 ASVs and 23,249,677 reads (mean = 84,544 ± 25,241), and two pool water samples with 1,028 ASVs and 113,601 reads (mean = 56,800 ± 31,484).

### Statistical analyses and data normalization

Microbial community profiles were analyzed using phyloseq (**Supplementary Code 2**). Normalization was performed by random subsampling (dplyr) to control for sample size or repeated measures. Rarefaction was performed for analyses sensitive to sequencing depth (richness, dominance, turnover and Unifrac), otherwise raw read counts were used. Counts were center log ratio-transformed (clr) for Aitchison-distance based analyses.

### Alpha diversity

Alpha diversity analyses were performed using both Shannon index and observed species counts to capture both richness and evenness. After examining comparison group normality and variance (stats), Dunn tests or t-tests were performed with Benjamini-Hochberg (BH) p-value adjustment (ggpubr). Dominance was examined by identifying the most abundant ASV in each sample and calculating the relative frequency at which each ASV was dominant across the dataset (microbiomeutilities).

### Beta diversity and analysis of variance

Aitchison dissimilarity, Bray-Curtis, weighted and unweighted Unifrac (UWU) distance metrics were used for unsupervised data analyses. Aitchison dissimilarity, designed for compositional data and more robust to subsetting^44^, revealed the clearest partitioning of the data and was used moving forward. However, the main analyses were repeated using UWU to evaluate whether Aitchison results were primarily driven by abundance. Aitchison distance matrices were visualized using an unconstrained principal component analysis. The relative contribution of study covariates to community variation was determined using permutational MANOVA (’adonis2’, vegan) with ID in the permutation block (nperm=999) to account for repeated measures. Group dispersions and homogeneity were assessed using vegan.

### Differential abundance analyses

Taxa with differential abundance between KW groups were identified using the MaAsLin2 package. Workflow parameters were tailored to the structure of each dataset. Each linear model was run with BH p-value adjustment. To ensure robust interpretation of the results, three additional methods were utilized: treeDA, ALDEx2, and coda4microbiome. Indicator importance was determined based on the consensus across DA methods and effect size.

### Longitudinal diversity analyses

Diversity measures were regressed against time or age using an autoregressive general mixed effect linear model fit by restricted maximum likelihood with ‘ID’ treated as a random effect (nlme); marginal R^2^ values are shown. Turnover of microbiota composition between consecutive time points (t_1_, t_2_…) was calculated using the codyn package. Changes in mean turnover were regressed via linear models (stats) to identify non-zero temporal trends. SplinectomeR identified non-zero temporal diversity trends and significant trajectory differences between groups. To identify temporal indicator taxa, TITAN2 was performed to detect community-level change points over time. TITAN2 detects changes in taxonomic abundance, occurrence, and direction along a temporal gradient. For population-level analyses, TITAN2 was also performed using age as the temporal gradient to distinguish age-versus time-related trends.

### Taxonomic volatility and host survival outcome

Longitudinal taxonomic relative abundance (RA) was examined in the microbiotas of the most frequently sampled SRKWs (n ≥ 9) to identify “volatile” (temporally unstable with regards to baseline abundance) taxa in individuals with different health outcomes. For each SRKW, an individualized baseline of ASV abundance was created using all but their final two samples. For each ASV, RA was regressed against time to calculate a conservative baseline (95% prediction interval * 2.5) expected to capture all new observations and identify RA outliers that were indicative of taxonomic volatility. ASV RA in the final two samples was then examined to identify “blooming” taxa with RA that exceeded their baseline’s upper limit. Volatility was also examined with TITAN2 and QIIME2.

### Metagenomic sequence-based analysis

Metagenomic sequence data were generated from fecal samples that 1) had sufficient remaining DNA, 2) were collected from individuals within 6 months of death, and 3) had ASVs of interest based on the results of a preliminary 16S rRNA sequence-based analysis. Libraries were prepared for metagenomic sequencing using the Nextera DNA Flex Library Prep kit (Illumina 20018705) and sequenced to a target depth of 10Gbp per sample on one lane of an Illumina NovaSeq S4 at the DNA Services Lab, Roy J. Carver Biotechnology Center, University of Illinois at Urbana-Champaign.

Sequencing reads were filtered for host contamination (bbduk) with the Oorc_1.1 *Orcinus orca* reference genome (GCA_000331955.2). Each set was randomly down-sampled to 50 million read pairs (seqtk) and shuffled (fq2fa). Illumina adapters were removed (BBTools) and sequencing reads were trimmed (Sickle). Quality-filtered reads were then assembled using IDBA-UD. For each sample, reads were mapped back to the assembled scaffolds (bowtie2) to compute coverage values. Genes were predicted using Prodigal and predicted proteins were annotated using USEARCH and the KEGG, UniRef, and UniProt databases. Genome binning was performed both manually (ggkbase.berkeley.edu) and using automated bin generation (bowtie2, MetaBAT2). Bins were then reconciled with DAS Tool to create the best merged set. Preliminary taxonomic classifications were assigned using GTDB-Tk. Quality metrics were calculated manually and estimated using CheckM. Bins with ≥25% completeness were de-replicated (dRep) at 99% average nucleotide identity (ANI) for manual curation. All quality-filtered bins were visualized in Anvi’o and refined by removing sets of scaffolds with aberrant coverage profiles. Refined bins were then re-assessed for quality as described above.

Two metagenomically-assembled *Fusobacterium* genomes (MAGs) were selected for analysis. ANI was calculated (using the Proksee online interface) between both MAGs and 14 NCBI *Fusobacterium* reference genomes. MAGs were annotated using Proksee. A custom *Fusobacterium* virulence factor (VF) database was constructed using the relevant literature^45^ and potential VFs detected in each MAG were analyzed against the BLASTN and FusoPortal^46^ databases to identify the closest gene homologues.

### Data Availability

The data supporting the findings of this study are available within **Supplementary Tables 1 –11**, **Supplementary Data 1-9** (**1**, software citations; **2**, additional data about KWs and sampling; **3-7,** ASV tables; **8,** BLASTN results; **9**, virulence factor analysis), and two supplementary folders: “**QIIME2_Analysis**” and “**R_Files**” (**Supplementary Code 1**, raw data processing; **Supplementary Code S2**, data analysis; ps.rds; and sequences.xlsx). A link to the code and data repository will be made accessible to the public upon publication. 16S rRNA gene amplicon and metagenomic sequences are available at NCBI SRA BioProject ID PRJNA1370166.

## Results and Discussion

### Wild killer whale 16S rRNA sequence dataset

We examined taxonomic composition in the fecal microbiotas of 56 SRKW individuals (**Supplementary Table 1**) sampled from 08/12/2005 through 08/09/2019 (**Fig. 1a, b**), representing approximately 77% of the living population^5^. Of these, 42 SRKW individuals were sampled longitudinally over a duration ranging from 5 days to 13 years (**Supplementary Fig. 1**; n = 328, mean samples per individual = 8, range = 2-17). A total of 342 SRKW fecal samples were analyzed. After amplicon sequencing and quality filtering, 5,655 unique ASVs (mean = 85 ± 69 ASVs per sample) were identified in the SRKW dataset (considering ‘ps_alpha’ in **Supplementary Code 2**; mean = 130,465 ± 60,664 reads per sample).

To provide context for interpreting SRKW fecal microbiota structure, 39 fecal samples were collected from the geographically adjacent and partially sympatric NRKW and ARKW populations from 05/30/2017 to 08/18/19 (mean = 1 sample per individual, range = 1-3 samples, duration = 9-364 days). The 13 samples collected from 9 NRKWs yielded 161 unique ASVs (mean = 31 ± 15 ASVs, 83,492 ± 13,826 reads). The 26 samples collected from 21 ARKWs yielded 1,133 ASVs (mean = 91 ± 93 ASVs, 96,669 ± 25,127 reads). Three opportunistic samples were collected from three marine mammal-eating transient KWs (TKWs); these yielded 303 ASVs (mean = 160,965 ± 36,836 reads). Combined, the wild KW dataset (n = 384 from 89 individuals) yielded 6,460 unique ASVs (**Supplementary Data 2-7**).

To examine temporal microbiota stability in KWs under relatively controlled conditions, we characterized the fecal microbiotas of 12 aquarium-housed KWs (AqKWs) at two U.S. aquariums (**Supplementary Fig. 2**) over a four-month period. Individuals were clinically healthy upon study enrollment and did not receive antibiotics during the study period. A total of 275 fecal samples were collected (3309 unique ASVs; mean = 66 ± 48 ASVs, 84,544 ± 25,241 reads; **Supplementary Data 7**).

To address the potential for bacterial contamination, seawater was collected adjacent to SRKW and NRKW fecal samples during sample collection in 2018 and 2019 and analyzed in parallel. For the AqKWs, one pool water sample was collected at each facility (1028 ASVs; mean = 56,800 ± 31,484 reads). Consistent with previous studies demonstrating clear dissimilarities between marine mammal microbiotas and those of their aquatic environments^27,30,47,48^, KW fecal microbiota composition was distinct from that of their paired seawater samples (R^2^ = 55.0%, p <0.001) and no overlap was observed between the most abundant ASVs in each sample type (**Supplementary Fig. 3a-d**). The AqKW fecal microbiotas were also distinct from those of their pool water (**Supplementary Fig. 3e, f**). A more detailed discussion of the results is provided in **Supplementary Information**.

### *Paeniclostridium* ASV1 was detected in all wild killer whale fecal samples

*Paeniclostridium* ASV1 dominated the wild KW fecal microbiotas, regardless of ecotype or population (SRKW, 31.9% mean RA; NRKW 46.0%; ARKW 29.2%; TKW 41.1%) (**Supplementary Fig. 4a-d**), and had 100% prevalence among the wild KWs in this study (**Supplementary Fig. 4e, f; Supplementary Table 2**). This ASV was identical to the corresponding 16S rRNA gene region of *Paeniclostridium sordellii* (**Supplementary Data 8,** BLASTN results), a recognized pathobiont of humans and animals^49^. However, a previous study of the healthy pilot whale fecal microbiota^31^ found *P. sordellii* at similar abundance (RA=2.2%) as reported here for the AqKWs (3.4%, **Supplementary Fig. 4f**), which suggests it may act as a gut commensal. In terrestrial carnivores^50,51^ and raptors^52^, a high abundance of *P. sordellii* has been correlated with high dietary protein intake^53^. *P. sordellii* may enhance host immunity by secreting antimicrobials in the gut and support digestion by transforming bile acids^54^. In the present study, the ubiquity of ASV1 in the wild KWs was striking since it spanned two ecotypes with lifestyle differences (the piscivorous residents and the marine mammal-eating transients), suggesting an intimate association with these hosts^55^.

To date, the only information about the TKW microbiome comes from drone-captured respiratory samples^36^. Here, we provide, to our knowledge, the first characterization of the TKW fecal microbiota, albeit based on few samples (**Supplementary Fig. 5a-c**, **Supplementary Text**).

### The resident killer whale fecal microbiota was population-specific

The wild resident KW populations (SRKW, NRKW, and ARKW) share similar diet, morphology, and behavior; yet their fecal microbiotas displayed significant differences in alpha and beta diversity (**Fig. 1c-f**, **Supplementary 6a-d**). Analyses of between-population Aitchison dissimilarity revealed that each population had a distinct fecal microbial community structure (p = 0.001). The ARKW population had the richest microbiotas (SRKW vs ARKW, p = 1.47e^-3^; NRKW vs ARKW, p = 8.73e^-4^), while the NRKW population had the least diverse microbiotas (**Supplementary Table 3**). In humans, low gut diversity has been associated with disease^56,57^, although the literature suggests that the relationship between richness and health is more nuanced and variable^58–60^. A recent study found no correlation between dolphin health status and fecal microbial richness^61^. Here, overall richness did not seem to indicate population health; temporal demographic trends have shown the NRKW population to be arguably ‘healthier’ (i.e., demonstrating higher survivorship over decades) than the SRKW population^12^. The NRKW microbiotas had the highest frequency of samples dominated by universal taxon *Paeniclostridium* ASV1 (NRKW 63%, SRKW 41% and ARKW 42%; **Supplementary Table 3**). The dominance of core microbiota may be equally or more beneficial to KW intestinal ecology and host health than overall gut diversity.

### The SRKW fecal microbiotas were distinguished by taxa with pathogenic potential

Compared to the relatively fit resident populations, the microbiotas of the endangered SRKW were enriched in several genera known to include veterinary pathogens: *Fusobacterium*, *Photobacterium*, and *Edwardsiella* (**Fig. 2a**). The SRKW microbiotas had a mean RA of *Fusobacterium* = 24.5% (versus RA_NRKW_ = 0.16%, p <0.001 and RA_ARKW_ = 1.68%, p = 0.02), *Photobacterium* = 2.80% (versus RA_NRKW_ = 0.11%, p <0.001 and RA_ARKW_ = 0.09%, p = 0.003), and *Edwardsiella* = 8.31% (versus RA_NRKW_ = 0.89%, p <0.026 and RA_ARKW_ = 1.57%, p = 0.022).

**Fig. 2.**
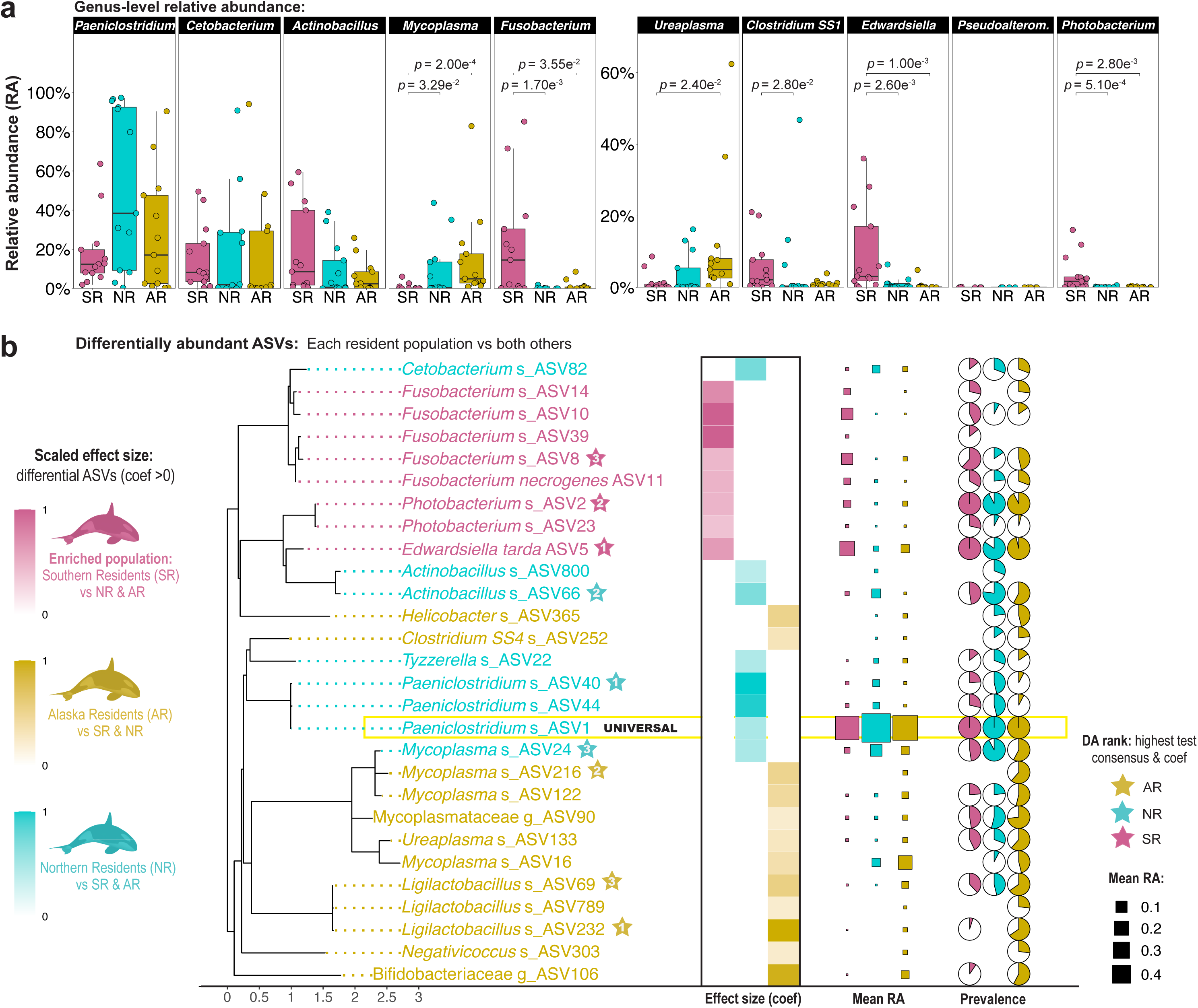
Southern resident fecal microbiotas had higher relative abundances of *Fusobacterium*, *Photobacterium* and *Edwardsiella* compared to those of relatively fit resident populations. Differential abundance (DA) between the fecal microbiotas of Southern (19 SRKWs, 22 samples), Northern (9 NRKWs, 13 samples) and Alaska (21 ARKWs, 26 samples) Resident killer whale populations (2017-2019). **(a)** Genus-level DA. Each dataset was normalized (≤3 samples/individual; 13 samples/population) and relative abundance (RA) boxplots are shown for the 10 most abundant genera in the combined dataset. Comparisons between populations were based on pairwise Dunn tests with Benjamini-Hochberg p-value adjustment. **(b)** Amplicon sequence variant (ASV) DA. Left: MaAsLin2 identified ASVs enriched in each population’s fecal microbiotas versus those of both others (**Supplementary Fig. 7**). ASVs with ≥10% prevalence and ≥0.1% RA in ≥1 population, MaAsLin2 coef >1.5, and p.adj <0.05 (correction=BH) are shown in a phylogenetic tree. ASV color indicates the enriched population. Heatmap gradients indicate MaAsLin2 coefs scaled to the overall maximum. Taxa enriched in both fit populations relative to the SRKWs (e.g., ASV252) are colored by the population with the highest RA. Stars denote robust indicator taxa identified by ≥2 additional DA tests (**Supplementary Table 4**) and DA rank (1 = highest coef). Right: Sizes of squares indicate ASV RA. Pie charts denote ASV prevalence. “UNIVERSAL” denotes the only ASV with 100% prevalence in the wild KW dataset (**Supplementary Fig. 4f**).

Commensal fusobacteria have been identified in the guts of a variety of terrestrial^88^ and marine mammals, including dolphins^27^, pinnipeds^47^, and sea otters^48^. *Fusobacterium* ASV8 differentiated the SRKWs from the other resident KWs (rank_DA_= 3, **Fig. 2b**, **Supplementary Fig. 7**, **Supplementary Table 4**) and most closely resembled sequences of isolates from healthy dogs (Per.ID=99.6%). Still, certain fusobacteria are opportunistic pathogens that can cause inflammation and tissue necrosis^62^. In humans, *Fusobacterium* has been implicated in chronic intestinal diseases such as inflammatory bowel disease and colorectal cancer (CRC)^63^. Overgrowth of *Fusobacterium* in the gut has been associated with intestinal inflammation, malabsorption^64^ and malnutrition in elderly adults^65^. In the SRKW population, the role of *Fusobacterium* is unclear— the other four differentially abundant ASVs matched sequences from rectal swabs of healthy, managed dolphins^27^ but ASV11 and ASV39 also matched *Fusobacterium mortiferum*, which has been implicated in pancreatitis^66^ and CRC^67^ in humans. Of note, ASV39 was undetected in the fecal microbiotas of either relatively fit resident population.

*Photobacterium* has previously been identified in the guts of wild dolphins^68^ and salmon^69^. Certain strains have been associated with weakened gut barrier function and increased disease susceptibility in salmon^70^. In humans, *Photobacterium* can cause tissue necrosis and septicemia^71^. Here, the ASV that distinguished the SRKWs from the other resident KWs (ASV2, rank_DA_= 2) was identical to *Photobacterium damselae*, a marine generalist and fish pathogen^72^. This taxon is prevalent in pinnipeds foraging near salmon net farms, yet it is undetermined whether *P. damselae* represents a gut commensal or a commensal pathogen^73^. Its role in cetaceans is also unclear; *P. damselae* has been documented in the guts of healthy managed pilot whales^31^ and dolphins^74^, but has also been implicated in cetacean strandings based on post-mortem findings^75^. Here, *P. damselae* was more abundant in the SRKWs (RA = 1.83%) and less abundant in the NRKWs (RA = 0.10%) and ARKWs (RA = 0.69%). However, *P. damselae* had similarly high prevalence in all three resident populations (92.31–100.00%), which could indicate a commensal association with the wild KW host or shared environmental exposures.

*Edwardsiella* was primarily represented by *Edwardsiella tarda* ASV5 (all other ASVs had RA<0.15% in each of the KW subgroups), which was the most robust indicator for the SRKW fecal microbiotas (rank_DA_= 1). *Edwardsiella tarda* is a zoonotic pathogen of fish, birds, reptiles and mammals^76^. High prevalence has been described in the blow and feces of wild bottlenose dolphins^74^, but *E. tarda* has also been implicated in cetacean strandings^77^. A meta-analysis of infectious disease threats in the SRKW population suggested there was minimal risk of an *E. tarda* epizootic due to its low transmissibility in other mammalian hosts^78^. However, *E. tarda* was identified as the cause of death in a SRKW individual^79^, which suggests it has the capability to act as an opportunistic pathogen.

### Northern and Alaska Resident killer whale fecal microbiotas were enriched for mycoplasmas and lactic acid bacteria

The fecal microbiotas of both relatively fit resident KW populations were characterized by their differential RA of *Mycoplasma* ASVs. The NRKW microbiotas had the highest RA (5.44%, **Supplementary Table 4**) and prevalence (92.31%) of *Mycoplasma* ASV24 (rank_DA_= 3, **Fig. 2b**), which is related to Mycoplasma-like taxa in the stomach of healthy cows (Per.ID=98.7%). Low-abundance *Mycoplasma* ASV216 (rank_DA_= 2, RA = 0.51%, prevalence = 61.5%), was specific to the ARKW microbiotas and had no close match (>95.7%) in the NCBI nucleotide database.

The genus *Mycoplasma* includes known pathogens and species that have been isolated from dead marine mammals^80^, as well as from the feces of healthy dolphins^81^ and the blow of humpback whales^82^. Mycoplasmas are highly host-adapted^83^. A serosurvey across 23 marine mammal species identified distinct cetacean- and pinniped-associated *Mycoplasma* clades and demonstrated phylosymbiosis^84^. Some mycoplasmas dominate the intestinal tract of Chinook salmon^85^, the preferred prey of all three resident KW populations. There is evidence for co-evolution and mutualism between salmon and certain mycoplasmas; *Mycoplasma salmoninae* can utilize ammonia byproducts to biosynthesize vitamins, detoxifying the gut in the process^83^. *Mycoplasma* RA correlates with salmon body condition, prompting its development as a population health biomarker^70,86^.

The ARKW microbiotas had the highest diversity of lactic acid-producing bacteria (LAB, **Supplementary Table 5**). All 10 LAB ASVs were most similar (Per.ID=98.7-99.6%) to sequences previously recovered from healthy managed dolphins living in seawater^27,87^. Three LAB were unclassified Bifidobacteriaceae. Bifidobacteriaceae ASV106 was differentially abundant in the ARKW microbiotas (RA_mean_ = 2.57%, prevalence = 57.7%) compared to both other resident populations (RA_mean_ <0.01%). Seven LAB belonged to *Ligilactobacillus*, a genus that previously comprised the *Lactobacillus salivarius* group^88^. *Ligilactobacillus* ASV232 (RA_mean_= 0.66%, prevalence = 65.38%) was the most differential taxon in the ARKW microbiotas (rank_DA_= 1). The only LAB detected in the SRKWs (RA = 0.15%) and NRKWs (RA = 0.01%), *Ligilactobacillus* ASV69, was prevalent across the resident populations (prevalence: SR = 36.36%, NR = 46.15%, AR = 65.38%) but most abundant in the ARKW microbiotas (RA = 1.75%).

LAB promote host health by enhancing epithelial integrity and modulating immunity^89^ and have been extensively studied as probiotics^90^. *Bifidobacterium* species are gut commensals of terrestrial and marine mammals that include probiotic strains^91^. *Ligilactobacillus* species are highly host-adapted; at least 18 different species have been isolated from a wide range of vertebrates, including cetaceans^88^. Probiotic use of *Ligilactobacillus salivarius* has been studied in several host species, revealing anti-inflammatory^92^ and antibacterial^93^ benefits. Orally administered *L. salivarius* has been shown to improve gut mucosal health and weight gain in chicken^94^, mice^92^ and pigs^95^. Interestingly, *Ligilactobacillus* ASV69 (the only LAB found in all resident KW populations) shared an identical amplified 16S rRNA gene sequence to antimicrobial *L. salivarius* strains that we previously isolated for the development of a marine mammal probiotic^87^.

ARKW may harbor more LAB because this resident population is farther removed from the relatively polluted waters of the Salish Sea and Puget Sound^96^. In mice, exposure to organochloride pesticides reduced abundance of *Ligilactobacillus*, increasing vulnerability to opportunistic infection^97^. PCB exposure has also been associated with lower levels of *Bifidobacterium* and *Lactobacillus* in the mouse gut^98^. The relatively low PCB burden of the ARKW population^99^ could promote LAB acquisition or persistence. Other anthropogenic threats associated with urbanized waterways could also influence LAB abundance and diversity. A population-wide depletion of LAB was observed in a longitudinal study of wild meerkats as environmental stressors intensified^100^. In mice, *L. salivarius* abundance is correlated with survivorship and weight^92^. Similarly, LAB may benefit ARKW health and survivorship.

### A shared common core fecal microbiota in wild resident killer whale populations

Taxa described as “core” for a specific population can be consistently identified in their microbiotas to a degree that implies a functionally important host-association^101^. Having already identified “universal wild KW taxon” *Paeniclostridium* ASV1 in every wild KW fecal sample, we identified other core taxa (prevalence ≥ 50% and RA ≥ 1.0%) for the resident KWs. In the 2017–2019 dataset, all three populations shared five additional core ASVs that comprised the “resident KW core”: *Cetobacterium* ASV6 (RA = 10.63%, **Supplementary Table 2**), *Actinobacillus delphinicola* ASV3 (7.96%), *Clostridium SS1* ASV7 (3.74%), *Cetobacterium* ASV20 (3.45%) and *Ureaplasma* ASV17 (2.41%). The four most abundant were identical to gastric sequences from healthy managed dolphins^27^ and ASV17 matched fecal sequences from free-swimming wild porpoises (**Supplementary Data 8**). Only one core taxon, *Fusobacterium* ASV8 was SRKW-specific (RA = 5.41%) and rare (RA<1.0%) in the other resident populations. This specificity could reflect the differences in population health status, environmental exposures, or signify an emerging pathogen. Several resident KW universal/core taxa were also members of the AqKW core fecal microbiota: *Paeniclostridium* ASV1 (RA = 3.37%, prevalence = 91.64%), *Cetobacterium* ASV6 (RA = 3.07%, prevalence = 88.36%) and *Clostridium SS1* ASV7 (RA = 1.19%, prevalence = 93.4%).

Previous studies have described *Cetobacterium* as one of the most dominant taxa in the fecal microbiotas of wild dolphins^68^ and porpoises^102^ and in SRKW respiratory mucus^36^. Here, ASV6 was identical to *Cetobacterium ceti* previously isolated from dolphins that demonstrated the ability to biosynthesize vitamin B_12_ – an essential cofactor for myo-and hemoglobin production^103^. Marine mammals cannot biosynthesize vitamin B_12_, so microbial production may be integral for producing the high concentrations of myo- and hemoglobin required for prolonged underwater activity. *Clostridium SS1* ASV7 sequences were identical to those of *Clostridium perfringens.* Despite its pathogenic potential^49^, *C. perfringens* has been described as an abundant distal gut microbe in healthy dolphins^27^, pilot whales^31^, and pinnipeds^47^, indicating a commensal role in these fish-eating hosts. *C. perfringens* is also common in the guts of wild terrestrial carnivores^104^, and its abundance may correlate with protein intake^53^. Here, our finding that *C. ceti* and *C. perfringens* were core taxa for the fish-eating (residents and AqKWs) but not marine mammal-eating (TKWs) subgroups, suggests an association with a piscivorous diet.

While the AqKW core fecal microbiota overlapped with that of the wild KWs, differences were observed. Although *Paeniclostridium* ASV1 was a core taxon for the AqKW microbiotas (**Supplementary Table 2**), *Romboutsia* ASV4 was dominant (RA = 26.69%) and “universal” (prevalence = 100.0%). This taxon and five others were unique to the AqKW core fecal microbiota. Overall, the AqKW fecal microbiotas were compositionally distinct from those of the wild residents (Pr(>F) <0.001, R^2^ = 12.4%), but these differences were largely driven by phylogenetically related taxa (**Supplementary Figs. 8**, **9**). This suggests that while environmental conditions may select for different taxa, some degree of functional redundancy may persist to maintain homeostasis. Here, because our primary motivation for studying AqKWs was to characterize microbial temporal variation, husbandry-related analyses are provided in the **Supplementary Text**.

### Temporal signatures of individuality and stability in the wild Southern Resident and aquarium-housed killer whale fecal microbiotas

To identify longitudinal trends in the wild SRKW population, it was important to establish whether the KW fecal microbiome exhibits individuality and relative temporal stability to support the feasibility of discriminating normal versus abnormal variation. Temporal microbial stability has previously been studied in managed dolphins^27–29^, belugas^30^ and pilot whales^31^; these gut communities were found to be individualized and relatively stable over several months. We predicted that fecal microbiotas of KW individuals would similarly demonstrate short-term stability contributing to a population-level baseline. Because sampling of wild KWs is both seasonal and opportunistic, we first sought to identify and characterize a temporal baseline for KWs of known health status living in a controlled environment.

### Killer whale distal gut microbiotas exhibited individualized structure and relative temporal stability for up to one year

Host individuality accounted for the greatest contribution to microbiota variation in the AqKW dataset (adonis2: Pr(>F) = 0.001, R^2^ = 33.86%) (**Fig. 3a**, **Supplementary Fig. 10**, **Supplementary Table 6**), followed by facility (Pr(>F) = 0.001, R^2^ = 5.41%), sex (Pr(>F) = 0.001, R^2^ = 4.71%), study month (Pr(>F) = 0.001, R^2^ = 1.02%) and age (Pr(>F) = 0.001, R^2^ = 0.44%). Considering only adult AqKWs (9 AqKWs, 206 samples), mean between-sample dissimilarity (Aitchison distance) was plotted against the time interval between sample collection (7–105 days) (**Fig. 3b**). This showed a significant effect of time (trendyspliner: p = 0.001) in which samples collected farther apart in time were slightly more dissimilar. However, this slope was visually subtle (time lag = 0.03 days), so significance was likely inflated due to pseudo-replication (pairwise comparisons = 12,636). No significant temporal trends in richness (p = 0.167) or Shannon index (p = 0.668) were observed in these AqKWs (**Supplementary Fig. 11a**, **b**). The time lag analysis was repeated for intra-individual and inter-individual comparisons separately (**Fig. 3c**) which revealed that samples collected from the same AqKW were more compositionally alike than were those from different AqKWs. This indication of individuality persisted throughout the 4-month AqKW study period (slidingspliner: p-value range:1.09e^-05^–7.92e^-05^, time lag = 7–105 days).

**Fig. 3.**
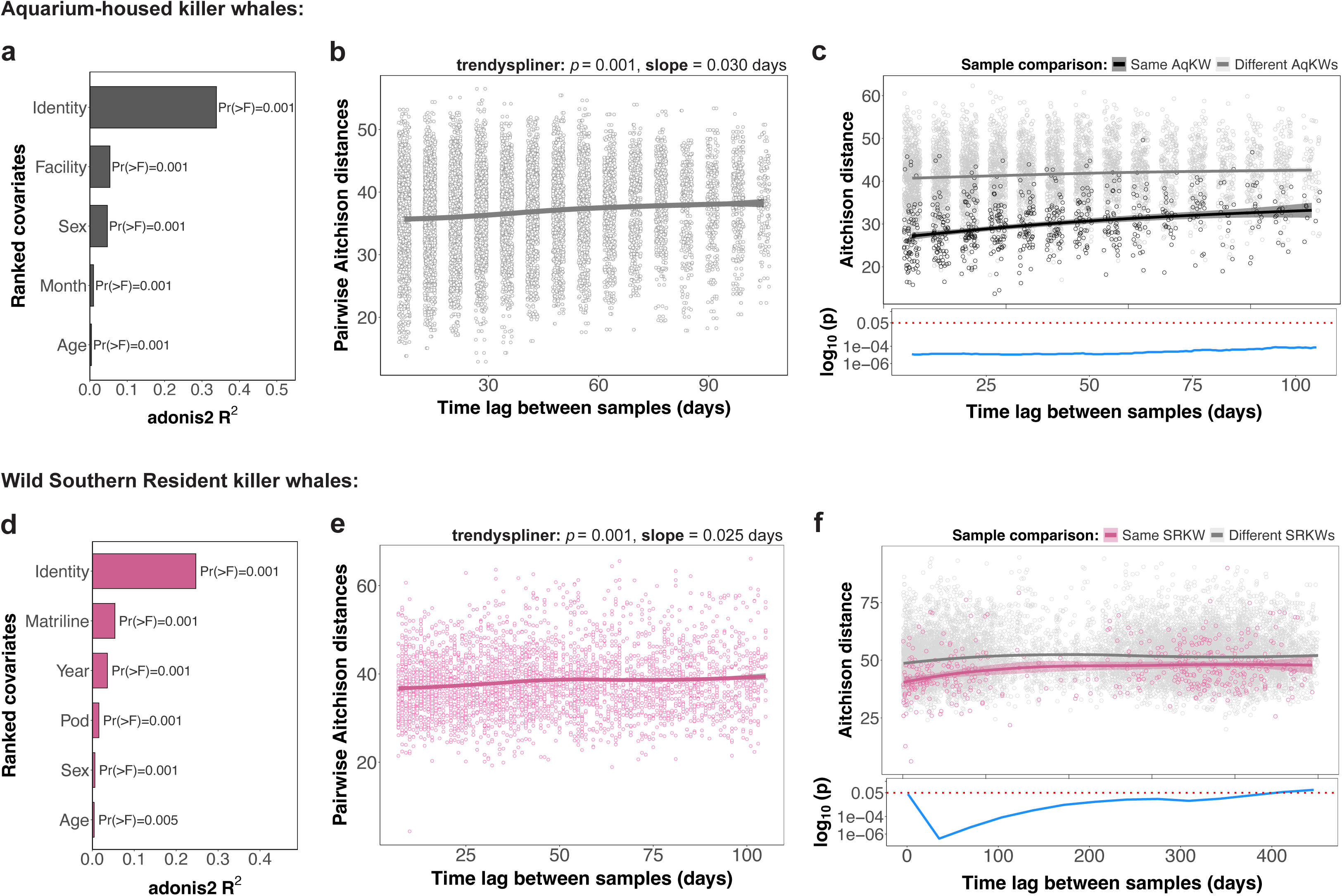
Killer whale fecal microbiotas exhibited temporal stability and individualized structure over a four-month period. (**a-c**) Aquarium-housed killer whale (AqKW) fecal samples (collected 2019). (**a**) Multivariate analysis of variance (MANOVA, ‘adonis2’) of clr-transformed Aitchison distances among the AqKW fecal microbiotas. The bar chart shows covariates included in the model ranked by the proportion of variance explained (R^2^, **Supplementary Table 6**). Data were normalized (n=22 per AqKW). (**b**) Time lag graph of Aitchison distances showing pairwise dissimilarities among weekly fecal samples from the nine adults (>9 years old, n=206 samples) over the study period (time lag=7-105 days). Lines denote Loess smoothing (dark gray=95% confidence interval). Temporal trends were identified using a longitudinal permutation test (‘trendyspliner’, splinectomeR). (**c**) Top: Time lag graph of Aitchison distances showing pairwise dissimilarities among fecal samples (from **b**) collected from the same versus different AqKWs. Lines show Loess smoothing colored by comparison group. Bottom: Longitudinal p-values (blue line) denoting temporal windows of significant differences between group trajectories (‘slidingspliner’). Red line indicates the significance threshold (p=0.05). (**d-e**) Wild Southern Resident killer whale (SRKW) fecal samples (collected 2005–2019). (**d**) MANOVA results (as in **a**) for all SRKWs with confirmed identity (53 SRKWs, n=338). (**e**) Time lag graph (as in **b**) of fecal samples collected 7-105 days apart (44 adult SRKWs, n=291) to match the AqKW time lag range in **b**. (**f**) Time lag graph (as in **c**) of all adult SRKW fecal samples (46 SRKWs, n=299).

The SRKW longitudinal dataset covered a considerably longer, 15-year timeframe, yet SRKW identity also had the greatest influence on variation (adonis2: Pr(>F) = 0.001, R^2^ = 24.17%) (**Fig. 3d, Supplementary Fig. 12a-e**), followed by matriline (Pr(>F) = 0.001, R^2^ = 5.39%), year (Pr(>F) = 0.001, R^2^ = 3.61%), pod (Pr(>F) = 0.001, R^2^ = 1.58%), sex (Pr(>F) = 0.001, R^2^ = 0.62%) and age (Pr(>F) = 0.005, R^2^ = 0.46%). Considering only adult SRKWs (46 SRKWs, 299 samples), between-sample dissimilarity also showed a subtle increase (slope = 0.025 days, trendyspliner = 0.001, 7400 pairwise comparisons) over longer time periods (7–105 days apart), suggesting relative stability over this time frame (**Fig. 3e**). The SRKW dataset was restricted to June–Sept (the 4-month period with the largest number of samples) and no significant trends in richness (p = 0.120–0.991) or Shannon index (p = 0.08–0.735) were identified (**Supplementary Fig. 11c**, **d**). The time lag analysis of intra-versus inter-individual samples found that those collected from the same SRKW individual were more alike in composition than those from different SRKWs (slidingspliner: p<0.05) for a similar 105-day period as examined in the AqKWs. When all available SRKW fecal samples were considered, this difference was significantly different using a time interval ≤377 days (p-value range: 3.05e^-07^-0.036, **Fig. 3f**). Aitchison distances between fecal samples from the six J pod SRKW individuals with ≥10 longitudinal samples showed an influence of time along PC1 (adonis2: Pr(>F) = 0.001, R^2^ = 8.3%) and ID along PC2 (Pr(>F) = 0.001, R^2^ = 13.3%) and each individual’s trajectory along PC2 was relatively stable over several years (trendyspliner p=0.103-0.978, **Supplementary Fig. 12f**).

These results were consistent with other studies of managed cetaceans (dolphins and belugas) that previously demonstrated the presence of individualized distal gut microbiotas that were temporally stable over time periods from 6 weeks^29,30^ to 6 months^27^. To our knowledge, this is the first study to characterize temporal stability in the gut microbiota of a wild cetacean population. Establishing a temporal baseline within the SRKW population is an important first step in identifying unexpected variation in the microbiota potentially associated with disease and emerging threats.

### Over 15 years, taxa became undetected, and richness declined in the Southern Resident killer whale population

The size of the SRKW population declined from 89 to 75 individuals over the 15-year study period^8^, so the full longitudinal dataset (2005–2019) was analyzed to identify temporal trends that may have reflected or contributed to a decline in population health. Ordination of between-sample Aitchison dissimilarity between SRKW fecal samples (ps_filt, n = 340) showed that date of sample collection accounted for separation of the data along the primary axis (**Fig. 4a, b**), indicating change in microbiota composition with time (p_day_ = 8.0e^-3^; UWU p_day_ = 4.0e^-3^, **Supplementary Fig. 13a**, **b**). To assess the degree of confounding with host age, the analysis was repeated using age as the independent variable (**Supplementary Fig. 13c**, **d**), which revealed a subtle but significant temporal trend (p_age_ = 0.021). However, examination of the KW age distribution showed that 91.5% of samples belonged to SRKWs ≤58 years old and all samples from older SRKWs (73–102 years old) belonged to three age-outliers (J2, K11 and J8; total of 29 samples). Repeating the analysis excluding those samples (**Supplementary Fig. 13e**, **f**) showed a significant time-related trend (p_day_ = 3.0e^-3^) and the age-related trend was no longer observed (p_age_ = 0.07).

**Fig. 4.**
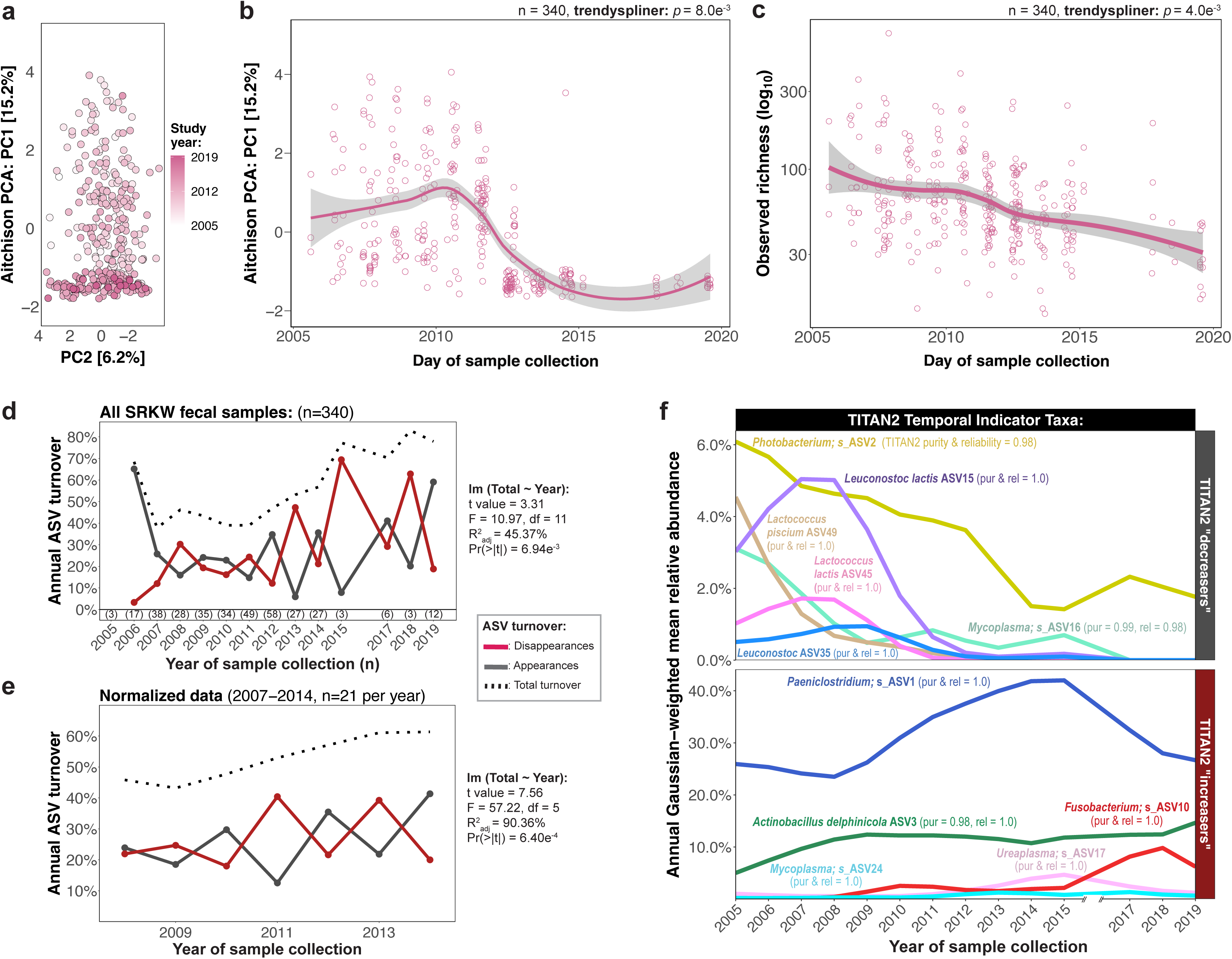
Richness in the Southern Resident killer whale fecal microbiotas declined over time as certain taxa became undetectable and others bloomed. (**a**) Unconstrained principal component analysis ordination of clr-transformed Aitchison distances among all available Southern Resident killer whale (SRKW) fecal samples (n=340, points colored by year). (**b**) Principal component 1 loadings plotted against day of sample collection. (**c**) Richness of rarified data (23,712 reads/sample) plotted against day of sample collection. For **b** and **c**, lines show Loess smoothing (confidence intervals=95%). Significance denotes non-zero temporal trends identified by ‘trendyspliner’ (splinectomeR). (**d**) Turnover of microbiota membership against year of sample collection (codyn) using all samples (annual n shown in parentheses). Solid lines represent the proportion of ASV appearances (gray) and disappearances (red) between consecutive years (t_1_, t_2_…). The dotted black line shows total ASV turnover (ASVs_Gained_ t_2_ + ASVs_Lost_ t_2_/Richness t_1_ + t_2_). A linear regression model was used to identify temporal trends. (**e**) Turnover of microbiota membership using normalized (2007–2014, n=21 per year) and rarified data (28,729 reads/sample). Only samples collected June-Sept were used to control for season. (**f**) Specific ASV contributions to temporal community change. Longitudinal mean annual relative abundance (RA) for ASVs with RA >0.40% identified as temporal indicator taxa by TITAN2 (purity and reliability scores ≥99.6%, **Supplementary Figs. 15-17**, **Supplementary Table 7**). Facets indicate TITAN2 ASV classifications (“increasers” or “decreasers”) based on RA trajectory. Lines denote the Gaussian-weighted RA averaged over consecutive 2-year windows.

Richness in the SRKW fecal microbiotas declined over time (p_day_ = 4.0e^-3^; **Fig. 4c**) independent of host age (p_age_ = 0.24, **Supplementary Fig. 13g**). Community turnover of ASVs significantly increased (Pr(>|t|) = 6.94e-^3^) throughout the study period (**Fig. 4d**). The mean rate of ASV appearances increased 3.95% between the first (2005-2011: 28.08%) and second half (2012-2019: 29.19%) of the study (**Supplementary Code 2**), while ASV disappearances increased 112.19% (from 7.55% to 37.42%). Turnover analysis of a normalized dataset (2007-2014, 21 samples/year, 28,729 reads/sample) also showed a significant increase in temporal ASV turnover (Pr(>|t|) = 6.40e-^4^, **Fig. 4e**). Overall, the 40.78% increase in ASV disappearances (2007-2010 = 21.53%, 2011-2014 = 30.31%) exceeded the 15.55% increase in ASV appearances (24.05% to 27.79%). No significant temporal (p_day_ = 0.10) or age-related (p_age_ = 0.37) trends in Shannon diversity (**Supplementary Fig. 13h**, **i**) were found to correspond with declining richness, which implied the loss of low-abundance taxa. Rare ASVs (RA<2.0%) declined over time (**Supplementary Fig. 14a,** lm: t = -4.39, Pr(>|t|)_rare_ = 8.85e^-4^), but only one out of the seven abundant ASVs (all resident KW core ASVs) had decreasing RA (*Photobacterium* ASV2: TITAN2 purity & reliability = 0.98) (**Supplementary Fig. 15a**, **b**; **Supplementary Table 7**). The two most abundant taxa, *Paeniclostridium* ASV1 (pur = 0.996, rel = 1.0) and *A. delphincola* ASV3 (pur = 0.978, rel = 0.996) became more abundant and dominant over time (mean relative frequency: 2005–2011 = 45.6%; 2012–2019 = 66.6%) (**Supplementary Fig. 16a**, **b**). All seven abundant ASVs met core criteria throughout the 15-year study duration (**Supplementary Table 8**)— this “temporal core” of abundant taxa likely bolstered community evenness as rare taxa were lost from the SRKW microbiotas over time.

A temporal core typically includes abundant taxa (that are less prone to local extinction), but temporally stable low-abundance taxa may also be essential^105^. Low-abundance commensals play biologically important roles in immune modulation, microbiome stabilization and resilience^106^. Two lower-abundance LAB classified as early core taxa, *Leuconostoc lactis* ASV15 and *Leuconostoc* ASV35 (**Fig.** 4f, **Supplementary Fig. 15**), became undetected in the SRKW microbiotas over time. ASV15 dominated 2.9-13.2% of SRKW fecal microbiotas from 2007–2010 (**Supplementary Fig. 14b**) but declined and were undetected after 2014 (**Supplementary Table 8**). The decline of both *Leuconostoc* ASVs also correlated with age, but similar trends were observed among juveniles, 10-25-year-old adults, and older adults, implying a true effect of time (**Supplementary Fig. 17**). *Leuconostoc lactis* has previously been shown to have immunostimulatory properties with probiotic potential^107^. Strains isolated from guts of marine fish has been shown to inhibit pathogen growth^108^. *Leuconostoc* ASV35 had an identical 16S rRNA gene sequence to strains from healthy farmed salmon^109^. Another early core LAB, *Lactococcus lactis* ASV45, also declined to negligible abundance (RA<0.01%) over time. *Lactococcus lactis* has been shown to reduce inflammation in the gut^110^ and promote longevity in model organisms^111^. TITAN2 identified a total of seven ASVs that became undetected-including another rare LAB, *Lactococcus piscium* ASV49 (**Supplementary Fig. 15**). Fish-associated *L. piscium* isolates have demonstrated antimicrobial properties that prevent the spoilage of salmon meat^112^. Given this probiotic potential, the loss of core LAB may contribute to pathology within the SRKW population.

Resident KW core taxon *Ureaplasma* ASV17 was less abundant in the early SRKW microbiotas but reached core threshold midway through the study (2012-2015, RA 2.93%, prevalence 76.52%) (**Supplementary Table 8**, **Supplementary Fig. 16**). This corresponded with a decline in *Ureaplasma* ASV46 (**Supplementary Fig. 15**), which was shared with the relatively fit NRKWs (RA 0.38%) and ARKWs (RA 0.03%). A similar pattern was observed with *Mycoplasma* ASV24 (core for NRKW and ARKW) which gradually increased over time to become abundant (2012-2015, RA = 1.21%, prevalence = 49.57%) circa 2014 when the declining *Mycoplasma* ASV16 became undetected in the SRKW. ASV16 was abundant in both the NRKW (RA 2.80%, prevalence 7.69%) and ARKW (RA 8.06%, prevalence 46.15%) populations (**Supplementary Table 2**). These trends imply that certain ASVs may become abundant to counteract the loss of phylogenetically related ASVs, and that functional redundancy may exist within the SRKW microbiotas to prevent the loss of essential traits^106^.

### A transient bloom of a detoxifying bacterium in the Southern resident killer whale fecal microbiotas

A mid-study “bloom” of *Comamonas testosteroni* ASV73 was observed in the SRKW microbiotas (**Supplementary Fig. 16**). Initially, ASV73 was undetected (2005–2006 and 2011) or rare (2007-2010, mean RA ≤0.13%, n≤3 annually) (**Supplementary Code 2**) but it abruptly became abundant in 2012 (RA = 1.55%, 19 individuals, 33 samples). ASV73 then declined (2013-2019, RA ≤0.01%, n≤1 annually), becoming undetected after 2015. This bloom could indicate a population-wide perturbance or novel exposure— the Seattle sewer system overflowed in 2012, spilling 55 million gallons of waste into the Puget Sound^113^. *Comamonas testosteroni* is commonly identified in polluted environments and capable of degrading sewage-associated contaminants^114^, so ASV73 may have imparted detoxifying functionality to the SRKW microbiotas. Interestingly, *C. testosteroni* can also degrade PCBs^114^. Since SRKWs have a high PCB burden^20^, *C. testosteroni* may have had a competitive advantage in colonizing their guts.

### *Fusobacterium* ASV10 became a dominant taxon in later years

As mentioned above, taxa identified as TITAN2 “increasers” in the SRKW microbiotas could reflect new exposures or functional compensation. But since SRKW population health was declining over time, taxa that simultaneously became abundant could be indicators of sub-optimal host health or acts as pathogens. Emerging taxa that were rare in the other resident populations were of particular interest, given the SRKW’s lower fitness. *Fusobacterium* ASV10 was a temporal “increaser” in the SRKW microbiotas (TITAN2: pur = 1.0, rel = 0.996) (**Supplementary Table 7**) and had RA ≤0.01% in all other wild subgroups examined (**Supplementary Table 2**). ASV10 was detected every study year but initially had low abundance (**Supplementary Code 2**) (2006–2008, annual mean RA 0.1 ± 0.1%, prevalence 28.1 ± 1.6%) and was not dominant in any sample (**Supplementary Fig. 14b**). After 2008, *Fusobacterium* ASV10 became abundant (2009-2019, RA 1.9 ± 1.6%, prevalence 39.1 ± 5.4%, excluding years with ≤6 samples) and it dominated 3.7–8.8% of fecal samples. Interestingly, a decades-long study of wild meerkat fecal microbiotas found that worsening environmental stressors correlated with enrichment of *Fusobacterium* and higher population-wide mortality^100^. Here, the enrichment of *Fusobacterium* ASV10 in the SRKW microbiotas over time is correlated with, and may have contributed to, their declining survivorship.

### Analyzing the Southern Resident killer whale microbiotas in conjunction with survival data

Determining the health of wild cetaceans is challenging and the seasonal movement of SRKWs in response to prey abundance further limits sampling opportunities and visual health assessments in winter months (e.g., <8% of our samples were collected Nov-April). However, some health events can be inferred from strong social bonds: a missing pod member can be presumed dead, and a pregnant female can be presumed to have aborted or suffered a perinatal death if it is observed soon thereafter to be thin without a calf^11^.

A retrospective literature search and detailed population census records^5^ identified SRKW deaths and reproductive events^11^ during our study period (**Supplementary Fig. 18**). Six SRKW individuals were sampled within 6 months of disappearance and presumed death (classified as “Dying”, SRKW_DYING_) (10 samples). This 6 month cutoff was chosen because it approximated the mean survival time for SRKWs once observed to be emaciated (body condition = “BC1”)^115^. For comparison, data from SRKWs observed to be alive as of 2024 (“Survived”, 23 SRKWs, 115 samples) were normalized to the same timeframe as SRKW_DYING_ (SRKW_SURVIVED_: 9 SRKWs, 34 samples, collected 2008, 2010, 2012-2013, 2017). Reproductive events were identified in 11 SRKWs sampled during pregnancy (25 samples) and/or lactation (25 samples) (**Supplementary Text**).

### Decreased abundance of *Paeniclostridium* and increased abundance of *Fusobacterium* detected in dying Southern Resident killer whales

We compared the most abundant genera in the fecal microbiotas of SRKW_DYING_, SRKW_SURVIVED_, and the NRKW and ARKW populations (**Fig. 5a**, **Supplementary Fig. 19**). In accordance with earlier analyses, SRKWs (both survival groups) had higher abundance of *Fusobacterium* and *Edwardsiella*. NRKWs, ARKWs and SRKW_SURVIVED_ shared relatively high abundances of *Mycoplasma* (mean RA= 2.79-11.54%) and *Ureaplasma* (3.57-6.80%) but these genera were rare in SRKW_DYING_ (*Mycoplasma* RA 0.14%, *Ureaplasma* RA 0.82%) (**Supplementary Code 2**). SRKW_DYING_ had the lowest RA of *Paeniclostridium* and the highest RA of *Fusobacterium* and *Photobacterium*.

**Fig. 5.**
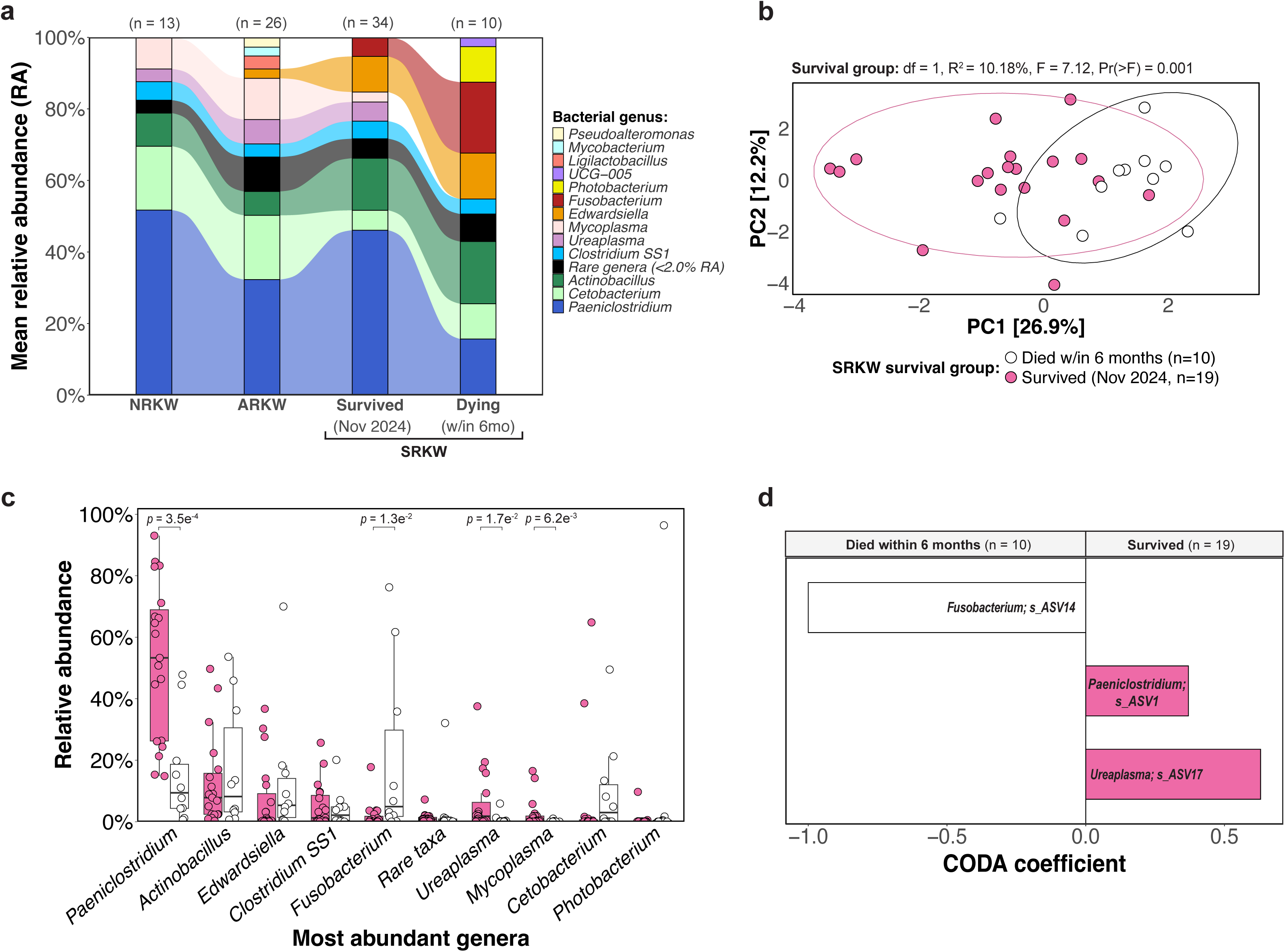
Southern Resident killer whales within 6 months of death had distinct fecal microbiotas from those of wild killer whales with better survival outcomes. (**a**) Alluvial plot comparing the relative abundance (RA) of genera with RA ≥2.0% in the fecal microbiotas of 9 Northern, 21 Alaska, and 44 Southern Resident killer whales. Two Southern Resident killer whales (SRKW) survival groups were examined: 6 SRKWs sampled within 6 months of death (“Dying”; n = 1-3 per individual) and 9 SRKWs that survived through 2024 (“Survived”; n = 1-10). SRKW_SURVIVED_ samples were selected from the same time year as SRKW_DYING_ samples (June-Oct 2008, 2010, 2012, 2013 & 2017). (**b, c)** For comparison of SRKW survival groups, data from **a** were normalized to n ≤ 3 per SRKW (SRKW_DYING_ n = 10, SRKW_SURVIVED_ n = 19). (**b**) Unconstrained principal component analysis of clr-transformed Aitchison distances among SRKW samples colored by “Survival_Group” with 95% data ellipsoids. The proportion of variance explained by “Survival_Group” (adonis2) relative to other study covariates (Fig. 3d) is shown above. (**c**) Boxplots showing the eight most abundant genera arranged by descending mean RA. Dunn tests were performed with Benjamini-Hochberg p-value adjustment. (**d**) Taxa with differential abundance between survival groups were identified using *coda-lasso* penalized linear regression on log-ratio transformed data (coda4microbiome, nfolds=6). Coefficient size is shown in the bar graph.

SRKW_SURVIVED_ were further restricted to samples collected during the same months (June-Oct, 19 samples) as SRKW_DYING_ for the direct comparison of the survival groups. Ordination of microbiota dissimilarity showed significant clustering by survival group along the primary axis (adonis2: Pr(>F) = 0.001, R^2^ = 10.18%) (**Supplementary Fig. 5b**). SRKW_SURVIVED_ had higher RA of *Paeniclostridium* (p = 3.5e^-4^), *Mycoplasma* (p = 0.017) and *Ureaplasma* (p = 0.006) whereas SRKW_DYING_ had higher RA of *Fusobacterium* (p = 0.013; **Fig. 5c**). *Ureaplasma* ASV17 and *Paeniclostridium* ASV1 were the most differentially abundant ASVs in the SRKW_SURVIVED_ fecal microbiotas and *Fusobacterium* ASV14 best discriminated SRKW_DYING_ (**Fig. 5d**). *Fusobacterium* ASV14 sequences were identical to rectal microbiota sequences from humans with colitis, but also to sequences from healthy pinnipeds, which suggests this taxon may act as an opportunistic pathogen. Overall, the differences seen between the SRKW survival groups mirrored those observed when the entire endangered SRKW population was compared to the relatively fit comparators (**Fig. 2**) and support the conclusion that *Paeniclostridium*, *Mycoplasma* and *Ureaplasma* may provide health benefits to the KW host. Interestingly, a study of wild porpoises during an infectious disease epidemic found that *Paeniclostridium* was more abundant in healthy versus diseased hosts^116^. These results also implicated *Fusobacterium* with disease, so we predicted *Fusobacterium* might exhibit temporal “volatility” (fluctuations of unexpected magnitude) prior to death.

### Volatile *Fusobacterium* abundance dynamics detected in fecal samples collected prior to death

Longitudinal samples were available from three “dying” SRKWs, J8 (15 samples), L2 (10 samples), and L26 (9 samples) (**Fig. 6a**). For each, taxonomic abundance in their early fecal microbiotas was used to create individualized baselines, with which to predict ASV RA over time and identify volatile taxa at later collection dates. This analysis identified 5 volatile ASVs in the final samples collected from J8 and L26, and 4 volatile ASVs from L2 (**Fig. 6b-d**). To determine whether taxonomic volatility was unique to SRKW_DYING_, we repeated these analyses using all other SRKWs with ≥10 samples (**Fig. 6e**). Three of these SRKWs (J17, K21 and J2) died during the study but their final sample was collected >6mo (1.2-3.3 years) from death (“died later”); a total of five volatile taxa were identified in these individuals. Considering both SRKW_DYING_ and SRKW_DIED.LATER_, *Fusobacterium* ASVs were the only volatile taxa to be identified in more than one individual and *Fusobacterium* ASV14 was the most frequently volatile. No volatile taxa were identified in SRKW_SURVIVED_ (individuals with ≥9 samples, J26, J31 and J27) using all analytical methods.

**Fig. 6.**
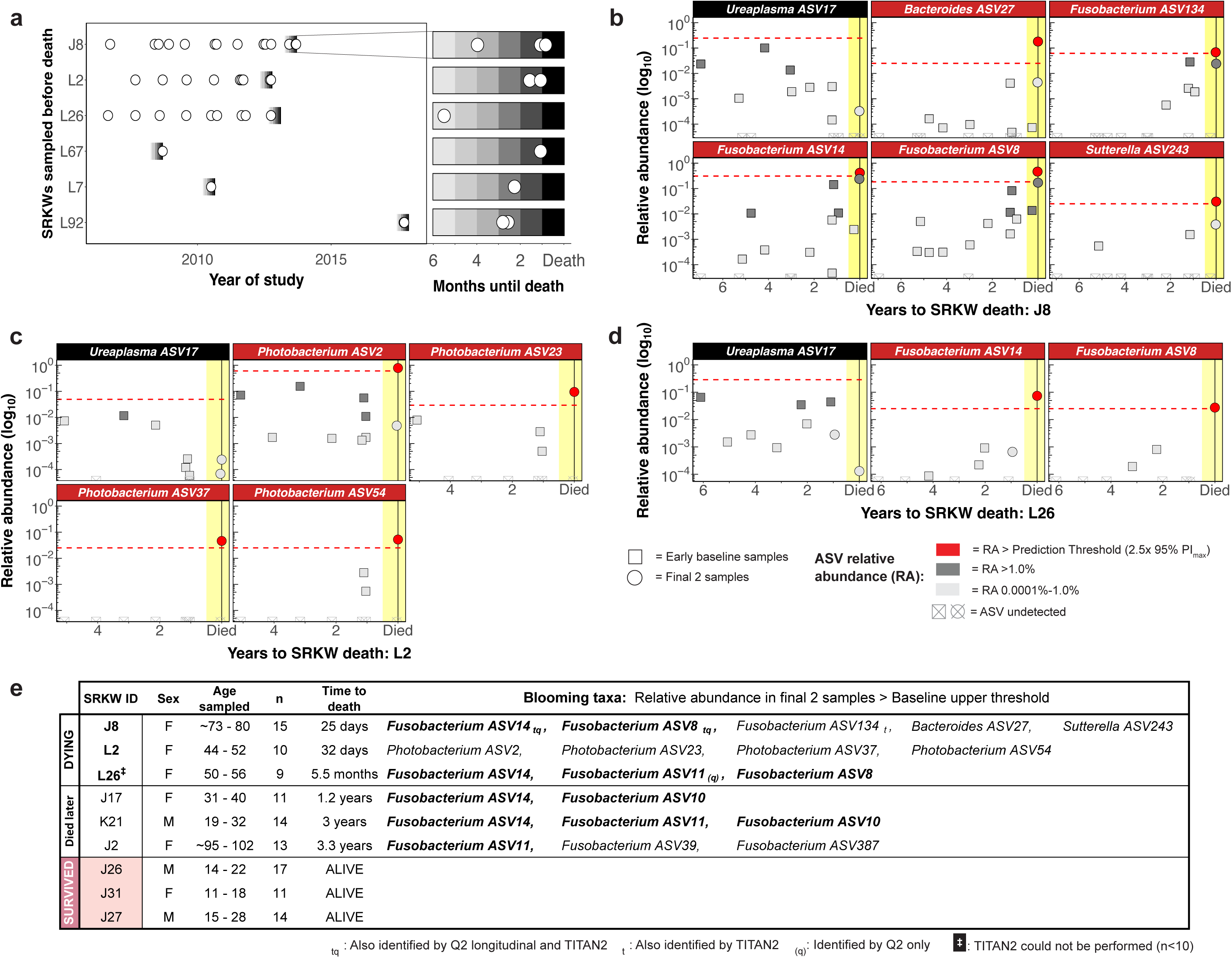
*Fusobacterium* and *Photobacterium* ASVs were identified as blooming taxa in final samples from dying Southern resident killer whales (SRKWs), but volatility was not detected in SRKWs that survived. (**a**) Timeline of samples collected from SRKWs within six months of their death (“Dying”). (**b-d).** Taxonomic volatility in dying SRKWs with ≥ 8 samples: J8 (**b**), L2 (**c**) and L26 (**d**). For each, the final two samples (circles) were excluded, and earlier samples (squares) were used in a linear regression model to predict ASV relative abundance (RA) over time. For each ASV, the upper limit of the 95% prediction interval (PI_max_) was used to determine “prediction threshold” (PT = 2.5 * PI_max_), indicated by the red line. ASVs with RA > PT in the final samples were considered “blooming” (red facets). Point fill denotes ASV RA: red = volatile (RA > PT), dark gray = abundant (RA >1.0%), light gray = rare (RA ≤1.0%), unfilled/crossed = undetected. Yellow shading denotes 6 months before death. For comparison, core taxon *Ureaplasma* ASV17 (no volatility detected) is also shown (black facets). (**e**) All SRKWs with frequent sampling (n > 8) were examined as in **b**-**d**. Volatile taxa identified in >1 SRKW are in bold. Two additional volatility analyses were performed: QIIME2 ‘feature-volatility’ and TITAN2. Subscripts denote volatile taxa identified by multiple methods.

Considering that more taxonomic volatility was identified in SRKW_DYING_ than in SRKW_DIED.LATER_ individuals, and volatility was not detected in SRKW_SURVIVED_, these results suggest that volatility itself is an indicator of ill health. Volatility may imply an unstable, less resilient gut microbiome that is more prone to adopting a state associated with ill health^105^. In humans and mice, volatility of the gut microbiota has been associated with chronic stress^117^ and gut inflammation^57^. Here, the most frequently volatile taxa in dying SRKWs were all affiliated with *Fusobacterium*. This is consistent with our earlier findings that *Fusobacterium* differentiated the endangered SRKWs versus healthier comparator populations, late versus early years in the declining SRKW population, and dying versus surviving SRKW health groups. In addition, *Fusobacterium* ASV11 was dominant among pregnant SRKWs with poor pregnancy outcomes (**Supplementary Fig. 20**, **Supplementary Table 9**), although the number of individuals with known pregnancy status was small and the outcome data were confounded by related factors.

We sought to better understand the role of *Fusobacterium* in SRKW health by examining their functional potential and recovered two *Fusobacterium* metagenome-assembled genomes from the final fecal sample collected from L26 (5.5 months prior to death). The *Fusobacterium* MAGs corresponded to distinct species (ANI = 72.44%) (**Supplementary Table 10**) and were also distinct from 14 NCBI *Fusobacterium* reference genomes (ANI = 69.16-78.85%). A total of 90 potential virulence factors (VFs) were identified in these two MAGs (**Table 1**, **Supplementary Table 11**, **Supplementary Data 9**), including 28 established VFs previously identified in other *Fusobacterium* species^45^ as involved in hemolytic activity (*hlyC/corC, tlyA, hlyD, shlB, tldD, septicolysin*), adherence (*ompH* and *upaG*), biofilm formation (*pfo, dnaJ*), and Type III (*ctpA*) and Type IV (*virB4* and *virB11*) secretion systems. The detection of numerous established VFs in both L26 *Fusobacterium* MAGs suggests a possible role of their host bacteria as pathogens in SRKW hosts. Further exploration of their activity within the SRKW gut using complementary methods (e.g., metabolomics and metatranscriptomics) could help elucidate their association with host health.

**Table 1.**
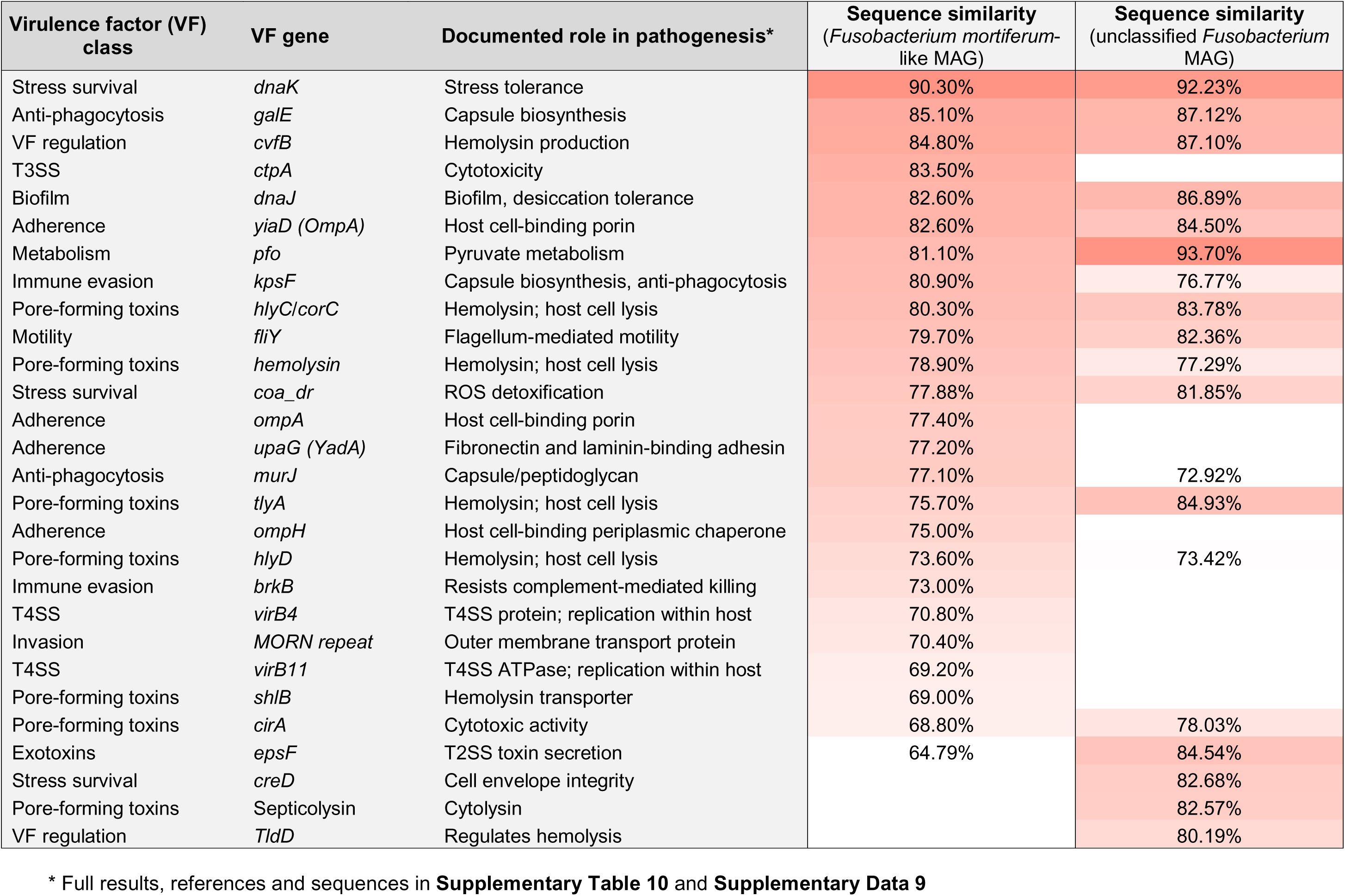
Putative virulence factor genes identified in the two novel *Fusobacterium* genomes recovered from the fecal microbiota of a dying SRKW. Two metagenomically-assembled genomes (MAGs) affiliated with *Fusobacterium* were recovered from the last fecal sample collected from SRKW L26 (5.5 months prior to death). The annotated MAGs were searched for putative virulence factor (VFs) genes homologous to those previously identified in *Fusobacterium necrophorum* using the FusoPortal database of *Fusobacterium* genomes^46^. VFs are listed in descending rank order based on sequence similarity score for the homologue found in each MAG. Darker shades of salmon highlighting indicate higher degrees of sequence similarity. Query sequences are provided in **Supplementary Data 9**. References and the results of a broader VF search are shown in **Supplementary Table 11**.

| Virulence factor (VF)<br>class | VF gene | Documented role in pathogenesis* | Sequence similarity<br>(*Fusobacterium mortiferum*-<br>like MAG) | Sequence similarity<br>(unclassified *Fusobacterium*<br>MAG) |
| --- | --- | --- | --- | --- |
| Stress survival | *dnaK* | Stress tolerance | 90.30% | 92.23% |
| Anti-phagocytosis | *galE* | Capsule biosynthesis | 85.10% | 87.12% |
| VF regulation | *cvfB* | Hemolysin production | 84.80% | 87.10% |
| T3SS | *ctpA* | Cytotoxicity | 83.50% |  |
| Biofilm | *dnaJ* | Biofilm, desiccation tolerance | 82.60% | 86.89% |
| Adherence | *yiaD (OmpA)* | Host cell-binding porin | 82.60% | 84.50% |
| Metabolism | *pfo* | Pyruvate metabolism | 81.10% | 93.70% |
| Immune evasion | *kpsF* | Capsule biosynthesis, anti-phagocytosis | 80.90% | 76.77% |
| Pore-forming toxins | *hlyC/corC* | Hemolysin; host cell lysis | 80.30% | 83.78% |
| Motility | *fliY* | Flagellum-mediated motility | 79.70% | 82.36% |
| Pore-forming toxins | *hemolysin* | Hemolysin; host cell lysis | 78.90% | 77.29% |
| Stress survival | *coa_dr* | ROS detoxification | 77.88% | 81.85% |
| Adherence | *ompA* | Host cell-binding porin | 77.40% |  |
| Adherence | *upaG (YadA)* | Fibronectin and laminin-binding adhesin | 77.20% |  |
| Anti-phagocytosis | *murJ* | Capsule/peptidoglycan | 77.10% | 72.92% |
| Pore-forming toxins | *tlyA* | Hemolysin; host cell lysis | 75.70% | 84.93% |
| Adherence | *ompH* | Host cell-binding periplasmic chaperone | 75.00% |  |
| Pore-forming toxins | *hlyD* | Hemolysin; host cell lysis | 73.60% | 73.42% |
| Immune evasion | *brkB* | Resists complement-mediated killing | 73.00% |  |
| T4SS | *virB4* | T4SS protein; replication within host | 70.80% |  |
| Invasion | *MORN repeat* | Outer membrane transport protein | 70.40% |  |
| T4SS | *virB11* | T4SS ATPase; replication within host | 69.20% |  |
| Pore-forming toxins | *shlB* | Hemolysin transporter | 69.00% |  |
| Pore-forming toxins | *cirA* | Cytotoxic activity | 68.80% | 78.03% |
| Exotoxins | *epsF* | T2SS toxin secretion | 64.79% | 84.54% |
| Stress survival | *creD* | Cell envelope integrity |  | 82.68% |
| Pore-forming toxins | Septicolysin | Cytolysin |  | 82.57% |
| VF regulation | *TldD* | Regulates hemolysis |  | 80.19% |

\* Full results, references and sequences in **Supplementary Table 10** and **Supplementary Data 9**

## Summary

Wildlife restoration efforts focused on environmental factors often neglect the host indigenous microbial ecosystem that is intertwined with health and vulnerable to disturbance by external factors. In association with a 15-year period of SRKW population decline, we illustrate the value of using fecal microbiota data to inform and support conservation practices, and present population-level microbiota comparisons, temporal analysis within the SRKW population, comparisons between SRKW survival groups, and changes within dying individuals. The SRKWs exhibited distinct fecal microbiotas from those of more fit resident populations, despite sharing a similar diet with both and an overlapping geographic range with NRKWs. Over time, SRKW fecal microbiota richness declined in tandem with host population size, and several rare taxa became undetected, including mycoplasmas that were likely associated with their piscivorous diet as well as LAB with gut-health promoting properties.

Remotely collected fecal samples provide a promising option for non-invasive SRKW health surveillance, yet the feasibility of detecting abnormal temporal trends necessitated establishing a baseline of normal variation. Under the controlled conditions of aquaria, we generated the first longitudinal description of baseline variation for this host species using clinically healthy representatives and found that the KW fecal microbiotas were temporally stable and individualized over four months. The larger wild SRKW population also revealed relative stability and individuality in samples collected <400 days apart, suggesting that SRKW fecal microbiotas might be harnessed to track individual health based on annual survey samples. Strategic bi-annual sampling in fall and spring (time lag ∼6 mo) could provide data from the SRKWs prior to their seasonal range shift to the outer coast and shortly after their return to the Salish Sea to help researchers infer health history over the winter, when the population is rarely observed.

We identified two taxonomic indicators associated with ill-health, *Fusobacterium* and *Photobacterium*, and both showed volatile abundance within SRKW individuals just prior to death. These taxa were prevalent at low abundance in other KW subgroups, suggesting that they might act as opportunistic pathogens within stressed populations or individuals. However, the lack of detailed health data for SRKW individuals, aside from survival outcomes, may have hindered our ability to identify other potential pathogens. Fortunately, advancements in aerial photogrammetry now allow for remote determination of SRKW body condition score - a quantifiable health metric that could better contextualize paired microbiome data in future studies.

This study provides the first large-scale, longitudinal characterization of the wild cetacean distal gut microbiome, and presents strong evidence that the endangered SRKW population’s established stressors (decreased prey availability, vessel disturbance, and environmental contaminants) influence the structure of their fecal microbiotas. This finding is consistent with previous studies of humans and mice^32–34^ that have shown malnutrition, stress, and toxins to alter gut communities, sometimes in manners that exacerbate or ameliorate the initial insult. Here, we argue that SRKW conservation efforts should adopt a holistic approach that considers interactions between the gut microbiome, the host, and the external environment, plus the looming risk of infectious disease. For instance, vessel disturbance can induce physiological stress among the SRKWs, leading to an altered microbiome and increased susceptibility to pathogens^35,118,119^. An altered microbiome could exacerbate the impact of prey scarcity if it reduced the ability of SRKWs to extract nutrients from their available prey^120^. Exposure to PCBs could select for detoxifying taxa or bio-activators that exacerbate toxicity^121^. This study emphasizes the interconnectedness between the host, its gut microbiome, and the environment and provides a complementary approach for understanding the complex pathophysiology that threatens the SRKW population.

## Acknowledgements

The authors gratefully acknowledge the support of the many staff, interns and volunteers who have supported killer whale fieldwork over the years, and the veterinary teams and trainers at SeaWorld San Diego and SeaWorld San Antonio for supporting the collection of fecal, prey and water samples. We thank Karen Steinman (SeaWorld and Busch Gardens Species Preservation Lab, SeaWorld California, CA) for her help with curating and sending samples. This is a SeaWorld Parks Technical Contribution, number 2025-6. We are grateful for the considerable efforts of the NOAA and DFO field crews who made possible the collection of wild killer whale fecal samples. We thank Robert Pesich and other members of the Relman Lab for coordinating the shipment, processing and storage of samples, as well as for their thoughtful input to this work.

Funding to support collection and processing of Southern Resident killer whale fecal samples was provided by the National Oceanic and Atmospheric Administration. Funding for collection of fecal samples from Alaska Resident killer whales was provided to the North Gulf Oceanic Society by the Exxon Valdez Trustee Council and the U.S. Marine Mammal Commission. This project was partially supported by Fisheries and Oceans Canada through Oceans Protection Plan/Marine Environmental Quality contribution agreement funding, and through grant MEQ-2023-24-009 to D.A.R.

## Author Contributions

A.D.S., K.M.P., L.P., D.A.R. conceptualized and designed the study; K.M.P., M.B.H., L.P., T.R., C.M., S.J.T. provided funding; K.M.P., M.B.H., L.P., D.A.R. managed project; K.M.P., M.B.H., C.E., L.P., T.R., K.H., S.O., C.M., S.W., S.J.T. provided samples; A.D.S., K.M.P., C.E., K.H., S.O., A.R., S.W., S.J.T. curated and/or managed data and samples; J.H., A.L.J., A.R., A.W., D.M. generated data; A.D.S., J.H., A.L.J., E.M.B., A.R., A.W., D.M. analyzed data; A.D.S. drafted manuscript; A.D.S., K.M.P., C.E., L.P., T.R., A.L.J., E.M.B., S.J.T., D.A.R. edited manuscript.

