## Supplementary Figures for "Fifteen-year microbiome survey of endangered killer whales (*Orcinus orca*) reveals declining diversity and population differences"

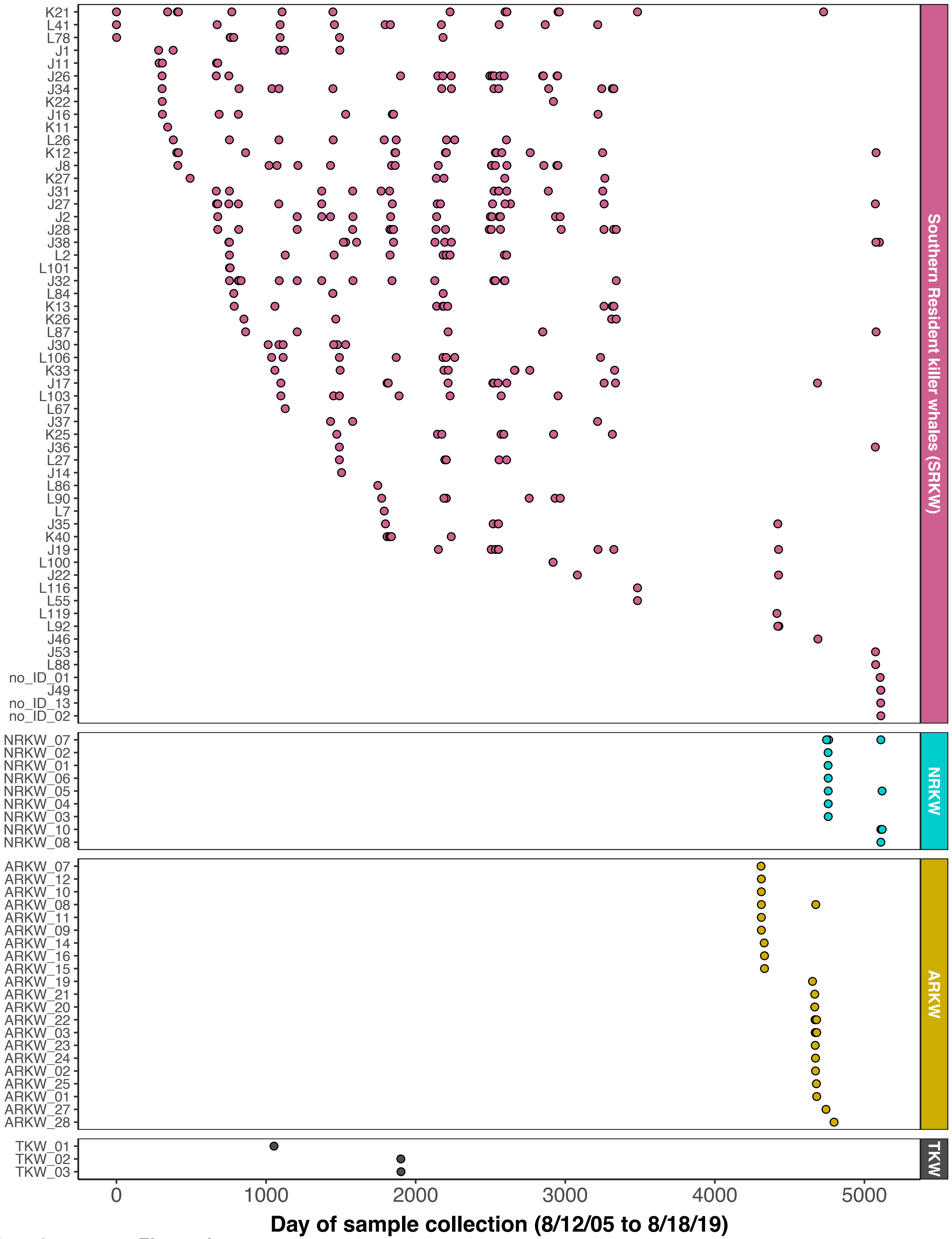


**Supplementary Figure 1**

**Supplementary Fig. 1. Sampling timeline for wild killer whales.** Timeline of wild killer whale (KW) fecal sample collection, faceted by ecotype (transient vs residents) and population (Southern, Northern and Alaska Residents). Individual identity was established for Southern Residents by genotyping host DNA in the feces using a reference database. Unidentified individuals represent young KWs whose DNA had not been cataloged. Reference databases were not available for the other groups.

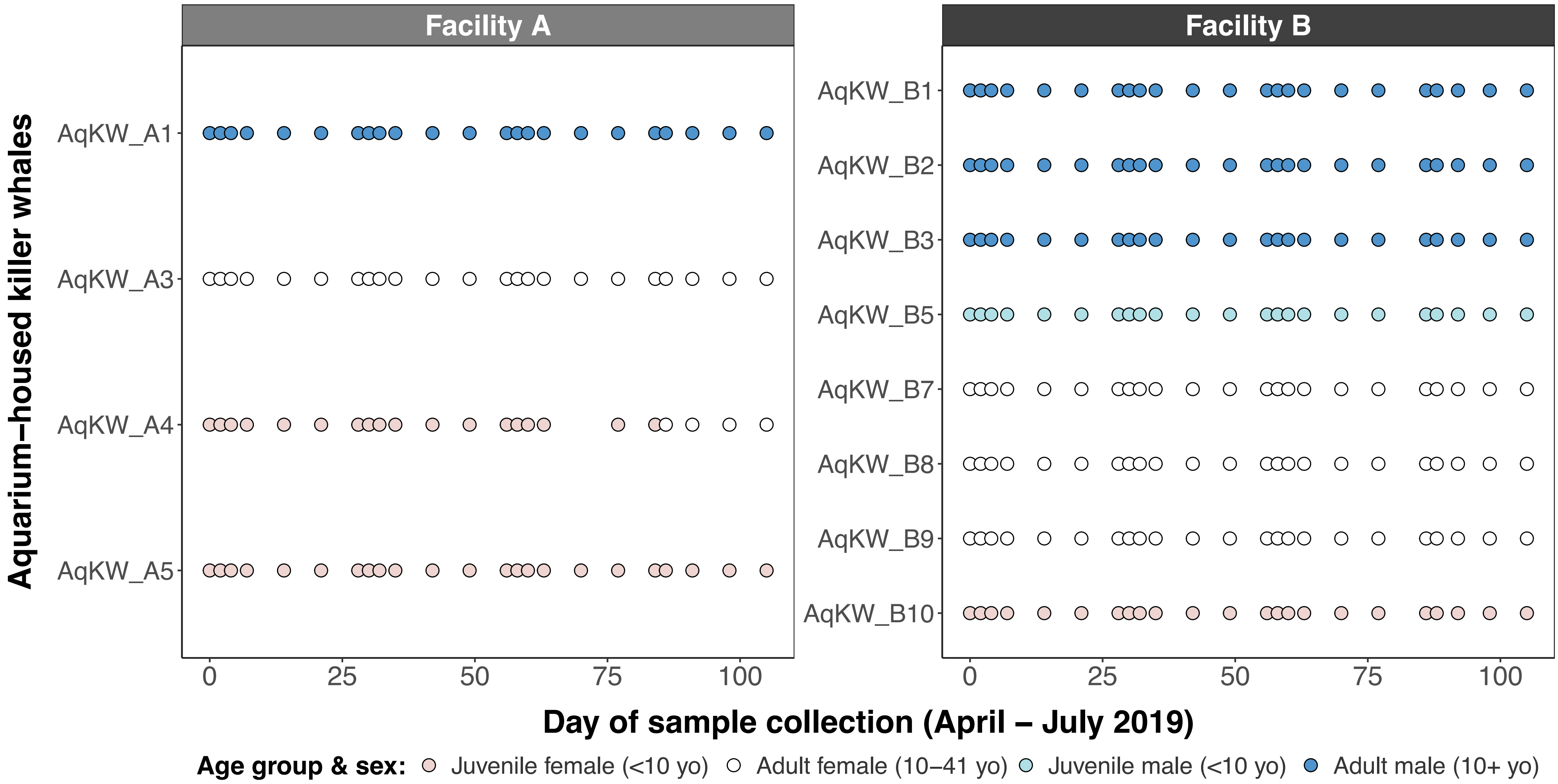


**Supplementary Figure 2**

**Supplementary Fig. 2. Sampling timeline for aquarium-housed killer whales.** Aquarium-housed killer whales were sampled 3 times a week for the first week of each month and then once weekly for the rest of each month. The dot plot shows the timeline of fecal samples that were collected at the two facilities and successfully sequenced. Dot fill denotes the sex and age class of individuals.

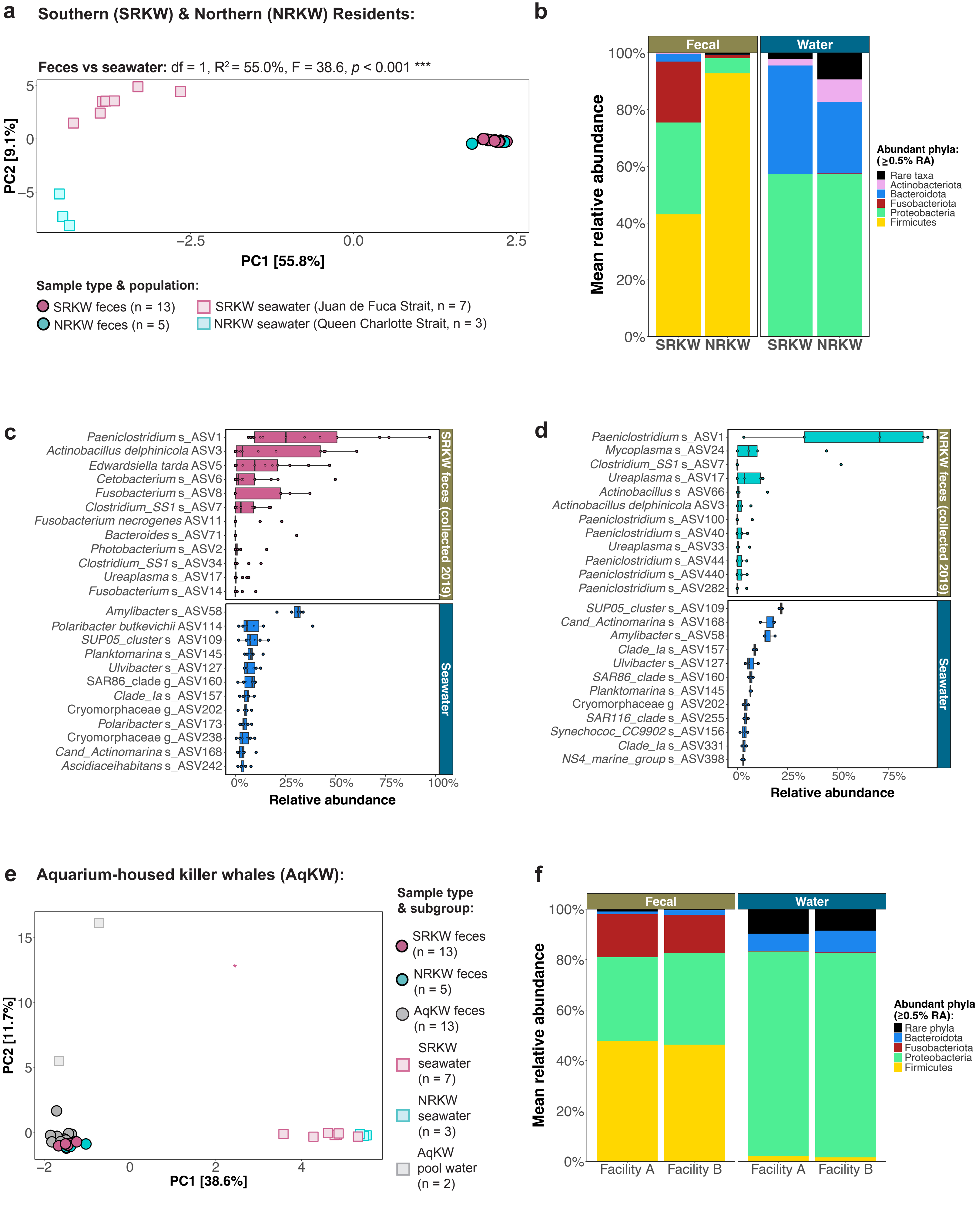


**Supplementary Figure 3**

**Supplementary Fig. 3. Killer whale fecal microbiotas were distinct from those of the surrounding water.** Surface water (seawater, n = 10; pool water, n = 2) was collected at the same time as the killer whale (KW) fecal samples during 2019. (**a**) PCA ordination of clr-transformed Aitchison distances among the Southern (SRKW) and Northern (NRKW) Resident fecal microbiotas and those of their surrounding seawater. (**b**) Stacked bar charts of the mean relative abundance (RA) of abundant phyla in the fecal and seawater microbiotas. Phyla with RA < 0.5% were defined as "Rare taxa". (**c, d**) RA boxplots of the most abundant ASVs in the (**c**) SRKW and (**d**) NRKW fecal microbiotas, and their surrounding seawater. Taxa are arranged in order of descending mean RA. (**e**) PCA ordination (as in **a**) of the wild and aquarium-housed (AqKW) KW fecal samples with seawater and pool water. AqKW fecal samples were normalized to 13 samples per individual. (**f**) Phylum-level relative abundance (as in **b**) in the AqKW fecal and pool water microbiotas. For all analyses shown here, only adult KWs (>9 years old) were considered.

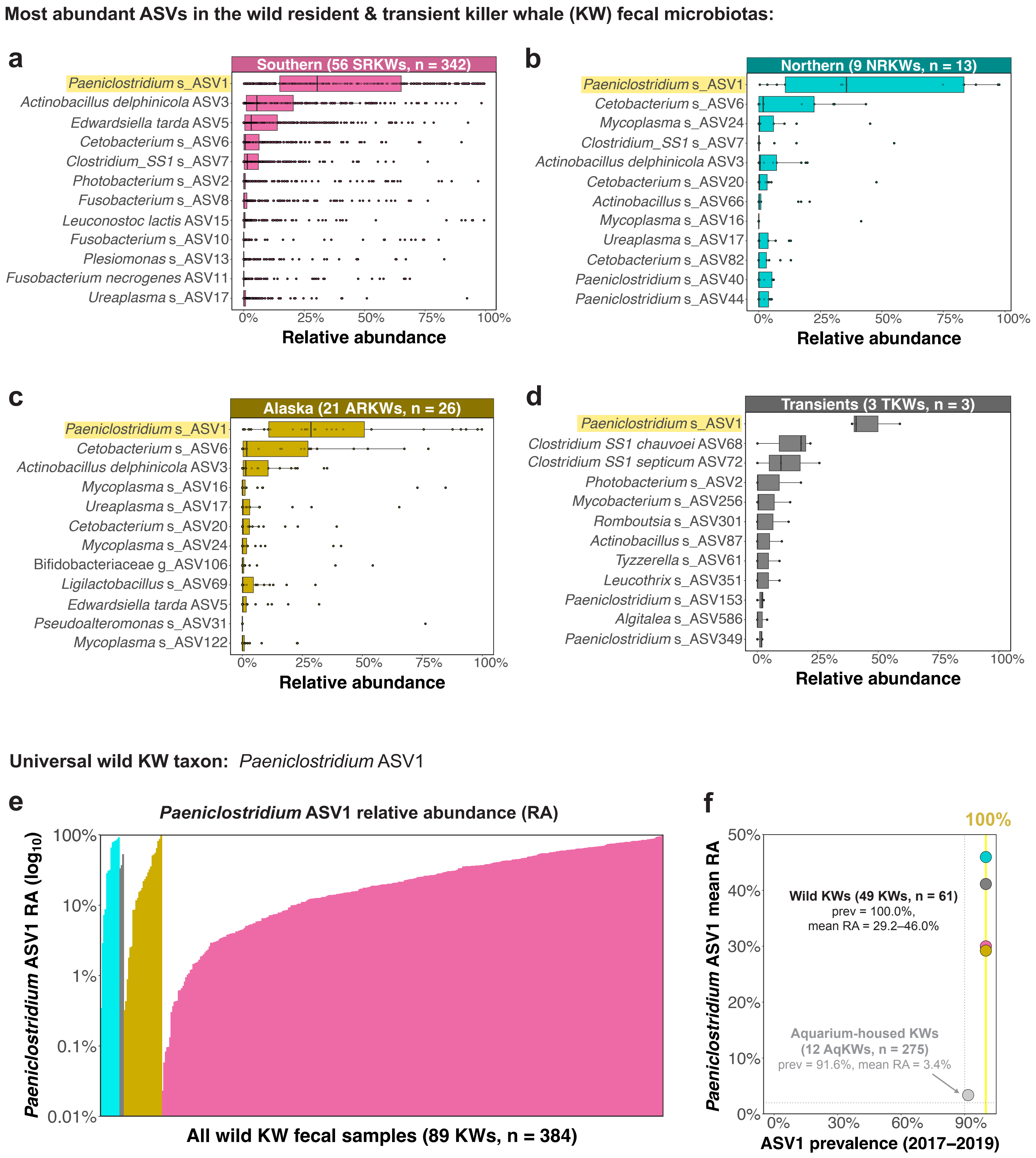


Supplementary Figure 4

**Supplementary Fig. 4. Core taxa were identified in the wild fecal microbiotas, including *Paeniclostridium* ASV1— a universal killer whale taxon regardless of diet or husbandry.** (**a–d**) Relative abundance (RA) boxplots of the most abundant ASVs in all fecal microbiotas from the four wild killer whale (KW) populations: the fish-eating (**a**) Southern (SRKW), (**b**) Northern (NRKW), and (**c**) Alaska (ARKW) Resident KWs and the marine mammal-eating (**d**) transient KWs (TKWs). Points represent individual samples. Taxa are arranged by descending mean RA. (**e**) Bar chart of *Paeniclostridium* ASV1 RA (the most abundant taxon in all KW populations) in all available fecal samples (arranged by population and RA). (**f**) Scatterplot of ASV1 prevalence and mean RA in each wild KW population and the aquarium-housed KWs, using only the fecal samples from 2017-2019. Yellow line highlights 100% prevalence. Dotted lines denote 90% prevalence and 2.0% RA.

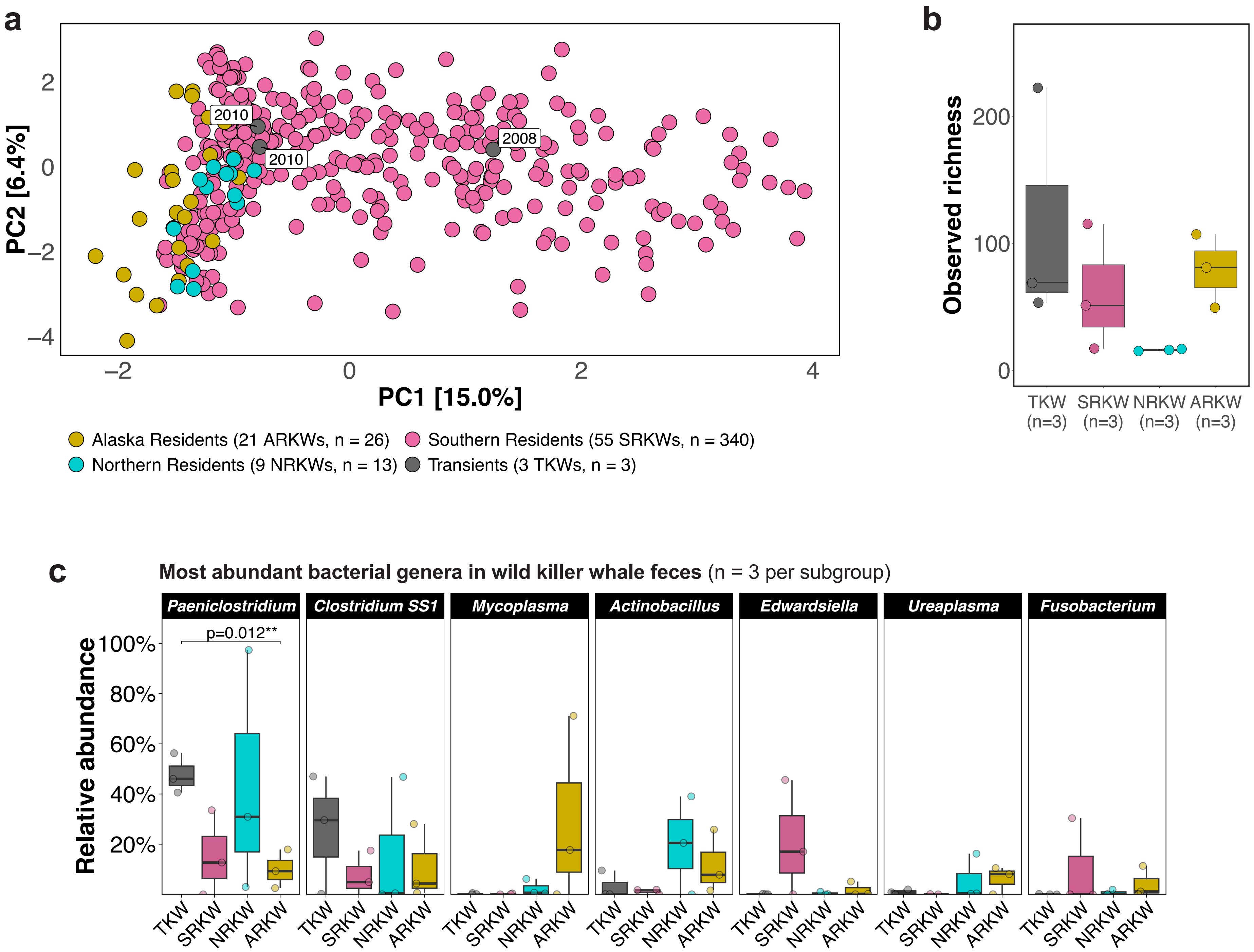


Supplementary Figure 5

**Supplementary Fig. 5. Transient killer whale fecal microbiotas were characterized by an abundance of *Paeniclostridium* ASV1 and *Clostridium SS1* spp.** (**a**) Principal component analysis of Aitchison distances comparing all fecal samples collected from wild killer whales (KWs). Point labels indicate the year of sample collection for the three transient KWs (TKWs, n = 3). (**b**) Boxplot comparison of observed richness in the wild KW microbiotas. Samples were restricted to years in which multiple subgroups (populations/ecotypes) were sampled (2008, 2010, and 2017–2019) and normalized to 3 samples per subgroup. (**c**) The seven most abundant genera are plotted in order of decreasing mean relative abundance. For **b** and **c**, t-tests were performed with Benjamini-Hochberg p-value adjustment to detect significant differences between the TKWs and each resident population.

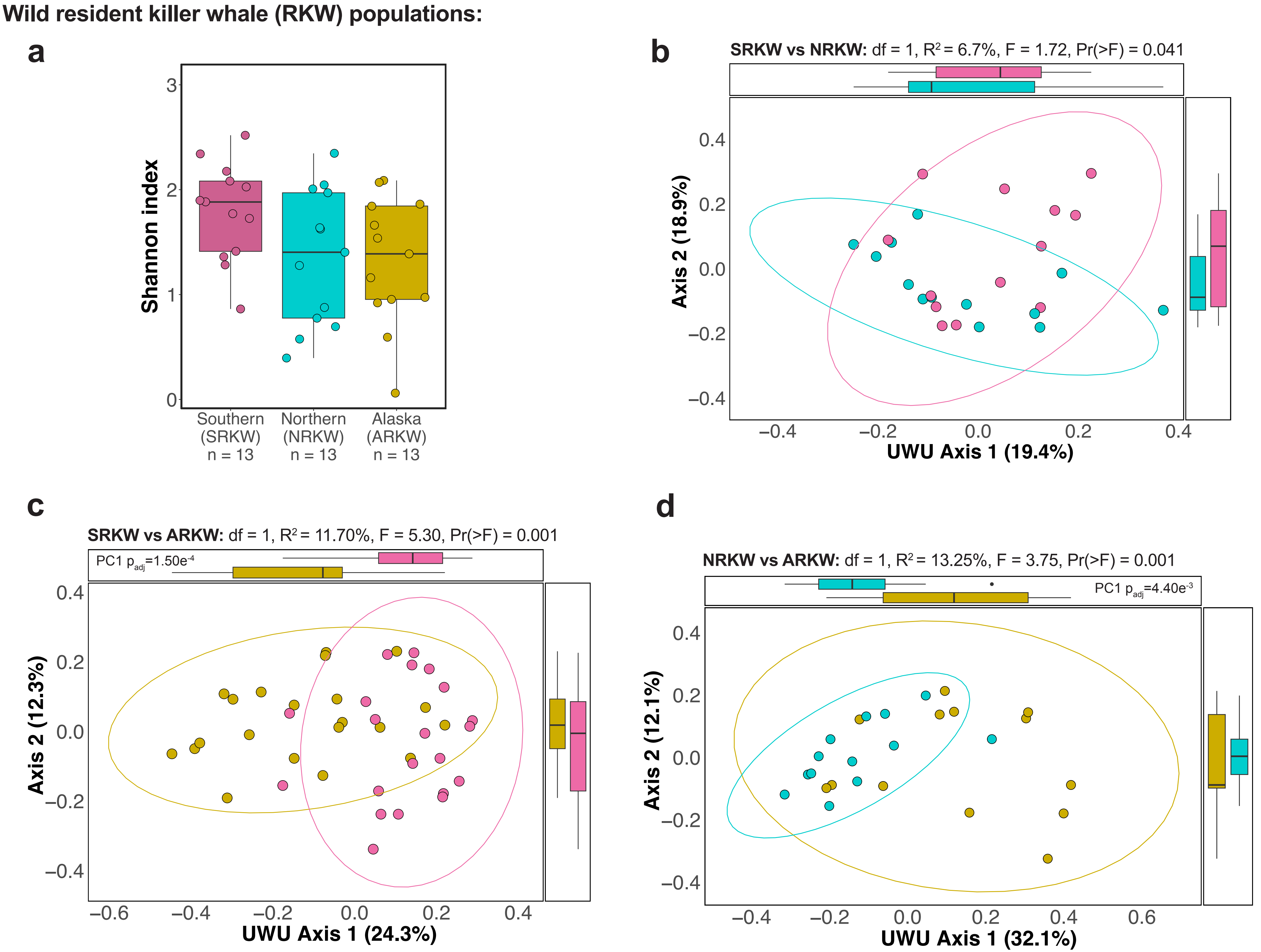


**Supplementary Figure 6**

**Supplementary Fig. 6. Fecal microbiotas of the wild resident killer whale populations were analyzed using alternative diversity measures.** (**a**) Boxplot of Shannon diversity. Shannon index of the fecal microbial read counts plotted for the Southern (SRKW), Northern (NRKW), and Alaska (ARKW) Resident killer whale populations. The dataset was restricted to fecal samples collected during the years when >1 population was sampled (2017–2019) and normalized (13 samples/population). A pairwise Dunn test with Benjamini-Hochberg p-value adjustment was performed (no significant differences identified). (**b–d**) Principal coordinate analysis (PCoA) of unweighted Unifrac distances (calculated from rarefied data) between the fecal microbiotas of the (**b**) SRKWs vs NRKWs (n = 13 each), (**c**) SRKWs vs ARKWs (n = 21 each), and (**d**) ARKWs vs NRKWs (n = 13 each). Title annotations denote the adonis2 results for the “Population” covariate. Margins show univariable distribution boxplots for each axis. Margin annotations indicate significant differences in axis loadings between populations (Dunn tests). Points, ellipses (95% CIs) and boxplots are colored by population.

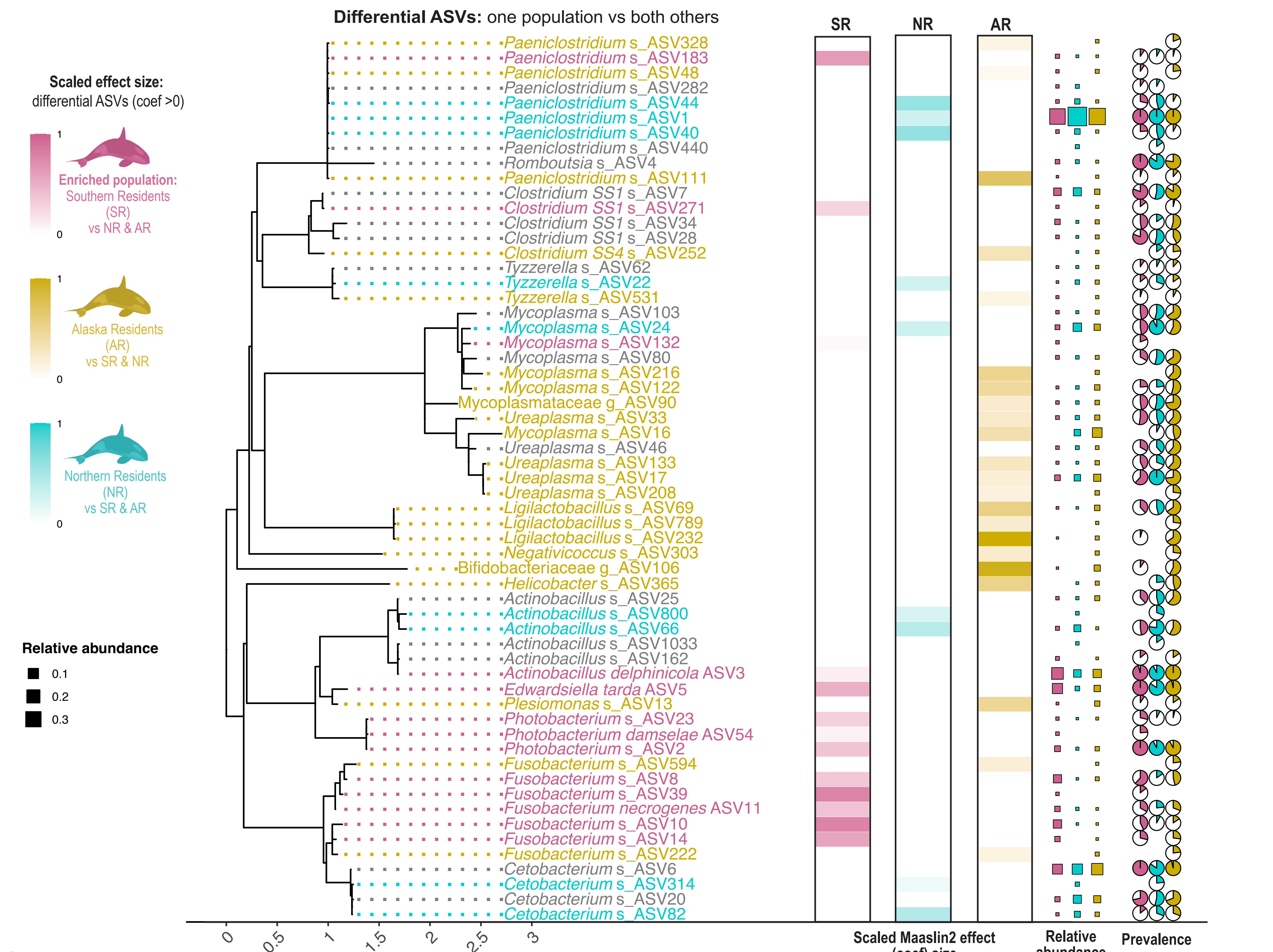


Supplementary Figure 7

**Supplementary Fig. 7. Full results of the differential abundance analysis comparing wild resident killer whale populations.** Amplicon sequence variant (ASV) differential abundance (DA) in fecal samples collected from the Southern (19 SRKWs; n = 22), Northern (9 NRKWs, n = 13) and Alaska (21 ARKWs, n = 26) Resident killer whale populations in 2017–2019. LEFT: Abundant ASVs (>1000 reads in ≥2 samples) were merged into a phylogeny. General and linear mixed models ('MaAsLin2') identified ASVs enriched in each population's fecal microbiotas versus those of both others. For significant ASVs (Benjamini-Hochberg adjusted p.adj <0.05), tip label color indicates the population in which it was enriched and the heatmap gradient shows the MaAsLin2 effect size (coef) scaled to the overall maximum value. RIGHT: Square size indicates ASV RA. Pie charts denote prevalence.

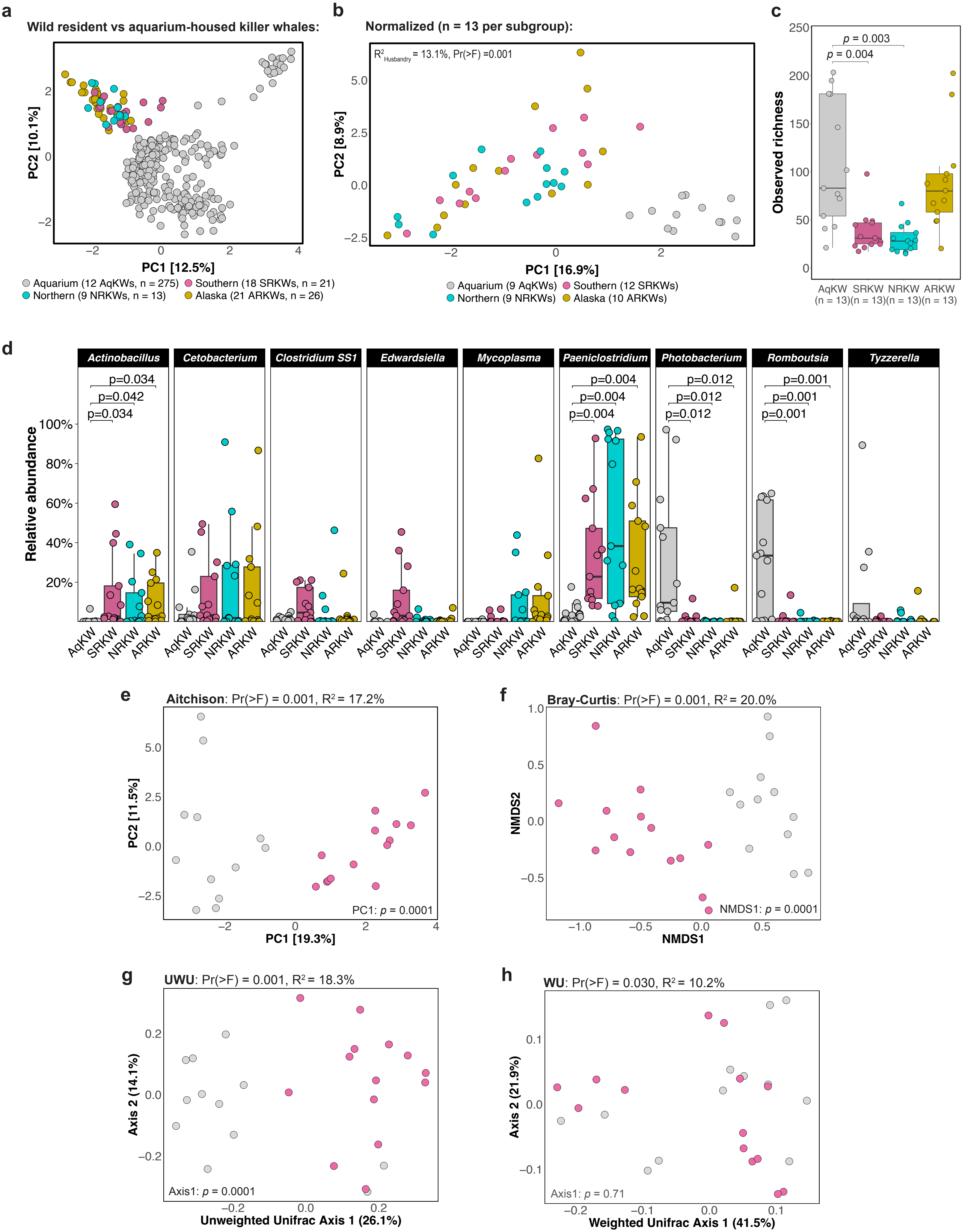


**Supplementary Figure 8**

**Supplementary Fig. 8. The fecal microbiotas of aquarium-housed killer whales were distinct from those of the wild resident populations.** (**a**) Principal component analysis (PCA) of clr-transformed Aitchison distances between killer whale (KW) fecal samples (points colored by KW subgroup). All samples from the aquarium-housed KWs (AqKWs) were included along with samples collected from the Southern (SRKW), Northern (NRKW) and Alaska (ARKW) Residents during 2017–2019. (**b**) PCA of the normalized dataset (n = 13 per KW subgroup) showing the adonis2 significance value and the proportion of variation explained ($R^2$) by husbandry (wild vs aquarium). (**c, d**) Boxplot comparisons of (**c**) richness and (**d**) relative abundance of the most abundant genera between the AqKWs and each resident population (n = 13 per subgroup). Pairwise t-tests with Benjamini-Hochberg p-value adjustment were used to identify significant differences. (**e–h**) PCA comparing the SRKW and AqKW microbiotas using different beta diversity distance metrics. Title annotations show adonis2 results for husbandry and plot annotations show significant differences in the primary axis loadings (t-test). Data were rarefied to the library size of the smallest population in **c**, **d**, **g** and **h**. UWU, unweighted Unifrac distance; WU, weighted Unifrac distance.

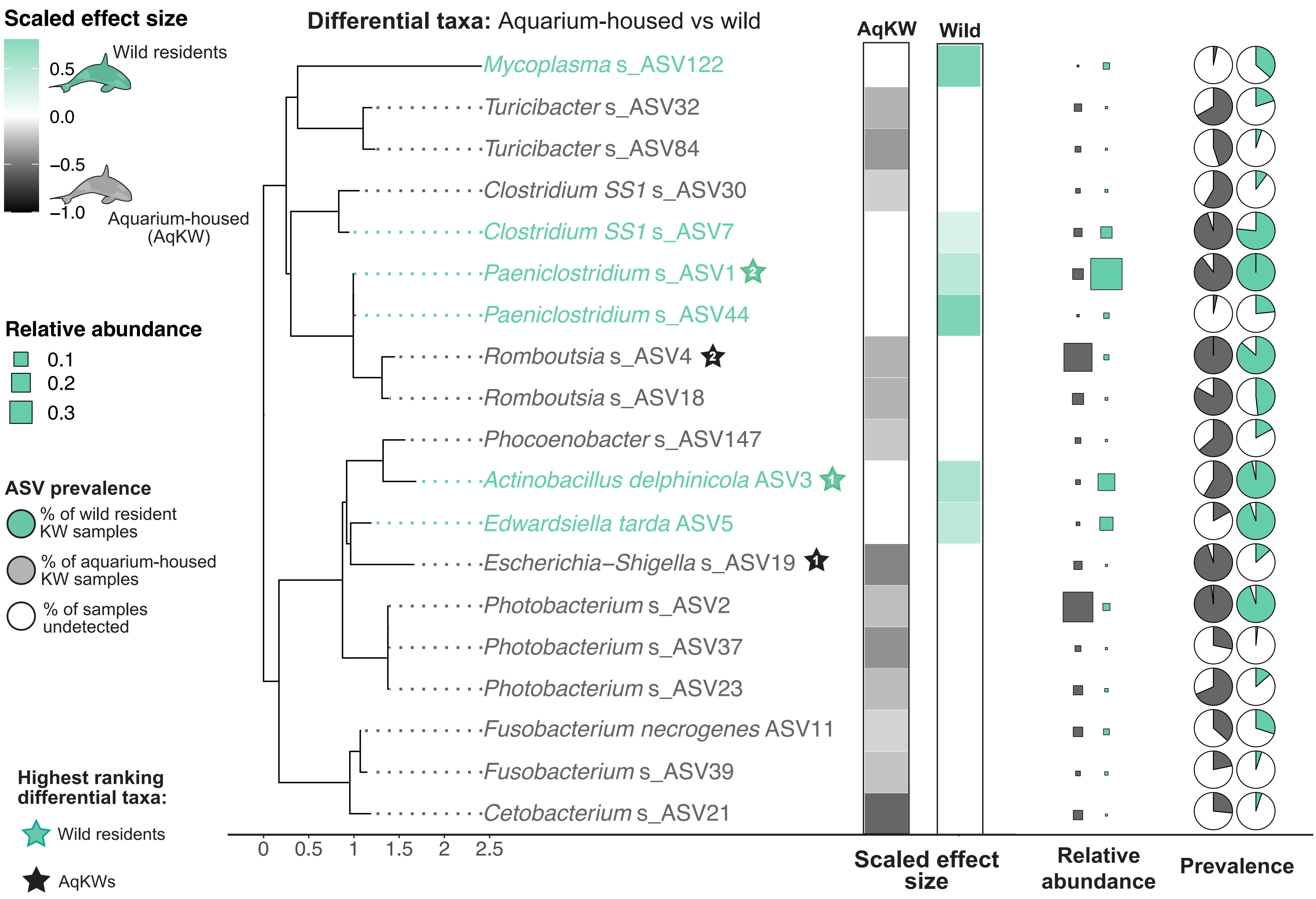


Supplementary Figure 9

**Supplementary Fig. 9.** ***Escherichia-Shigella* ASV19, *Romboutsia, and Photobacterium* spp. best distinguished the fecal microbiotas of aquarium-housed versus wild killer whales.** Phylogeny of taxa with differential abundance (DA) in the fecal microbiotas of the 12 aquarium-housed killer whales (AqKWs) versus all 60 samples collected from the three wild resident populations during 2017-2019: 19 Southern (n = 21), 13 Northern (n = 9), and 26 Alaska (n = 26) Resident individuals. The larger AqKW dataset was randomly normalized to n ≤ 5 per KW. LEFT: Abundant ASVs (>1000 reads in ≥5 samples) were merged into a phylogeny ('ggtree'). General and linear mixed models ('MaAsLin2') identified ASVs enriched in each population's fecal microbiotas versus the other. For significant ASVs (p.adj<0.05, correction = BH), tip label color indicates the population in which it was enriched and the heatmap gradient shows the MaAsLin2 effect size (coef) scaled to the overall maximum value. Stars indicate consensus between ≥3 DA methods (MaAsLin2, treeDA, ALDEx2, & coda4microbiome) (**Supplementary Table 4**) and the effect size rank (1=most differential). RIGHT: Square size indicates ASV RA. Pie charts ('scatterpie') denote prevalence.

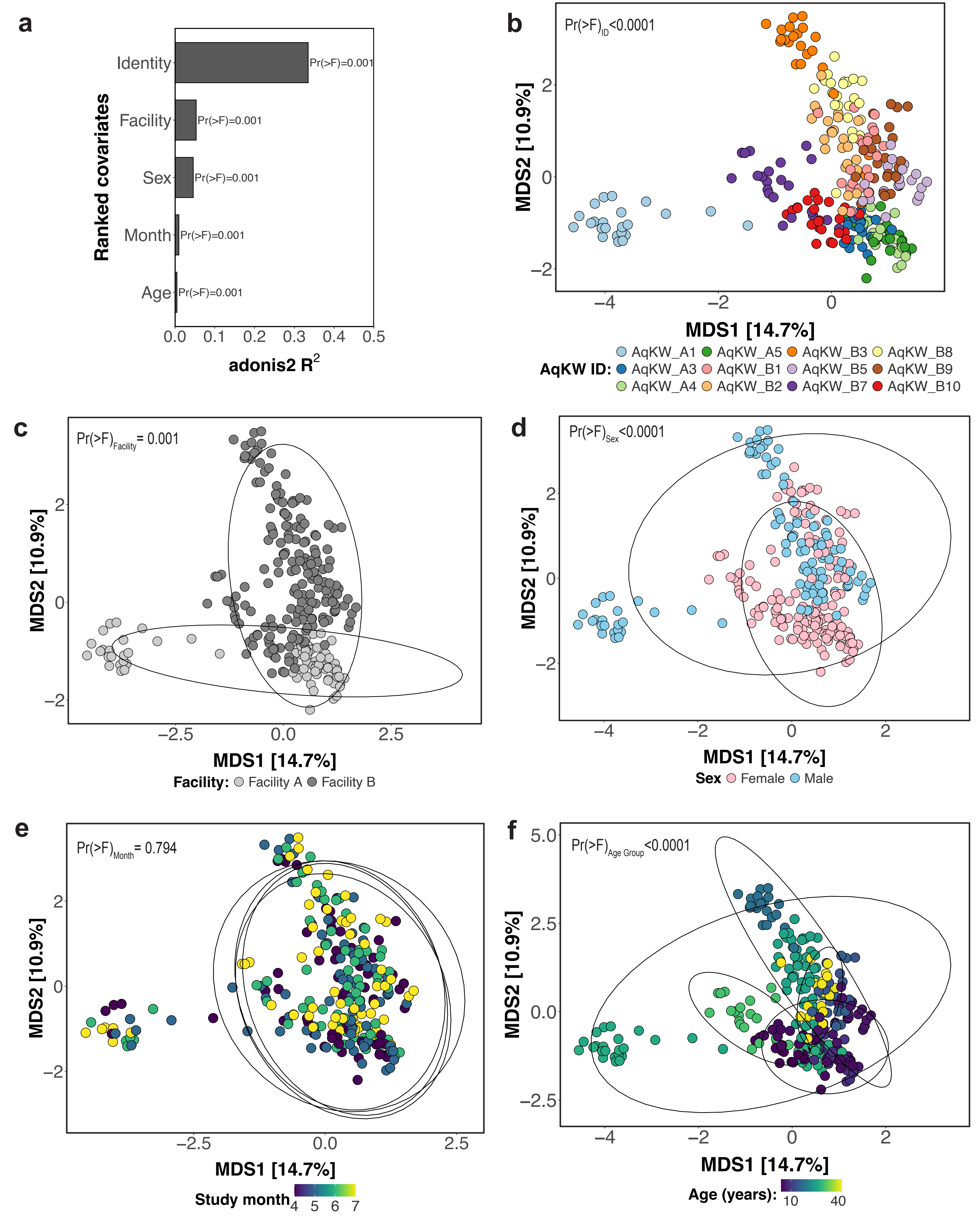


**Supplementary Figure 10**

**Supplementary Fig. 10. The individuality of the host was the strongest driver of variation in the fecal microbiota of aquarium-housed killer whales.** (**a**) Multivariate analysis of variance (MANOVA, 'adonis2', vegan v2.6-10) was performed on clr-transformed Aitchison distances between fecal samples collected from the aquarium-housed killer whales (AqKW). The bar chart shows study covariates ranked by the proportion of variance explained by each ($R^2$ values) and significance levels. (**b-f**) Principal component analysis of the Aitchison distance matrix. Points show individual samples colored by (**b**) AqKW identity, (**c**) facility, (**d**) sex, (**e**) study month, and (**f**) age. Ellipses show 95% confidence intervals for the groups denoted in the legend, except **f**, in which ellipses denote age bins (0–9, 10–19, 20–29, 30–39, ≥40 years old). Annotations show the homogeneity of group dispersions for each covariate ('betadisper", vegan); non-significant values indicate equal variance between groups.

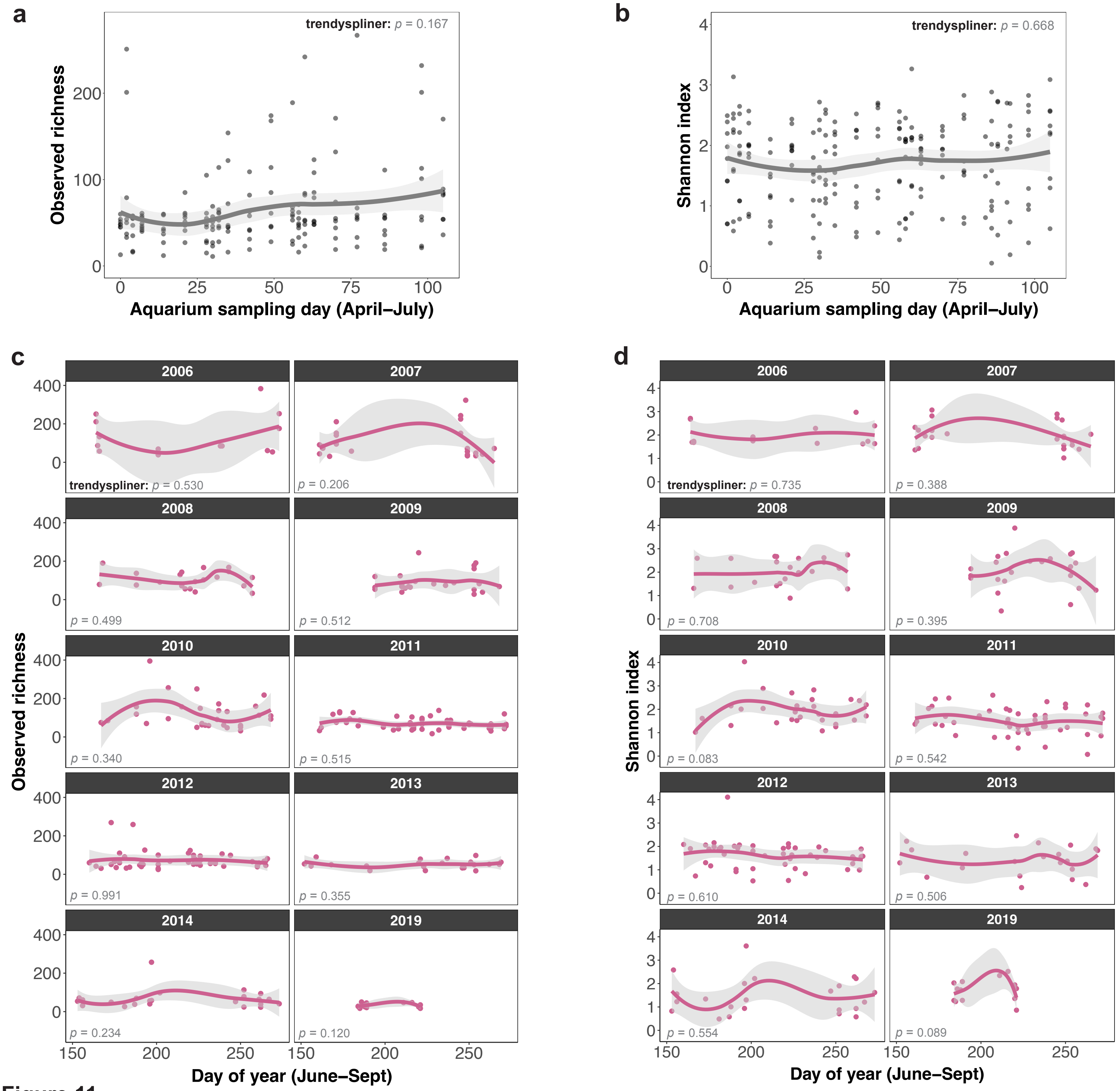


**Supplementary Figure 11**

**Supplementary Fig. 11. Alpha diversity in the fecal microbiotas of aquarium-housed and wild killer whales showed temporal stability over four-month timeframes.** (**a, b**) Alpha diversity in the aquarium-housed killer whale (AqKW) fecal microbiotas was plotted against the day of study. Data were rarefied to 16,236 reads/sample for (**a**) observed richness, and raw counts were used for (**b**) Shannon index. Juveniles (<10 years old) were excluded from the analysis. (**c, d**) Alpha diversity in the wild Southern Resident killer whale (SRKW) fecal microbiotas was plotted against time using an equivalent time frame as the AqKW sampling period, i.e., 105 days. Data were restricted to samples collected June-Sept (months with the most frequent sampling) and only study years with >10 samples. Juveniles were excluded. Data were rarefied to 23,712 reads/sample for (**a**) richness, and raw counts were used for (**b**) Shannon index. For **a-d**, points represent individual fecal samples and lines represent Loess smoothing of the data and gray shading indicates the 95% confidence intervals. Annotations show the p-values of permutation tests designed to detect non-zero temporal trends (‘trendyspliner’, splinectomeR, v0.1.0).

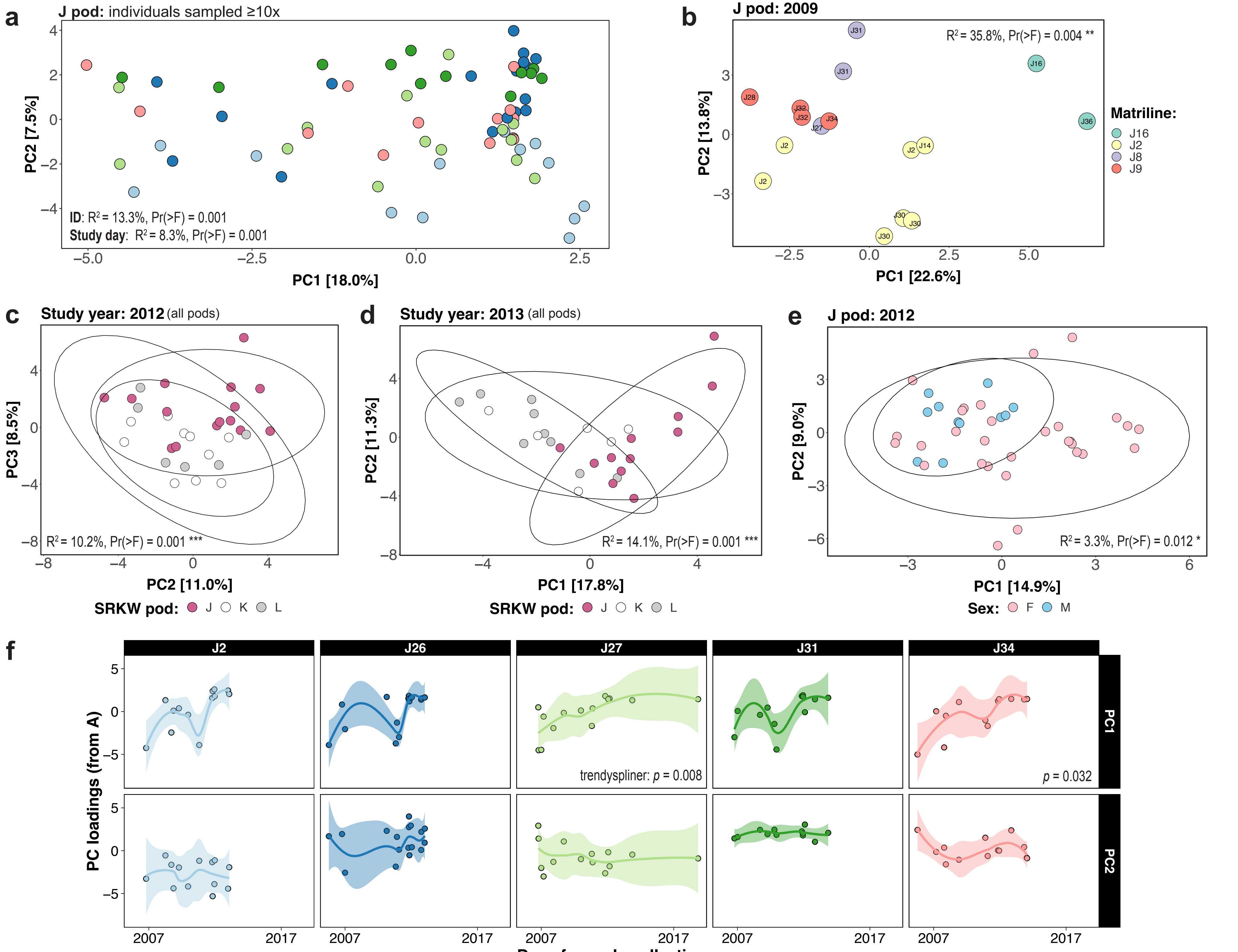

Supplementary Figure 12

**Supplementary Fig. 12. Social structure, ancestry and individuality influence the structure of Southern resident killer whale fecal communities.** Principal component analysis of clr-transformed Aitchison distances between adult (>9 years old) Southern Resident killer whale (SRKW) fecal microbiotas colored by (**a**) SRKW ID, (**b**) matriline, (**c, d**) pod, and (**e**) sex. The principal components that best clustered the data are shown. Annotations denote the multivariate analysis of variance (MANOVA, 'adonis2', vegan v2.6-10) significance level and $R^2$ value. For analysis of ID (**a**), data from all study years are shown for adults in J pod (the most frequently sampled pod) that were sampled ≥10x and survived ≥3 years after their last sample. Point colors correspond IDs shown in (**f**). For analysis of matriline (**b**), data were restricted to 2009 (the only year with all matrilines sampled). For analysis of pod (**c**, **d**), data were restricted to 2012 and 2013 (the only years with >5 samples per pod) and normalized to ≤15 samples per pod. Ellipses indicate 95% confidence intervals. **(f)** PC1 and PC2 loadings from (**a**) were plotted for individuals against time. Lines represent Loess smoothing of the data with shading indicating the 95% confidence intervals (colored by ID as in (**a**)).

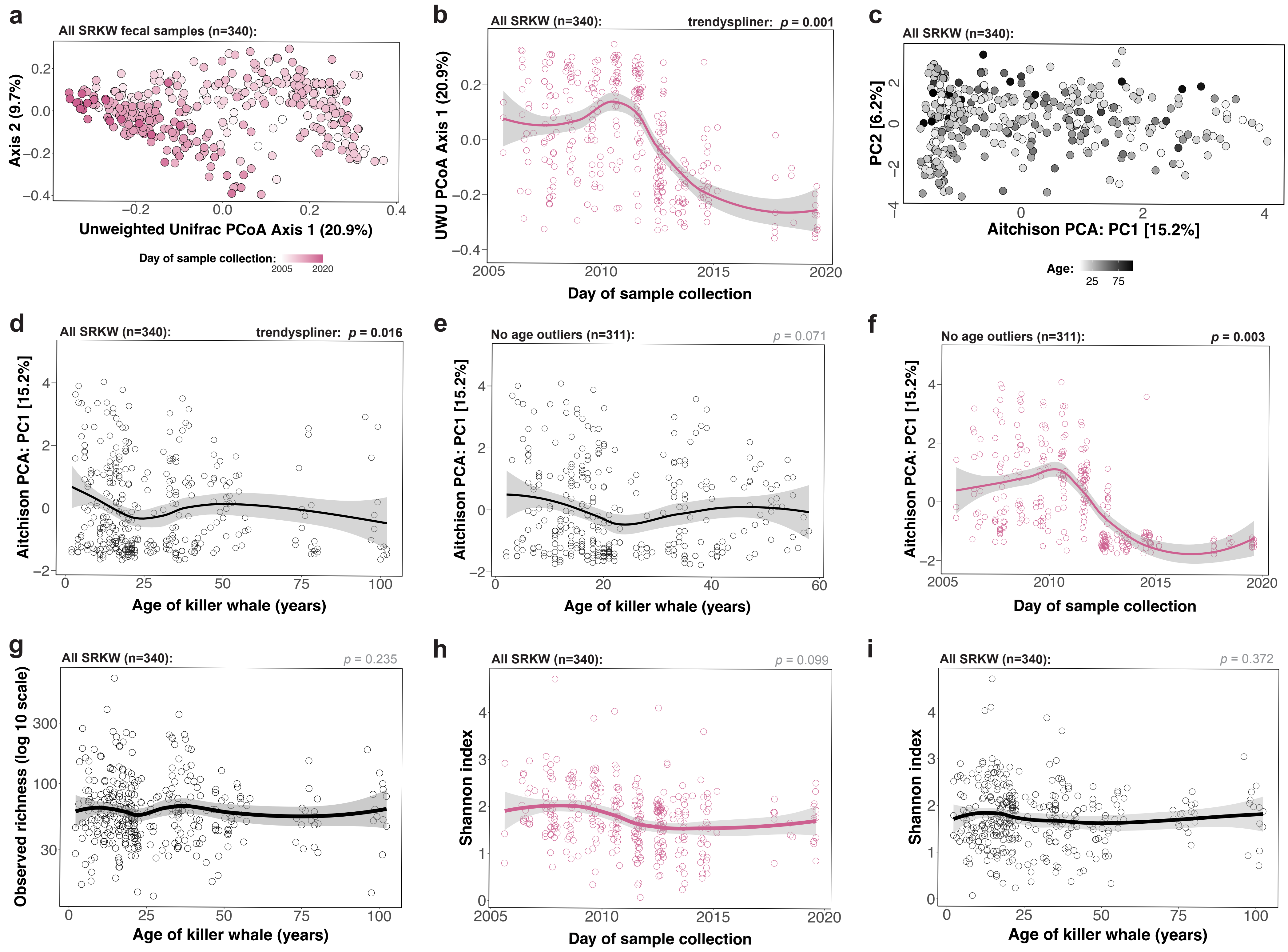


Supplementary Figure 13

**Supplementary Fig. 13. Fecal microbiota temporal changes were robust to alternative diversity distance metrics and independent of aging in SRKWs <70 years old.** (**a**) Principal coordinate analysis (PCoA) of unweighted Unifrac distances using rarefied data (23,712 reads/sample) from all Southern Resident killer whale (SRKW) fecal samples (n = 340, color indicates year). (**b**) Axis 1 loadings from (**a**), plotted against date of sample collection. (**c**) Principal component analysis (PCA) of clr-transformed Aitchison distances (color indicates age). (**d**) PC1 loadings from (**c**) plotted against SRKW age (n = 340). (**e**) Same analysis as in (**d**) but without samples from SRKWs ≥70 years old (J2, J8, and K11; n=29). (**f**) Same analysis as in **Fig. 4b** but without samples from SRKWs ≥70 years old (J2, J8, and K11; n=29). (**g**) Richness plotted against SRKW age (data normalized to 23,712 reads/sample). (**h, i**) Shannon index (using reads counts) plotted against (**h**) day and (**i**) SRKW age. Lines show Loess smoothing of the data (confidence intervals = 95%).

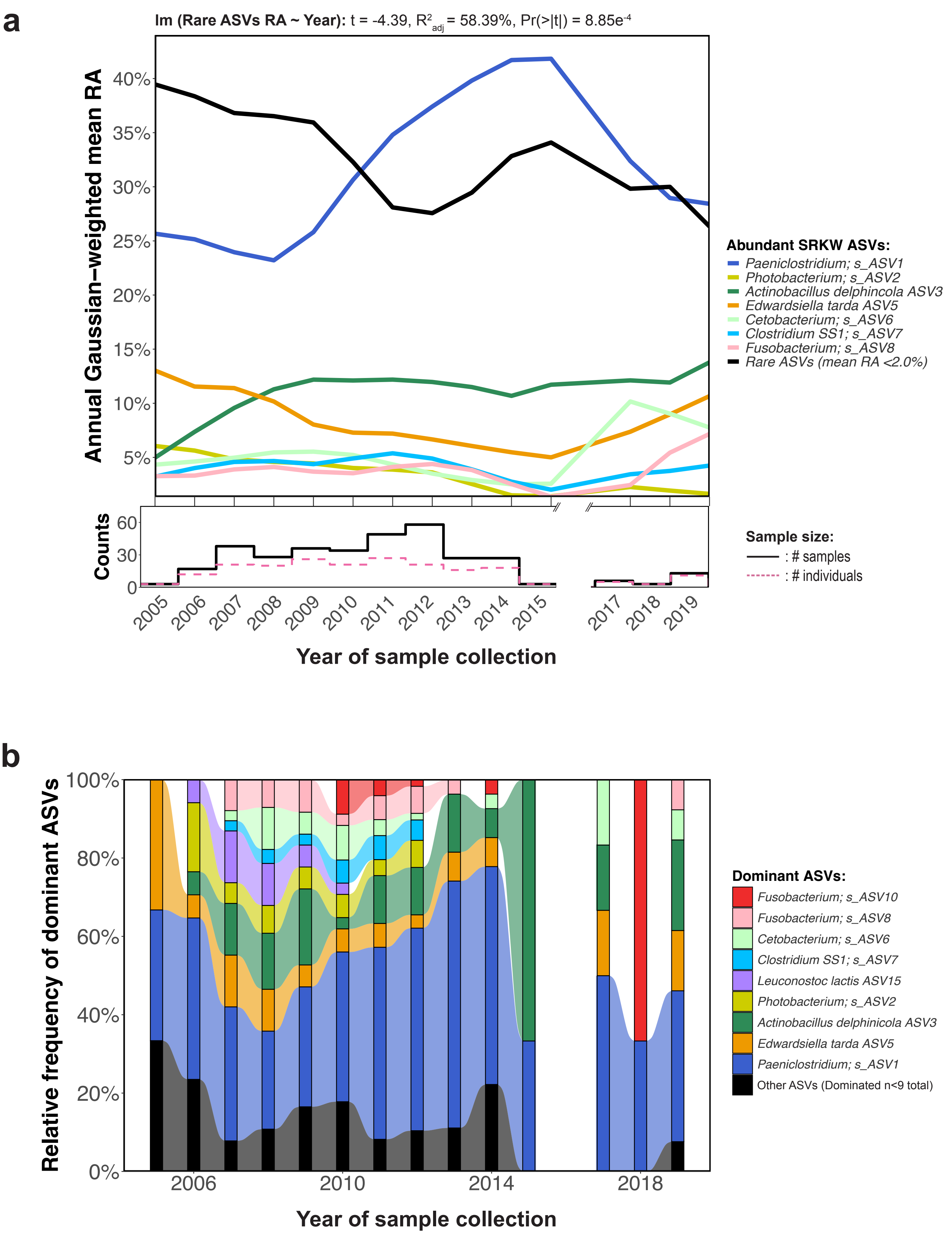


**Supplementary Figure 14**

**Supplementary Fig. 14. The relative abundances of rare taxa in the SRKW fecal microbiotas declined over time as several abundant ASVs became dominant.** (**a**) Relative abundance (RA) of the most abundant ASVs (RA >2.0%) in the SRKW microbiotas over time. For each ASV, lines denote the Gaussian-weighted mean RA averaged over consecutive 2-year windows. Less abundant genera were classified as "Rare taxa", and a simple linear regression model ('lm', stats) was used to identify a non-zero temporal trend in this group. The step chart below shows the sample size and number of SRKW individuals sampled each year. (**b**) Alluvial graph of annual ASV-level dominance. The dominant taxon in each fecal sample was identified ('dominant_taxa', microbiomeutilities) and the annual relative frequency of samples dominated by each was determined. For each year, taxa that were dominant in <9 samples were grouped as "Other". The visual break denotes 2016, when samples were not collected.

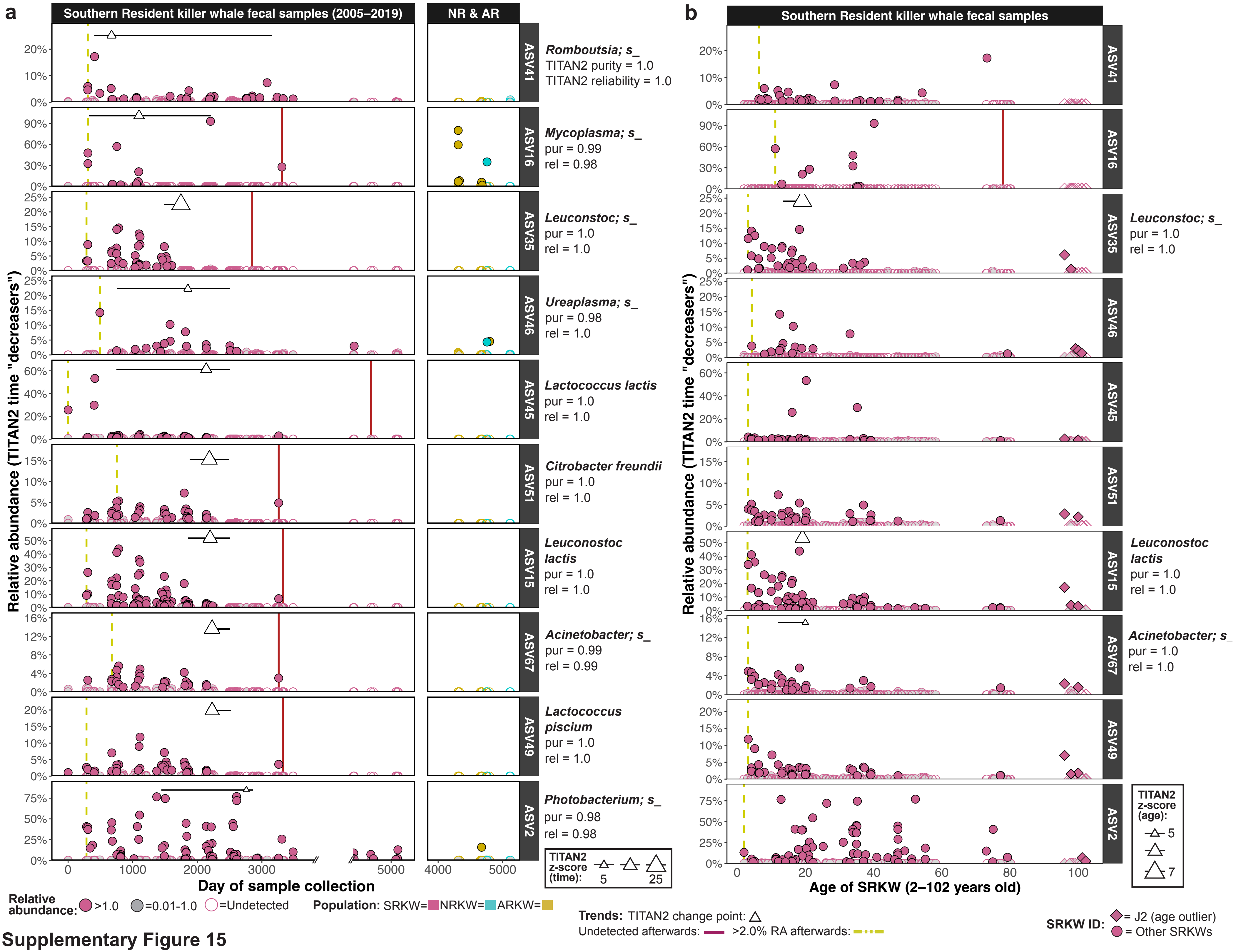


Supplementary Figure 15

**Supplementary Fig. 15. Several taxa from the Southern Resident killer whale (SRKW) fecal microbiotas became undetected over time in an age-independent manner.** TITAN2 (nBoot=500) was used to identify "decreasers" (purity and reliability scores ≥97.0%, **Supplementary Table 6**) in the SRKW microbiotas - ASVs that exhibited a community-level decrease in relative abundance (RA) and/or prevalence over the study period. (**a**) For TITAN2 decreasers with mean RA >0.25%, RA is plotted against the day of sample collection. Right column shows RA in the other resident KW populations for comparison. Point color indicates KW population and fill denotes whether an ASV was abundant (color fill, ≥1.0% RA), scarce (gray fill, RA=0.01–1.0%) or undetected (no fill) in a sample. Yellow dotted lines indicate the time when ASVs first became achieved >2.0% RA. Red solid lines indicate the time after which an ASV could no longer be detected. Triangles mark the TITAN2 median change point; their size indicates z score (normalized magnitude of change from mean). Horizontal lines show 5–95% quantiles from the bootstrapped distribution. (**b**) To assess the degree of confounding by host age, the same ASVs were plotted against SRKW host age as in (**a**). The TITAN2 analysis was repeated using host age. Purity and reliability scores are shown at the right for ASVs identified as having a significant association with age. Diamond-shaped points denote J2, a longevity outlier (~96–102 years old).

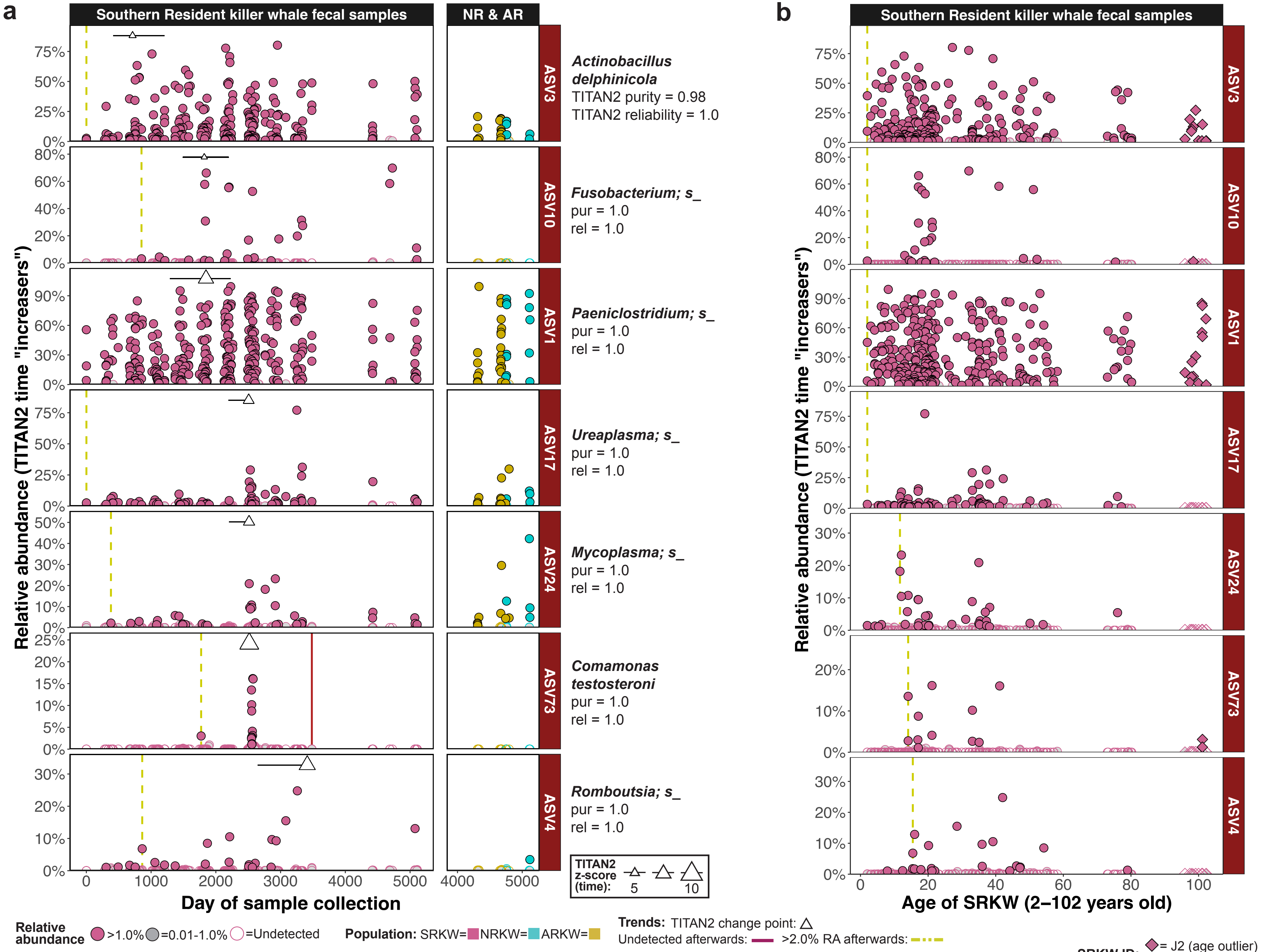


Supplementary Figure 16

**Supplementary Fig. 16. TITAN2 "increaser" ASVs in the Southern Resident killer whale (SRKW) fecal microbiotas that became abundant or bloomed over time in an age-independent manner.** TITAN2 (nBoot=500) was used to identify "increasers" (purity and reliability scores ≥97.0%, **Supplementary Table 6**) in the SRKW microbiotas - ASVs that exhibited a community-level increase in relative abundance (RA) and/or prevalence over the study period. (**a**) For TITAN2 increasers with mean RA >0.25% in the SRKW microbiotas, RA was plotted against the day of the study. The right column shows RA in the other resident populations for comparison. Point color indicates KW population and fill denotes whether an ASV was abundant (color fill, ≥1.0% RA), scarce (gray fill, RA=0.01–1.0%) or undetected (no fill) in a sample. Yellow dotted lines indicate the time point when ASVs first achieved >2.0% RA. Red solid lines indicate the time after which an ASV could no longer be detected. Triangles mark the TITAN2 median change point; their size indicates z score (normalized magnitude of change from mean). Horizontal lines show 5–95% quantiles from the bootstrapped distribution. (**b**) To assess the degree of confounding by host age, the same ASVs were plotted against SRKW host age as in (**a**). The TITAN2 analysis was repeated using host age. Purity and reliability scores were all non-significant. Diamond-shaped points denote J2, a longevity outlier (~96–102 years old).

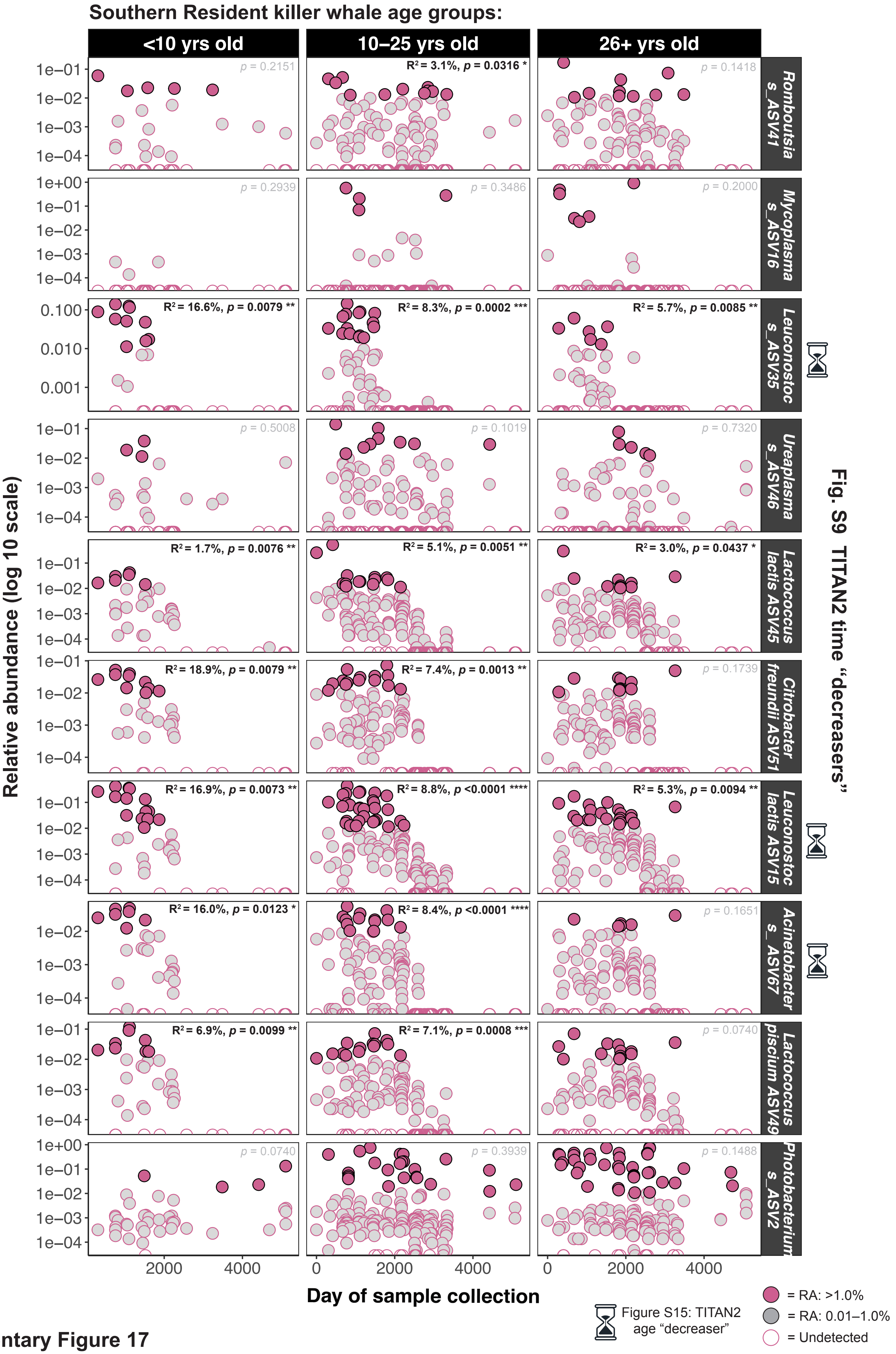


**Supplementary Figure 17**

**Supplementary Fig. 17. The abundance of the Leuconostoc ASVs identified as both TITAN2 time and age-associated "decreasers" declined over time in all Southern Resident killer whale (SRKW) age groups.** The relative abundance (RA) of TITAN2 time-associated "decreaser" ASVs from **Supplementary Figure 15** was plotted against the day of study and faceted by SRKW age group. Point fill indicates whether an ASV was abundant (pink fill, ≥1.0% RA), scarce (gray fill, RA=0.01–1.0%) or undetected (no fill) in each sample. Hourglass symbols indicate ASVs that were also identified as TITAN2 age-associated decreasers (**Supplementary Table 7**). For each ASV and age group, annotations denote significant (black text) and non-significant (gray text) p-values and marginal $R^2$ values from autoregressive general mixed-effect linear models that regressed RA against day of study with 'SRKW_ID' as a random effect ('lme', nlme).

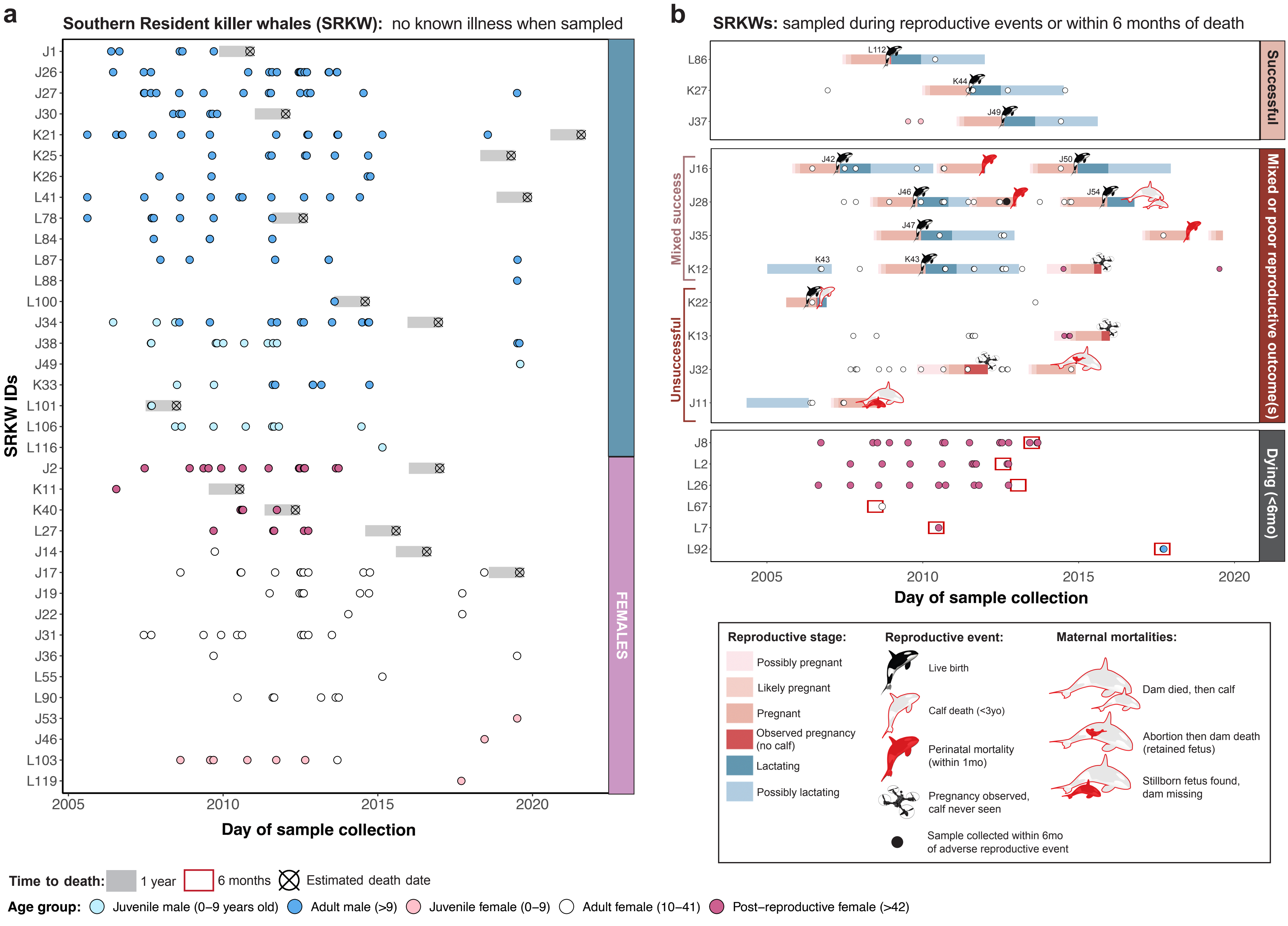


**Supplementary Figure 18**

**Supplementary Fig. 18. The dataset captured several births and adverse health events in the Southern resident killer whale (SRKW) population.** (**a**) Timeline of samples collected from SRKWs that were not known to be ill, pregnant, or lactating at the time and did not have a known adverse health event within 6 months of sampling. Points are colored by age and sex. Circled x's indicate the estimated date of death (based on when a SRKW was no longer seen with its pod), and the gray shading denotes the year prior to death. (**b**) Timeline for SRKWs that were sampled during a reproductive event or within 6 months of death (indicated by red boxes). Reproductive stages were estimated based on the date when the calf was first observed and the average duration of pregnancy and lactation in captivity (Duffield et al., 1995). The drone icon indicates when a female previously observed to be pregnant was seen no longer pregnant and without a calf.

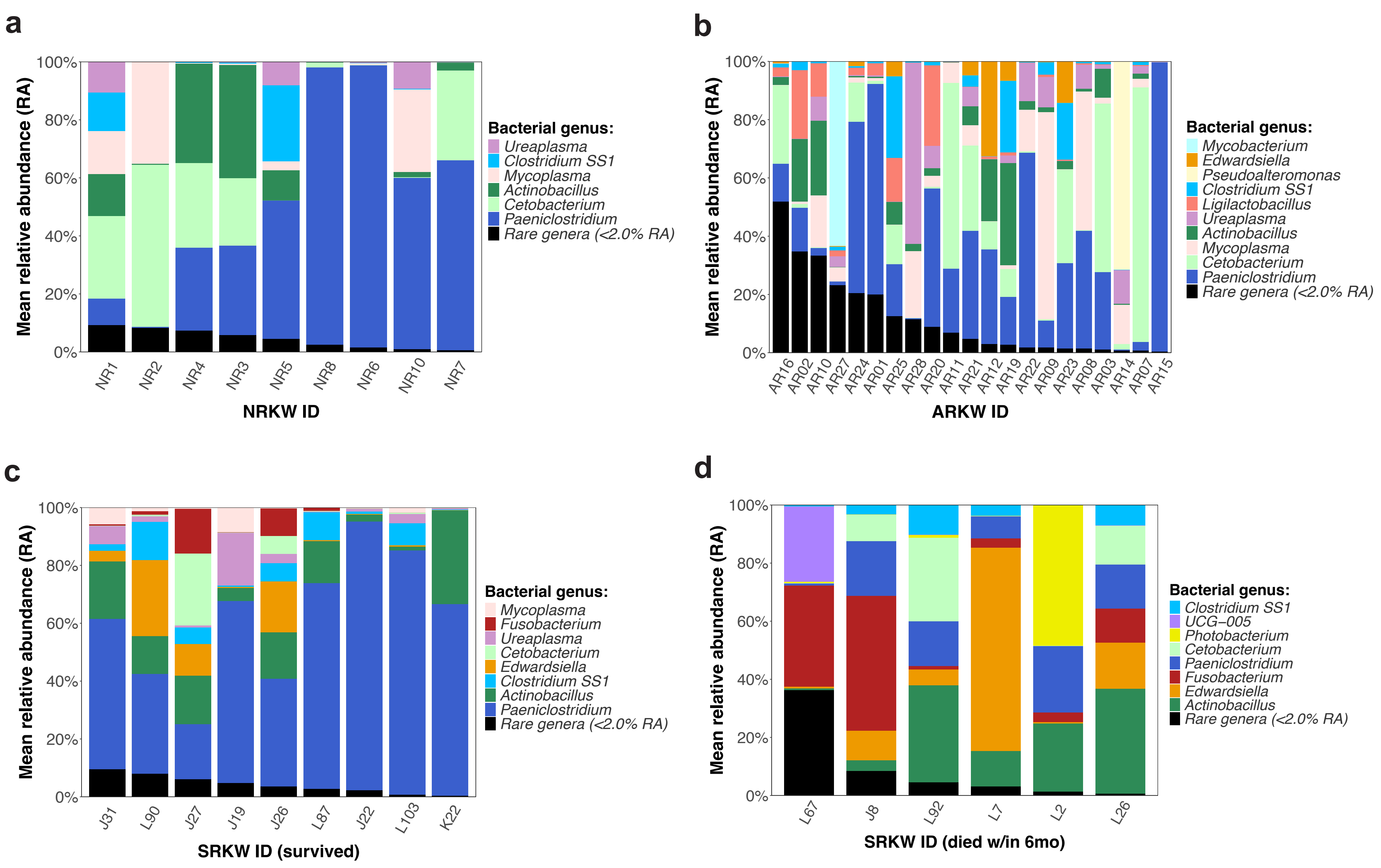


**Supplementary Figure 19**

**Supplementary Fig. 19. Abundance of *Fusobacterium* was highest in the Southern Resident killer whale (SRKW) population but varied between individuals, and dying SRKWs had less abundant *Paeniclostridium* compared to those who survived.** The most abundant genera (RA >2.0%) in the (**a**) Northern and (**b**) Alaska Residents, (**c**) the SRKWs that survived to Nov 2024, and (**d**) the SRKWs sampled within 6 months of death are shown in stacked bar plots, with less abundant genera (RA <2.0%) classified as "Rare genera". For the SRKW population (which had age data available), samples from juveniles (<9 years old) were excluded. Each bar denotes mean abundance per KW individual and bars are ordered by decreasing abundance of the rare genera.

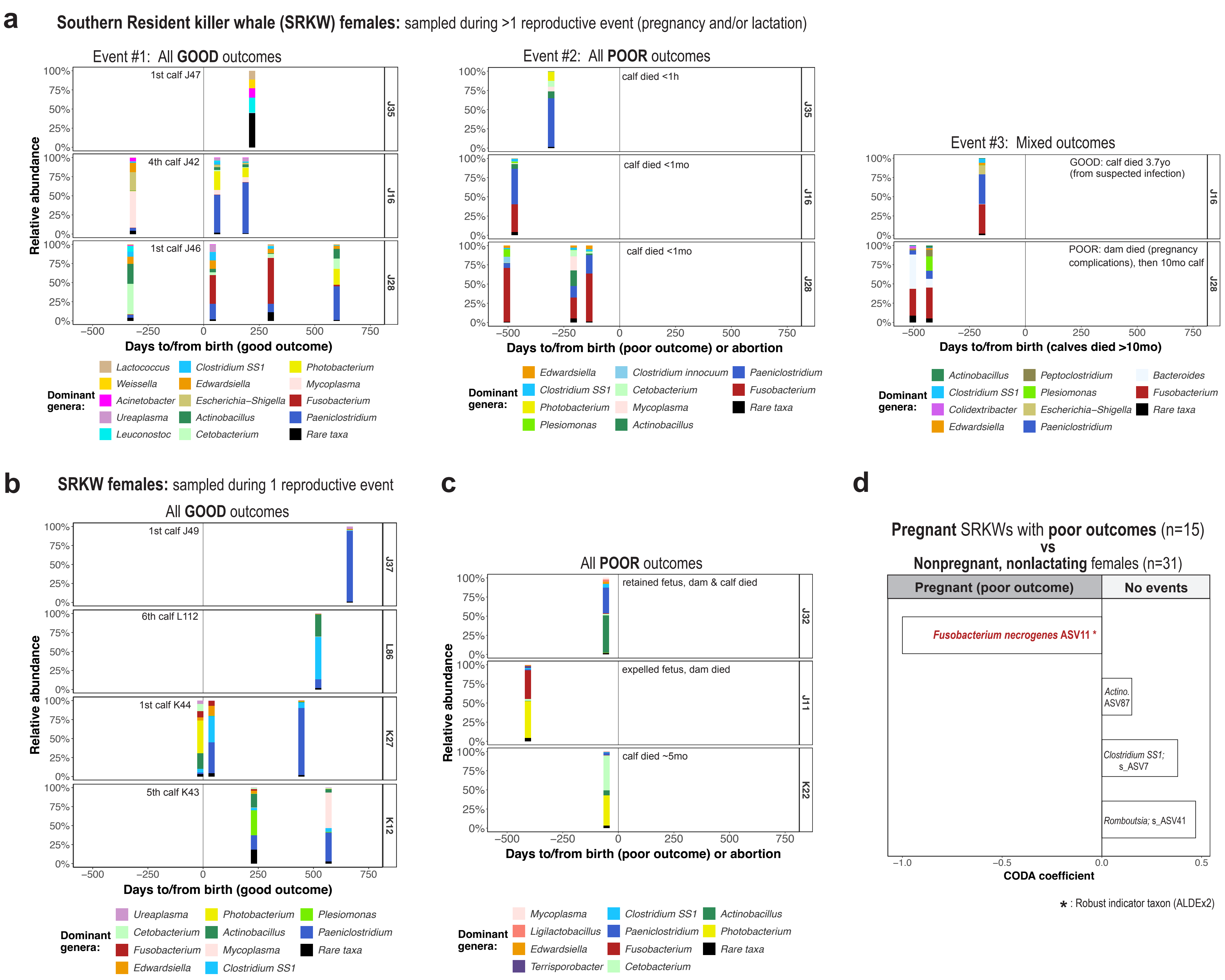


Supplementary Figure 20

**Supplementary Fig. 20. Microbial relative abundances in the fecal microbiotas of nonpregnant, and pregnant Southern Resident killer whales (SRKWs) with different outcomes.** (**a–c)** Genus-level relative abundance (RA) in the SRKW fecal samples (n = 40) collected during pregnancy (n = 24) or lactation (n = 16). Excluding pregnancies with no observed calf (**Supplementary Fig. 18**), 10 SRKWs were analyzed: (**a**) 3 SRKWs sampled during 2–3 reproductive events with mixed outcomes, (**b**) 4 SRKWs sampled during one good outcome event, and (**c**) 3 SRKWs sampled during one unsuccessful pregnancy. Day = 0 is the estimated date of calf birth or full-term abortion. Samples collected ≤3 weeks apart were merged for visibility. For each plot, the 4 most abundant genera per sample and 10 most abundant genera overall are shown arranged in order of increasing overall RA. (**d**) ASV differential abundance (DA) between samples collected from females in the reproductive event cohort while nonpregnant/nonlactating (n = 31) and those collected during unsuccessful pregnancies (n = 15). DA taxa were identified using *coda-lasso* (coda4microbiome, nfolds=5); coefficient size is shown as a bar graph. Asterisks denote robust taxa also identified as DA via ALDEx2 (**Supplementary Table 11**).
