## Supplementary Tables for "Fifteen-year microbiome survey of endangered killer whales (*Orcinus orca*) reveals declining diversity and population differences"

**Supplementary Table 1. Information about the killer whales (KWs) sampled for this study.**

| KW population | SRKW ID | # samples | First sample | Last sample | Age (yrs) | Sex | Matriline | Pod |
|---|---|---|---|---|---|---|---|---|
| Southern Resident (SRKW) | J1 | 5 | 05/21/06 | 09/13/09 | 54 - 58 | M | J2 | J |
| | J2 | 13 | 06/20/07 | 09/25/13 | 95 - 102 | F | J2 | J |
| | J8 | 15 | 09/26/06 | 09/10/13 | 73 - 80 | F | J8 | J |
| | J11 | 4 | 05/24/06 | 06/20/07 | 33 - 34 | F | J8 | J |
| | J14 | 1 | 09/25/09 | | 35 - 35 | F | J2 | J |
| | J16 | 7 | 06/15/06 | 06/05/14 | 33 - 41 | F | J16 | J |
| | J17 | 11 | 08/15/08 | 06/11/18 | 31 - 40 | F | J9 | J |
| | J19 | 8 | 07/04/11 | 09/24/17 | 32 - 38 | F | J8 | J |
| | J22 | 2 | 01/18/14 | 09/24/17 | 28 - 32 | F | J9 | J |
| | J26 | 17 | 06/13/06 | 09/08/13 | 14 - 22 | M | J16 | J |
| | J27 | 14 | 06/12/07 | 07/03/19 | 15 - 28 | M | J8 | J |
| | J28 | 16 | 06/20/07 | 10/05/14 | 13 - 21 | F | J9 | J |
| | J30 | 6 | 05/22/08 | 10/22/09 | 12 - 14 | M | J2 | J |
| | J31 | 12 | 06/10/07 | 07/07/14 | 11 - 19 | F | J8 | J |
| | J32 | 16 | 09/06/07 | 10/05/14 | 11 - 18 | F | J9 | J |
| | J34 | 13 | 06/13/06 | 09/19/14 | 7 - 16 | M | J9 | J |
| | J35 | 5 | 07/15/10 | 09/19/17 | 12 - 19 | F | J9 | J |
| | J36 | 2 | 09/10/09 | 07/03/19 | 10 - 20 | F | J16 | J |
| | J37 | 3 | 07/13/09 | 06/02/14 | 8 - 12 | F | J2 | J |
| | J38 | 12 | 09/02/07 | 07/30/19 | 4 - 16 | M | J9 | J |
| | J46 | 1 | 06/14/18 | | 8 - 8 | F | J9 | J |
| | J49 | 2 | 08/08/19 | 08/09/19 | 7 - 7 | M | J2 | J |
| | J53 | 1 | 07/04/19 | | 4 - 4 | F | J9 | J |
| | K11 | 1 | 07/20/06 | | 73 - 73 | F | K7 | K |
| | K12 | 13 | 09/19/06 | 07/08/19 | 35 - 48 | F | K4 | K |
| | K13 | 9 | 10/08/07 | 09/19/14 | 35 - 42 | F | K7 | K |
| | K21 | 14 | 08/12/05 | 07/22/18 | 19 - 32 | M | K18 | K |
| | K22 | 2 | 06/14/06 | 08/11/13 | 18 - 26 | F | K4 | K |
| | K25 | 7 | 08/24/09 | 09/09/14 | 18 - 23 | M | K7 | K |
| | K26 | 4 | 12/12/07 | 10/05/14 | 14 - 21 | M | K3 | K |
| | K27 | 5 | 12/17/06 | 07/21/14 | 12 - 20 | F | K7 | K |
| | K33 | 8 | 07/06/08 | 09/24/14 | 7 - 13 | M | K4 | K |
| | K40 | 5 | 07/26/10 | 09/28/11 | 47 - 48 | F | K3 | K |
| | L2 | 10 | 09/06/07 | 10/02/12 | 44 - 52 | F | L2 | L |
| | L7 | 1 | 07/07/10 | | 49 - 49 | F | L26 | L |
| | L26 | 9 | 08/27/06 | 10/01/12 | 50 - 56 | F | L26 | L |
| | L27 | 5 | 09/10/09 | 10/02/12 | 44 - 47 | F | L4 | L |
| | L41 | 9 | 08/12/05 | 06/03/14 | 28 - 36 | M | L12 | L |
| | L55 | 1 | 02/24/15 | | 37 - 37 | F | L4 | L |
| | L67 | 1 | 09/13/08 | | 23 - 23 | F | L2 | L |
| | L78 | 7 | 08/13/05 | 08/04/11 | 16 - 22 | M | L2 | L |
| | L84 | 4 | 10/05/07 | 08/05/11 | 17 - 21 | M | L9 | L |
| | L86 | 1 | 05/25/10 | | 18 - 18 | F | L4 | L |
| | L87 | 6 | 12/23/07 | 07/08/19 | 15 - 27 | M | L28 | L |
| | L88 | 1 | 07/05/19 | | 26 - 26 | M | L2 | L |
| | L90 | 6 | 06/20/10 | 09/26/13 | 16 - 20 | F | L26 | L |
| | L92 | 2 | 09/19/17 | 09/26/17 | 22 - 22 | M | L26 | L |
| | L100 | 2 | 08/08/13 | 08/09/13 | 12 - 12 | M | L35 | L |
| | L101 | 2 | 09/06/07 | 09/11/07 | 5 - 5 | M | L2 | L |
| | L103 | 7 | 08/15/08 | 09/11/13 | 5 - 10 | F | L4 | L |
| | L106 | 8 | 06/14/08 | 06/22/14 | 2 - 8 | M | L4 | L |
| | L116 | 1 | 02/24/15 | | 4 - 4 | M | L4 | L |
| | L119 | 1 | 09/13/17 | | 5 - 5 | F | L12 | L |
| | no_ID_01 | 1 | 08/04/19 | | calf | | | |
| | no_ID_02 | 1 | 08/09/19 | | calf | | | |
| | no_ID_13 | 1 | 08/08/19 | | calf | | | |

| KW population | KW ID | # samples | First sample | Last sample | Age (yrs) | Sex | Pod |
|---|---|---|---|---|---|---|---|
| Northern Resident (NRKW) | NRKW_01 | 1 | 08/21/18 | | | | A1 |
| | NRKW_02 | 1 | 08/21/18 | | | | A1 |
| | NRKW_03 | 1 | 08/22/18 | | | | I11 |
| | NRKW_04 | 1 | 08/22/18 | | | | I11 |
| | NRKW_05 | 2 | 08/22/18 | 08/17/19 | | | A1_A5 |
| | NRKW_06 | 1 | 08/22/18 | | | | A1_A5 |
| | NRKW_07 | 3 | 08/11/18 | 08/09/19 | | | A1_A5_I11 |
| | NRKW_08 | 1 | 08/09/19 | | | | A1_A5_I11 |
| | NRKW_10 | 2 | 08/09/19 | 08/18/19 | | | A1_A5_I11 |
| Alaska Resident (ARKW) | ARKW_01 | 1 | 06/06/18 | | | | AD31 or AD54 |
| | ARKW_02 | 1 | 05/29/18 | | | M | AD8 |
| | ARKW_03 | 3 | 06/06/16 | 06/06/18 | | M | AD8 or AD35 |
| | ARKW_07 | 1 | 05/30/17 | | | | AK6 |
| | ARKW_08 | 2 | 06/01/17 | 05/31/18 | | F | AD16/AK |
| | ARKW_09 | 1 | 06/01/17 | | | | AD16/AK |
| | ARKW_10 | 1 | 06/01/17 | | | M | AD16/AK |
| | ARKW_11 | 1 | 06/01/17 | | | M | AD16/AK |
| | ARKW_12 | 1 | 06/01/17 | | | F | AD16/AK |
| | ARKW_14 | 1 | 06/20/17 | | | F | many |
| | ARKW_15 | 1 | 06/22/17 | | | M | AK2 |
| | ARKW_16 | 1 | 06/22/17 | | | F | AK2 |
| | ARKW_19 | 1 | 05/09/18 | | | M | AE |
| | ARKW_20 | 1 | 05/24/18 | | | F | AD8 only |
| | ARKW_21 | 1 | 05/24/18 | | | M | AK9 |
| | ARKW_22 | 3 | 05/26/18 | 06/05/18 | | M | AD8 |
| | ARKW_23 | 1 | 05/27/18 | | | M | AD8 |
| | ARKW_24 | 1 | 05/29/18 | | | M | AD8 |
| | ARKW_25 | 1 | 06/04/18 | | | F | AX48s |
| | ARKW_27 | 1 | 08/07/18 | | | M | |
| | ARKW_28 | 1 | 09/30/18 | | | | |
| Transient (TKW) | TKW_01 | 1 | 06/30/08 | | | | |
| | TKW_02 | 1 | 10/26/10 | | | | |
| | TKW_03 | 1 | 10/27/10 | | | | |
| Aquarium-Housed (AqKW) | AqKW_A1 | 23 | 04/15/19 | 07/29/19 | 27 - 28 | M | |
| | AqKW_A3 | 23 | 04/15/19 | 07/29/19 | 28 - 28 | F | |
| | AqKW_A4 | 22 | 04/15/19 | 07/29/19 | 9 - 10 | F | |
| | AqKW_A5 | 23 | 04/15/19 | 07/29/19 | 5 - 6 | F | |
| | AqKW_B1 | 23 | 04/08/19 | 07/22/19 | 41 - 42 | M | |
| | AqKW_B10 | 23 | 04/08/19 | 07/22/19 | 4 - 5 | F | |
| | AqKW_B2 | 23 | 04/08/19 | 07/22/19 | 26 - 26 | M | |
| | AqKW_B3 | 23 | 04/08/19 | 07/22/19 | 18 - 18 | M | |
| | AqKW_B5 | 23 | 04/08/19 | 07/22/19 | 6 - 6 | M | |
| | AqKW_B7 | 23 | 04/08/19 | 07/22/19 | 31 - 31 | F | |
| | AqKW_B8 | 23 | 04/08/19 | 07/22/19 | 26 - 26 | F | |
| | AqKW_B9 | 23 | 04/08/19 | 07/22/19 | 14 - 15 | F | |

**Supplementary Table 2. Universal and core taxa identified in the killer whale fecal microbiotas.** The prevalence and relative abundance (RA) of each ASV was calculated for each wild resident killer whale (KW) population, the transient KWs, and the aquarium-house KWs (AqKW) and only those with a mean prevalence >1.0% and RA >0.1% across KW subgroups are shown. For the Southern Resident killer whales (SRKW), the dataset was restricted to the years in which the other resident populations were also sampled (2017–2019). ASVs that were prevalent (≥50%) and abundant (mean RA ≥1%) for a KW population were considered "core" members of their microbiotas and are highlighted according to KW population. Values that met the core criteria are shown in black text, sub-threshold values are shown in grey text, and values <0.01% are not shown. Bright yellow highlighting denotes ASV1, a "universal" taxon that was a core member in all wild KW populations (resident and transient). Green highlighting denotes the "resident KW core" ASVs for all three resident KW populations.

| | | WILD RESIDENT KILLER WHALES (2017-2019) | | | | | | | | | |
|---|---|---|---|---|---|---|---|---|---|---|---|
| | | SRKW (n=22) | | NRKW (n=13) | | ARKW (n=26) | | TKW | | AqKW | |
| CORE TAXA | | RA | Prevalence | RA | Prevalence | RA | Prevalence | RA | Prevalence | RA | Prevalence |
| *Paeniclostridium; s_ASV1* | UNIVERSAL | 29.98% | 100.00% | 45.96% | 100.00% | 29.21% | 100.00% | 41.13% | 100.00% | 3.37% | 91.64% |
| *Cetobacterium; s_ASV6* | | 8.78% | 95.45% | 9.98% | 84.62% | 13.16% | 96.15% | | 100.00% | 3.07% | 88.36% |
| *Clostridium SS1; s_ASV7* | CORE: | 4.02% | 77.27% | 5.00% | 53.85% | 1.54% | 84.62% | | 100.00% | 1.19% | 93.45% |
| *Actinobacillus delphinicola ASV3* | Wild Resident | 12.90% | 95.45% | 4.80% | 92.31% | 5.02% | 100.00% | | 100.00% | 0.55% | 62.91% |
| *Cetobacterium; s_ASV20* | populations | 1.24% | 68.18% | 4.69% | 53.85% | 3.73% | 69.23% | | | 0.62% | 47.27% |
| *Ureaplasma; s_ASV17* | | 1.61% | 59.09% | 2.46% | 100.00% | 3.17% | 73.08% | | | | 0.36% |
| *Photobacterium; s_ASV2* | | 1.83% | 100.00% | 0.10% | 92.31% | 0.69% | 92.31% | 5.43% | 100.00% | 25.73% | 98.18% |
| *Fusobacterium; s_ASV8* | | 5.41% | 59.09% | | 15.38% | 0.66% | 46.15% | | 66.67% | 0.91% | 25.09% |
| *Edwardsiella tarda ASV5* | | 9.56% | 95.45% | 0.89% | 84.62% | 2.45% | 96.15% | 0.12% | 100.00% | 0.14% | 21.82% |
| *Actinobacillus; s_ASV25* | | 0.80% | 40.91% | 0.16% | 46.15% | 1.04% | 61.54% | 0.11% | 66.67% | 0.28% | 48.73% |
| *Ureaplasma; s_ASV33* | | 0.29% | 50.00% | 0.51% | 46.15% | 1.00% | 61.54% | | | | |
| Mycoplasma; s_ASV122 | | | 22.73% | 0.23% | 23.08% | 1.30% | 53.85% | | | | 5.45% |
| *Ligilactobacillus; s_ASV69* | | | 36.36% | | 46.15% | 1.75% | 65.38% | | 66.67% | | 0.36% |
| *Bifidobacteriaceae; g_ASV106* | | | 9.09% | | | 2.57% | 57.69% | | | | |
| *Mycoplasma; s_ASV24* | | 0.89% | 45.45% | 5.44% | 92.31% | 2.07% | 57.69% | 0.19% | 66.67% | | 0.36% |
| *Actinobacillus; s_ASV66* | | 0.38% | 45.45% | 3.64% | 76.92% | | 57.69% | | | | |
| *Paeniclostridium; s_ASV349* | | | 4.55% | | | | | 1.08% | 66.67% | | 2.55% |
| *Paeniclostridium; s_ASV153* | | | | 0.45% | 15.38% | | | 1.31% | 66.67% | | |
| *Mycobacterium; s_ASV256* | | | | | | | | 3.80% | 66.67% | | |
| *Tyzzerella; s_ASV61* | | | | | | | 7.69% | 2.88% | 66.67% | 0.37% | 27.64% |
| *Clostridium SS1 chauvoei ASV68* | | | | | | | | 11.82% | 66.67% | | |
| *Clostridium SS1 septicum ASV72* | | | | | | | | 10.72% | 66.67% | | |
| *Romboutsia; s_ASV4* | | 0.69% | 95.45% | 0.39% | 84.62% | | 76.92% | | | 26.69% | 100.00% |
| *Romboutsia; s_ASV18* | | | 45.45% | | 46.15% | | 50.00% | | | 3.87% | 87.64% |
| *Photobacterium; s_ASV23* | | 0.16% | 27.27% | | 7.69% | | 3.85% | 0.58% | 33.33% | 1.97% | 66.55% |
| *Tyzzerella; s_ASV22* | | | 13.64% | 0.82% | 30.77% | 0.52% | 15.38% | | | 1.60% | 52.00% |
| *Turicibacter; s_ASV32* | | | 36.36% | | 7.69% | | 11.54% | | | 1.91% | 70.91% |
| *Escherichia-Shigella; s_ASV19* | | | 13.64% | | 7.69% | | 15.38% | | 100.00% | 1.49% | 92.00% |
| *Fusobacterium; s_ASV10* | | 6.38% | 40.91% | | 7.69% | | 15.38% | | 66.67% | 2.56% | 27.27% |
| *Fusobacterium necrogenes ASV11* | | 1.52% | 31.82% | 0.15% | 23.08% | | 30.77% | | 33.33% | 2.69% | 34.55% |
| *Fusobacterium; s_ASV14* | | 1.36% | 27.27% | | | | 26.92% | | 33.33% | 1.71% | 39.64% |
| *Fusobacterium; s_ASV39* | | 0.23% | 13.64% | | | | | | | 0.38% | 21.82% |
| *Clostridium SS1; s_ASV34* | | 1.52% | 45.45% | | 15.38% | 0.55% | 53.85% | | | | |
| *Bacteroides; s_ASV71* | | 1.06% | 9.09% | | 7.69% | | | | | | |
| *Paeniclostridium; s_ASV183* | | 0.55% | 9.09% | | 7.69% | | 3.85% | | | | |
| *Paeniclostridium; s_ASV44* | | 0.52% | 31.82% | 1.58% | 46.15% | 0.11% | 7.69% | | | | 1.82% |
| *Paeniclostridium; s_ASV40* | | 0.44% | 27.27% | 1.87% | 46.15% | 0.14% | 7.69% | | | | |
| *Bacteroides; s_ASV27* | | 0.39% | 36.36% | | 7.69% | | 3.85% | 0.11% | 33.33% | 0.59% | 28.73% |
| *Mycoplasma; s_ASV80* | | 0.32% | 31.82% | 0.23% | 53.85% | 0.34% | 65.38% | | | | |
| *Photobacterium damselae ASV54* | | 0.26% | 22.73% | | | | | | | 0.64% | 31.27% |
| *Clostridium SS1; s_ASV28* | | 0.26% | 81.82% | | 53.85% | 0.79% | 42.31% | | 33.33% | | 15.64% |
| *Ureaplasma; s_ASV46* | | 0.20% | 31.82% | 0.38% | 38.46% | 0.28% | 61.54% | | | 0.11% | 44.73% |
| *Mycoplasmataceae; g_ASV90* | | 0.20% | 45.45% | 0.19% | 53.85% | 0.65% | 73.08% | | | | |
| *Paeniclostridium; s_ASV48* | | 0.20% | 9.09% | | | 0.42% | 23.08% | | | 0.19% | 21.09% |
| *Paeniclostridium; s_ASV282* | | 0.18% | 9.09% | 0.35% | 7.69% | | | | | | |
| *Mycoplasma; s_ASV103* | | 0.18% | 45.45% | | 53.85% | 0.37% | 65.38% | | | | |
| *Cetobacterium; s_ASV82* | | 0.14% | 13.64% | 2.04% | 30.77% | 0.79% | 30.77% | | | | 9.82% |
| *Clostridium SS1; s_ASV271* | | 0.13% | 13.64% | | | | 3.85% | | | | |
| *Romboutsia; s_ASV301* | | 0.12% | 9.09% | | | | | 3.86% | 33.33% | | 5.09% |
| *Mycoplasma; s_ASV16* | | | | 2.67% | 7.69% | 6.26% | 46.15% | | | | 4.73% |
| *Ureaplasma; s_ASV136* | | | 31.82% | 0.11% | 30.77% | 0.51% | 53.85% | | | | 7.64% |
| *Ureaplasma; s_ASV133* | | | 40.91% | | 30.77% | 0.62% | 61.54% | | | | |
| *Ligilactobacillus; s_ASV232* | | | 4.55% | | | 0.66% | 65.38% | | | | 0.36% |
| *Tyzzerella; s_ASV62* | | | 9.09% | | 7.69% | 0.13% | 11.54% | | | 0.67% | 24.36% |
| *Pseudoalteromonas; s_ASV31* | | | 18.18% | | | 2.74% | 7.69% | | 33.33% | | 20.73% |
| *Plesiomonas; s_ASV13* | | | 9.09% | | | 1.54% | 15.38% | | 33.33% | 0.19% | 25.45% |
| *Actinobacillus; s_ASV87* | | | 13.64% | | | | 11.54% | 3.02% | 33.33% | | |
| *Paeniclostridium; s_ASV111* | | | 4.55% | | | 0.12% | 11.54% | 0.44% | 33.33% | | 8.36% |
| *Photobacterium; s_ASV37* | | | 4.55% | | | | | | | 1.19% | 36.36% |
| *Psychrobacter; s_ASV146* | | | 4.55% | | | | 3.85% | 0.61% | 66.67% | | 0.73% |
| *Mycobacterium; s_ASV184* | | | 9.09% | | 7.69% | 1.76% | 23.08% | | | | |
| *Bacteroides; s_ASV36* | | | 4.55% | | | | | | | 0.65% | 17.09% |
| *Candidatus Arthromitus; s_ASV170* | | | 4.55% | 0.57% | 15.38% | | 11.54% | | | | 2.18% |
| *Cetobacterium; s_ASV314* | | | | 0.69% | 23.08% | | | | | | |
| *Paeniclostridium; s_ASV100* | | | | 0.50% | 15.38% | | | 0.19% | 33.33% | | |
| *Cetobacterium; s_ASV21* | | | | | 7.69% | | 7.69% | | 33.33% | 2.76% | 26.18% |
| *Ureaplasma; s_ASV208* | | | | | | 0.62% | 26.92% | | | | |
| *Mycoplasma; s_ASV216* | | | | | | 0.51% | 61.54% | | | | |
| *Fusobacterium; s_ASV222* | | | | | | 0.51% | 23.08% | | | | |
| *Leucothrix; s_ASV351* | | | | | | | | 2.49% | 33.33% | | |
| *Algitalea; s_ASV586* | | | | | | | | 1.09% | 33.33% | | |
| *Clostridium SS1; s_ASV390* | | | | | | | | 0.98% | 66.67% | | |
| *Ureaplasma; s_ASV254* | | | | | | | | 0.82% | 66.67% | | |
| *Leuconostoc lactis ASV15* | | | | | | | | 0.52% | 100.00% | | |
| *Vibrio; s_ASV42* | | | | | | | | | 33.33% | 1.14% | 25.45% |
| *Fusobacterium; s_ASV53* | | | | | | | | | | 0.54% | 17.09% |
| *Clostridium SS4; s_ASV252* | | | | | 15.38% | 0.18% | 23.08% | | | | |
| *Helicobacter; s_ASV365* | | | | | 23.08% | 0.41% | 46.15% | | | | |

**Supplementary Table 3.** ***Paeniclostridium*** **ASV1 was the dominant taxon in the fecal microbiotas of the wild resident killer whale populations.** The 'dominant_taxa' function (microbiomeutilities) was used to identify the most abundant taxon in fecal samples of the wild resident killer whales collected from 2017–2019 and calculate the number (n) and relative frequency of samples in each population that were dominated by each taxon. The top section shows the results for all samples during that time period, and the bottom shows the results for the dataset rarefied to 39,824 reads/sample and 13 samples/individual killer whale. Mean #ASVs/KW indicates the number of unique ASVs identified in each individual killer whale averaged across the population.

**All 2017-2019 samples**

| Population | Dominant ASV | n | Relative frequency |
|---|---|---|---|
| **Alaska RKWs** (n=26) richness= 1,135 ASVs mean # ASVs/KW= 95 | **Paeniclostridium; s_ ASV1** | **11** | **42%** |
| | Cetobacterium; s_ ASV6 | 3 | 12% |
| | Bifidobacteriaceae; g_ ASV106 | 2 | 8% |
| | Mycoplasma; s_ ASV16 | 2 | 8% |
| | Plesiomonas; s_ ASV13 | 1 | 4% |
| | Ureaplasma; s_ ASV17 | 1 | 4% |
| | Mycobacterium; s_ ASV184 | 1 | 4% |
| | Cetobacterium; s_ ASV20 | 1 | 4% |
| | Mycoplasma; s_ ASV24 | 1 | 4% |
| | Actinobacillus delphinicola ASV3 | 1 | 4% |
| | Pseudoalteromonas; s_ ASV31 | 1 | 4% |
| | Clostridium SS1; s_ ASV7 | 1 | 4% |
| **Northern RKWs** (n=13) richness= 161 ASVs mean #ASVs/KW= 31 | **Paeniclostridium; s_ ASV1** | **8** | **62%** |
| | Cetobacterium; s_ ASV6 | 2 | 15% |
| | Cetobacterium; s_ ASV20 | 1 | 8% |
| | Mycoplasma; s_ ASV24 | 1 | 8% |
| | Clostridium SS1; s_ ASV7 | 1 | 8% |
| **Southern RKWs** (n=21) richness= 497 ASVs mean #ASVs/KW= 46 | **Paeniclostridium; s_ ASV1** | **8** | **41%** |
| | Actinobacillus delphinicola ASV3 | 4 | 18% |
| | Edwardsiella tarda ASV5 | 3 | 14% |
| | Fusobacterium; s_ ASV10 | 2 | 9% |
| | Cetobacterium; s_ ASV6 | 2 | 9% |
| | Bacteroides; s_ ASV71 | 1 | 5% |
| | Fusobacterium; s_ ASV8 | 1 | 5% |

**Rarified & normalized** (39,824 reads/sample, 13 samples/population)

| Population | Dominant ASV | n | Relative frequency |
|---|---|---|---|
| **Alaska RKWs** (n=13) richness= 605 ASVs mean # ASVs/KW= 88 | **Paeniclostridium; s_ ASV1** | **5** | **38%** |
| | Bifidobacteriaceae; g_ ASV106 | 2 | 15% |
| | Cetobacterium; s_ ASV6 | 2 | 15% |
| | Plesiomonas; s_ ASV13 | 1 | 8% |
| | Ureaplasma; s_ ASV17 | 1 | 8% |
| | Mycoplasma; s_ ASV24 | 1 | 8% |
| | Pseudoalteromonas; s_ ASV31 | 1 | 8% |
| **Northern RKWs** (n=13) richness= 157 ASVs mean #ASVs/KW= 30 | **Paeniclostridium; s_ ASV1** | **8** | **62%** |
| | Cetobacterium; s_ ASV6 | 2 | 15% |
| | Cetobacterium; s_ ASV20 | 1 | 8% |
| | Mycoplasma; s_ ASV24 | 1 | 8% |
| | Clostridium SS1; s_ ASV7 | 1 | 8% |
| **Southern RKWs** (n=13) richness= 167 ASVs mean #ASVs/KW= 32 | **Paeniclostridium; s_ ASV1** | **5** | **38%** |
| | Actinobacillus delphinicola ASV3 | 3 | 23% |
| | Edwardsiella tarda ASV5 | 2 | 15% |
| | Cetobacterium; s_ ASV6 | 2 | 15% |
| | Bacteroides; s_ ASV71 | 1 | 8% |

**Supplementary Table 4. Resident and Aquarium-housed killer whale population ASV differential abundance analysis.** Amplicon sequence variant (ASV) differential abundance (DA) in fecal samples collected from the Southern (19 individuals; 22 samples), Northern (9 individuals, 13 samples), and Alaska (21 individuals, 26 samples) Resident killer whale populations in 2017–2019. Abundant ASVs (>1000 reads in ≥2 samples) were analyzed (Fig. 2b, Supplementary Fig. 7). General and linear mixed models ('MaAsLin2') (M2) identified ASVs enriched in each population. To identify robust population indicators, 3 additional consensus DA methods were utilized: treeDA (tDA), ALDEx2, and coda4microbiome (CODA). Effect sizes are shown for ASVs enriched for each population (M2 coef>0) vs both others via ≥1 DA method. "Consensus" indicates the number of positive DA tests. "DA rank" shows ASVs with the greatest consensus and effect (1 = most differential). ASVs differential for NRKW (vs both others) and ARKW (vs both others), or for both relatively fit populations ("Fit Pops": NRKW & ARKW) vs SRKW (effect <0) are noted. ASV prevalence and mean RA is shown for each population. The aquarium-housed KW (AqKW) ASV differential abundance analysis compared ASV RA in 60 samples from 12 AqKWs collected in 2019 to ASV RAs in the wild KWs combined (n=60).

**RESIDENT KILLER WHALES**

| | | | | | | | | | | | Relative Abundance | | | Prevalence | | |
|---|---|---|---|---|---|---|---|---|---|---|---|---|---|---|---|---|
| **ASV taxonomy** | **Enriched population** (vs both others) | **Enriched both fit** (DA tests) | **ASV unique to population(s)** | **M2 padj** | **M2 coef** | **treeDA** | **CODA** | **ALDEx2** | **Consensus** | **DA rank** | SRKW | NRKW | ARKW | SRKW | NRKW | ARKW |
| Edwardsiella tarda ASV5 | | | | 3.669E-03 | 3.86 | 0.04 | 0.23 | 0.83 | 4 | 1 | 9.56% | 0.89% | 2.45% | 95.45% | 84.62% | 96.15% |
| Photobacterium; s_ASV2 | | | | 2.226E-03 | 2.89 | 0.05 | 0.31 | 0.88 | 4 | 2 | 1.83% | 0.10% | 0.69% | 100.00% | 92.31% | 92.31% |
| Fusobacterium; s_ASV8 | | | | 2.853E-02 | 2.76 | 0.06 | 0.17 | | 3 | 3 | 5.41% | 0.01% | 0.66% | 59.09% | 15.38% | 46.15% |
| Mycoplasma; s_ASV132 | | | Unique to SRKW | 4.970E-02 | 0.35 | 0.20 | 0.29 | | 3 | 4 | 0.20% | | | 18.18% | | |
| Fusobacterium; s_ASV10 | | | | 2.637E-10 | 6.06 | 0.04 | | | 2 | 5 | 6.38% | 0.01% | 0.01% | 40.91% | 7.69% | 15.38% |
| Paeniclostridium; s_ASV183 | | | | 2.034E-03 | 4.95 | 0.02 | | | 2 | 6 | 0.55% | 0.03% | 0.01% | 9.09% | 7.69% | 3.85% |
| Clostridium SS1; s_ASV271 | | | | 0.000E+00 | 2.09 | 0.02 | | | 2 | 7 | 0.13% | | 0.08% | 13.64% | | 3.85% |
| Photobacterium damselae ASV54 | **Southern Resident killer whales:** (SR vs NR & AR, coef>0) | | Unique to SRKW | 3.330E-02 | 0.65 | 0.03 | | | 2 | 8 | 0.26% | | | 22.73% | | |
| Fusobacterium; s_ASV14 | | | | 1.182E-03 | 4.47 | | | | 1 | 9 | 1.36% | | 0.03% | 27.27% | | 26.92% |
| Fusobacterium; s_ASV39 | | | Unique to SRKW | 1.353E-02 | 6.06 | | | | 1 | 10 | 0.23% | | | 13.64% | | |
| Fusobacterium necrogenes ASV11 | | | | 3.005E-02 | 2.94 | | | | 1 | 11 | 1.52% | 0.15% | 0.03% | 31.82% | 23.08% | 30.77% |
| Photobacterium; s_ASV23 | | | | 0.000E+00 | 2.33 | | | | 1 | 12 | 0.16% | 0.00% | 0.06% | 27.27% | 7.69% | 3.85% |
| Actinobacillus delphinicola ASV3 | | | | 4.703E-02 | 0.88 | | | | 1 | 13 | 12.90% | 4.80% | 5.02% | 95.45% | 92.31% | 100.00% |
| Clostridium SS1; s_ASV7 | | | | | | 0.02 | | | 1 | 14 | 4.02% | 5.00% | 1.54% | 77.27% | 53.85% | 84.62% |
| Romboutsia; s_ASV4 | | | | | | | | 0.27 | 1 | 15 | 0.69% | 0.39% | 0.08% | 95.45% | 84.62% | 76.92% |
| Clostridium_SS1; s_ASV34 | | | | | | 0.03 | | | 1 | 16 | 1.52% | 0.07% | 0.55% | 45.45% | 15.38% | 53.85% |
| Clostridium_SS1; s_ASV28 | | | | | | 0.02 | | | 1 | 17 | 0.26% | 0.06% | 0.79% | 81.82% | 53.85% | 42.31% |
| Cetobacterium; s_ASV82 | | "Fit Pops" vs SR (M2, tDA, CODA) | | 4.597E-02 | -3.36 | -0.02 | -0.11 | | 3 | 1 | 0.14% | 2.04% | 0.79% | 13.64% | 30.77% | 30.77% |
| Clostridium SS4; s_ASV252 | **Both relatively fit populations** (SR vs NR & AR, coef<0) | "Fit Pops" vs SR (M2, tDA, CODA) | Unique to "Fit Pops" | 2.999E-228 | -2.42 | -0.04 | | | 3 | 2 | | 0.03% | 0.18% | | 15.38% | 23.08% |
| Ureaplasma; s_ASV17 | | "Fit Pops" vs SR (M2) | | 1.353E-02 | -1.49 | | | | 1 | 3 | 1.61% | 2.46% | 3.17% | 59.09% | 100.00% | 73.08% |
| Paeniclostridium; s_ASV40 | | | | 0.000E+00 | 4.62 | 0.04 | 0.18 | | 3 | 1 | 0.44% | 1.87% | 0.14% | 27.27% | 46.15% | 7.69% |
| Actinobacillus; s_ASV66 | | | | 1.655E-03 | 3.20 | 0.05 | 0.23 | | 3 | 2 | 0.38% | 3.64% | 0.07% | 45.45% | 76.92% | 57.69% |
| Mycoplasma; s_ASV24 | | | | 1.330E-03 | 2.23 | 0.05 | 0.18 | | 3 | 3 | 0.89% | 5.44% | 2.07% | 45.45% | 92.31% | 57.69% |
| Cetobacterium; s_ASV314 | | | Unique to NR | 1.789E-02 | 0.83 | 0.14 | 0.27 | | 3 | 4 | | 0.69% | | | 23.08% | |
| Paeniclostridium; s_ASV44 | **Northern Resident killer whales:** (NR vs SR & AR, coef>0) | | | 0.000E+00 | 4.50 | 0.03 | | | 2 | 5 | 0.52% | 1.58% | 0.11% | 31.82% | 46.15% | 7.69% |
| Actinobacillus; s_ASV800 | | | Unique to NR | 7.165E-03 | 1.82 | 0.05 | | | 2 | 6 | | 0.32% | | | 30.77% | |
| Paeniclostridium; s_ASV1 | | | | 5.799E-07 | 2.23 | | | | 1 | 7 | 29.98% | 45.96% | 29.22% | 100.00% | 100.00% | 100.00% |
| Tyzzerella; s_ASV22 | | | | 0.000E+00 | 2.10 | | | | 1 | 8 | 0.02% | 0.82% | 0.52% | 13.64% | 30.77% | 15.38% |
| Paeniclostridium; s_ASV440 | | | Unique to NR | 4.338E-02 | 0.01 | | | | 1 | 9 | | 0.49% | | | 15.38% | |
| Actinobacillus; s_ASV1033 | | | Unique to NR | 9.828E-03 | 0.00 | | | | 1 | 10 | | 0.23% | | | 15.38% | |
| Paeniclostridium; s_ASV282 | | | | 0.000E+00 | | 0.03 | | | 1 | 11 | 0.18% | 0.35% | | 9.09% | 7.69% | |
| Ligilactobacillus; s_ASV232 | | | | 2.326E-05 | 7.75 | 0.15 | 0.65 | 0.84 | 4 | 1 | 0.00% | | 0.66% | 4.55% | | 65.38% |
| Mycoplasma; s_ASV216 | | | Unique to AR | 1.250E-06 | 3.89 | 0.06 | 0.35 | 0.80 | 4 | 2 | | | 0.51% | | | 61.54% |
| Ligilactobacillus; s_ASV69 | | | | 3.192E-09 | 4.30 | 0.04 | 0.11 | | 3 | 3 | 0.02% | 0.01% | 1.76% | 36.36% | 46.15% | 65.38% |
| Bifidobacteriaceae; g_ASV106 | | | | 1.581E-12 | 7.44 | | | 0.58 | 2 | 4 | 0.00% | | 2.57% | 9.09% | | 57.69% |
| Helicobacter; s_ASV365 | | "Fit Pops" vs SR (M2, CODA) | Unique to "Fit Pops" | 1.317E-03 | 4.02 | | -0.16 | | 2 | 5 | | 0.03% | 0.41% | | 23.08% | 46.15% |
| Ligilactobacillus; s_ASV789 | | | | 1.603E-02 | 1.70 | 0.04 | | | 2 | 6 | | | 0.13% | | | 26.92% |
| Tyzzerella; s_ASV531 | | | | 0.000E+00 | 0.92 | 0.02 | | | 2 | 7 | 0.13% | | 0.17% | 4.55% | | 7.69% |
| Paeniclostridium; s_ASV328 | | | Unique to AR | 4.010E-02 | 0.88 | 0.10 | | | 2 | 8 | | | 0.23% | | | 19.23% |
| Paeniclostridium; s_ASV111 | | | | 2.199E-263 | 5.46 | | | | 1 | 9 | 0.01% | | 0.12% | 4.55% | | 11.54% |
| Plesiomonas; s_ASV13 | **Alaska Resident killer whales:** (AR vs SR & NR, coef>0) | | | 0.000E+00 | 3.71 | | | | 1 | 10 | 0.03% | | 1.54% | 9.09% | | 15.38% |
| Mycoplasma; s_ASV122 | | | | 0.000E+00 | 3.43 | | | | 1 | 11 | 0.06% | 0.23% | 1.30% | 22.73% | 23.08% | 53.85% |
| Mycoplasma; s_ASV16 | | AR vs others, NR vs others & "Fit Pops" vs SR (M2) | Unique to "Fit Pops" | 4.970E-02 | -2.87 | | | | 1 | 12 | | 2.67% | 6.26% | | 7.69% | 46.15% |
| Ureaplasma; s_ASV133 | | | | 4.421E-04 | 2.19 | | | | 1 | 13 | 0.10% | 0.05% | 0.62% | 40.91% | 30.77% | 61.54% |
| Ureaplasma; s_ASV33 | | | | 4.703E-02 | 1.90 | | | | 1 | 14 | 0.29% | 0.51% | 1.00% | 50.00% | 46.15% | 61.54% |
| Negativicoccus; s_ASV303 | | | Unique to AR | 2.078E-02 | 1.77 | | | | 1 | 15 | | | 0.48% | | | 26.92% |
| Mycoplasmataceae; g_ASV90 | | | | 4.060E-02 | 1.69 | | | | 1 | 16 | 0.20% | 0.19% | 0.65% | 45.45% | 53.85% | 73.08% |
| Fusobacterium; s_ASV594 | | | Unique to AR | 2.928E-02 | 1.42 | | | | 1 | 17 | | | 0.13% | | | 23.08% |
| Ureaplasma; s_ASV208 | | | Unique to AR | 1.603E-02 | 1.22 | | | | 1 | 18 | | | 0.62% | | | 26.92% |
| Fusobacterium; s_ASV222 | | | Unique to AR | 3.026E-02 | 1.06 | | | | 1 | 19 | | | 0.51% | | | 23.08% |
| Paeniclostridium; s_ASV48 | | | | 2.046E-168 | 0.55 | | | | 1 | 20 | 0.20% | | 0.42% | 9.09% | | 23.08% |
| Actinobacillus; s_ASV162 | | | | | | | | | | | 0.34% | | 0.03% | 13.64% | | 15.38% |
| Tyzzerella; s_ASV62 | | | | | | | | | | | 0.04% | 0.08% | 0.13% | 9.09% | 7.69% | 11.54% |
| Cetobacterium; s_ASV6 | | | | | | | | | | | 8.78% | 9.98% | 13.16% | 95.45% | 84.62% | 96.15% |
| Cetobacterium; s_ASV20 | | | | | | | | | | | 1.24% | 4.69% | 3.74% | 68.18% | 53.85% | 69.23% |
| Actinobacillus; s_ASV25 | | | | | | | | | | | 0.80% | 0.16% | 1.04% | 40.91% | 46.15% | 61.54% |
| Mycoplasma; s_ASV80 | | | | | | | | | | | 0.32% | 0.23% | 0.34% | 31.82% | 53.85% | 65.38% |
| Ureaplasma; s_ASV46 | | | | | | | | | | | 0.20% | 0.38% | 0.28% | 31.82% | 38.46% | 61.54% |
| Mycoplasma; s_ASV103 | | | | | | | | | | | 0.18% | 0.08% | 0.37% | 45.45% | 53.85% | 65.38% |

**AQUARIUM-HOUSED KILLER WHALES**

| | | | | | | | | | | Relative Abundance | | Prevalence | |
|---|---|---|---|---|---|---|---|---|---|---|---|---|---|
| **ASV taxonomy** | **Enriched population** | **ASV unique to one group** | **M2 padj** | **M2 coef** | **treeDA** | **CODA** | **ALDEx2** | **Consensus** | **DA rank** | AqKW | Wild | AqKW | Wild |
| Escherichia-Shigella; s_ASV19 | | | 3.797E-47 | -6.71427 | -0.10327 | -1 | -1.89662 | 4 | 1 | 1.49% | 0.00% | 92.00% | 12.24% |
| Romboutsia; s_ASV4 | | | 3.679E-28 | -4.11559 | -0.04268 | | -1.43855 | 3 | 2 | 26.69% | 0.38% | 100.00% | 85.66% |
| Clostridium SS1; s_ASV30 | | | 2.293E-03 | -2.54425 | | | -0.73384 | 2 | 3 | 0.28% | 0.03% | 53.82% | 9.91% |
| Phocoenobacter; s_ASV147 | | | 2.374E-05 | -2.96086 | | | -0.7252 | 2 | 4 | 0.19% | 0.00% | 57.45% | 16.67% |
| Fusobacterium; s_ASV39 | | | 0.000E+00 | -3.16292 | | | -0.19059 | 2 | 5 | 0.38% | 0.08% | 21.82% | 4.55% |
| Photobacterium; s_ASV2 | | | 4.154E-26 | -3.47797 | | | -1.13143 | 2 | 6 | 25.74% | 0.87% | 98.18% | 94.87% |
| Photobacterium; s_ASV23 | | | 8.912E-07 | -3.63251 | | | -0.95923 | 2 | 7 | 1.97% | 0.07% | 66.55% | 12.94% |
| Turicibacter; s_ASV32 | | | 5.764E-10 | -4.11486 | | | -0.85892 | 2 | 8 | 1.91% | 0.00% | 70.91% | 18.53% |
| Romboutsia; s_ASV18 | | | 4.037E-16 | -4.13373 | | | -0.82836 | 2 | 9 | 3.87% | 0.02% | 87.64% | 47.20% |
| Turicibacter; s_ASV84 | | | 2.749E-09 | -5.55717 | | | -0.60416 | 2 | 10 | 0.41% | 0.00% | 37.82% | 5.59% |
| Photobacterium; s_ASV37 | **Aquarium-housed killer whales** (vs wild resident populations) | | 2.583E-06 | -5.96139 | | | -0.28809 | 2 | 11 | 1.19% | 0.00% | 36.36% | 1.52% |
| Cetobacterium; s_ASV21 | | | 1.079E-12 | -8.54376 | | | -0.24163 | 2 | 12 | 2.77% | 0.00% | 26.18% | 5.13% |
| Photobacterium damselae ASV54 | | | 0.000E+00 | -0.91829 | | | | 1 | 13 | 0.64% | 0.09% | 31.27% | 7.58% |
| Fusobacterium necrogenes ASV11 | | | 3.250E-02 | -2.34344 | | | | 1 | 14 | 2.69% | 0.57% | 34.55% | 28.55% |
| Bacteroides; s_ASV36 | | | 6.901E-11 | -11.0059 | | | | 1 | 15 | 0.65% | 0.00% | 17.09% | 1.52% |
| Fusobacterium; s_ASV53 | | Unique to AqKW | | | | | -0.1852 | 1 | | 0.54% | | 17.09% | |
| Fusobacterium; s_ASV14 | | | | | | | -0.25266 | 1 | | 1.71% | 0.46% | 39.64% | 18.07% |
| Vibrio; s_ASV42 | | Unique to AqKW | | | | | -0.3484 | 1 | | 1.14% | | 25.45% | |
| Romboutsia; s_ASV41 | | | | | | | -0.41366 | 1 | | 0.30% | 0.03% | 36.36% | 12.47% |
| Oscillospira; s_ASV142 | | Unique to AqKW | | | | | -0.43037 | 1 | | 0.24% | | 32.00% | |
| Tyzzerella; s_ASV22 | | | | | | | -0.6811 | 1 | | 1.60% | 0.45% | 52.00% | 19.93% |
| Actinobacillus delphinicola ASV3 | | | 2.687E-27 | 4.867542 | 0.020848 | 0.11 | 1.022239 | 4 | 1 | 0.55% | 7.57% | 62.91% | 95.92% |
| Paeniclostridium; s_ASV1 | | | 2.110E-29 | 4.095187 | 0.042133 | 0.84 | 0.90376 | 4 | 2 | 3.38% | 35.05% | 91.64% | 100.00% |
| Edwardsiella tarda ASV5 | | | 2.223E-11 | 3.957393 | 0.041356 | | 1.396703 | 3 | 3 | 0.14% | 4.30% | 21.82% | 92.07% |
| Mycoplasma; s_ASV122 | | | 3.393E-12 | 7.454556 | | | 0.416463 | 2 | 4 | 0.00% | 0.53% | 5.45% | 33.22% |
| Mycoplasma; s_ASV16 | | | 1.695E-06 | 8.035287 | | | | 1 | 5 | 0.01% | 2.98% | 4.73% | 17.95% |
| Paeniclostridium; s_ASV44 | | | 2.193E-06 | 7.341664 | | | | 1 | 6 | 0.00% | 0.74% | 1.82% | 28.55% |
| Clostridium SS1; s_ASV7 | | | 2.514E-07 | 2.243763 | | | | 1 | 7 | 1.19% | 3.52% | 93.45% | 71.91% |
| Cetobacterium; s_ASV82 | | | 0.000E+00 | 1.533567 | | | | 1 | 8 | 0.04% | 0.99% | 9.82% | 25.06% |
| Actinobacillus; s_ASV25 | **Wild resident killer whale populations** (vs AqKWs) | | 3.602E-02 | 1.444235 | | | | 1 | 9 | 0.29% | 0.67% | 48.73% | 49.53% |
| Ureaplasma; s_ASV17 | | | | | | | 1.11996 | 1 | | 0.00% | 2.41% | 0.36% | 77.39% |
| Mycoplasma; s_ASV24 | | | | | | | 0.860631 | 1 | | 0.00% | 2.80% | 0.36% | 65.15% |
| Mycoplasmataceae; g_ASV90 | | Unique to wild | | | | | 0.824737 | 1 | | | 0.35% | | 57.46% |
| Ureaplasma; s_ASV33 | | Unique to wild | | | | | 0.755341 | 1 | | | 0.60% | | 52.56% |
| Actinobacillus; s_ASV66 | | Unique to wild | | | | | 0.753379 | 1 | | | 1.36% | | 60.02% |
| Mycoplasma; s_ASV80 | | Unique to wild | | | | | 0.68705 | 1 | | | 0.30% | | 50.35% |
| Ureaplasma; s_ASV133 | | Unique to wild | | | | | 0.612596 | 1 | | | 0.26% | | 44.41% |
| Ligilactobacillus; s_ASV69 | | | | | | | 0.585079 | 1 | | 0.00% | 0.60% | 0.36% | 49.30% |
| Clostridium SS1; s_ASV34 | | Unique to wild | | | | | 0.546618 | 1 | | | 0.72% | | 38.23% |
| Cetobacterium; s_ASV6 | | | | | | | | 0 | | 3.07% | 10.64% | 88.36% | 92.07% |
| Fusobacterium; s_ASV10 | | | | | | | | 0 | | 2.56% | 2.13% | 27.27% | 21.33% |
| Cetobacterium; s_ASV20 | | | | | | | | 0 | | 0.62% | 3.22% | 47.27% | 63.75% |
| Fusobacterium; s_ASV8 | | | | | | | | 0 | | 0.91% | 2.03% | 25.09% | 40.21% |
| Bacteroides; s_ASV27 | | | | | | | | 0 | | 0.59% | 0.13% | 28.73% | 15.97% |
| Paeniclostridium; s_ASV40 | | Unique to wild | | | | | | 0 | | | 0.82% | | 27.04% |
| Paeniclostridium; s_ASV48 | | | | | | | | 0 | | 0.19% | 0.21% | 21.09% | 10.72% |

**Supplementary Table 5. The ARKW population had the highest abundance and diversity of lactic acid-producing bacteria.** The distribution of relative abundances (RAs) of putative lactic acid-producing bacteria in the wild resident KW dataset was examined. Fecal samples were restricted to years when multiple resident populations were sampled (2017–2019). Read counts were proportion transformed. The data were restricted to members of *Lactobacillus* , *Lactococcus* , *Leuconostoc* , *Pediococcus* , *Streptococcus* , *Aerococcus* , *Alloiococcus* , *Carnobacterium* , *Dolosigranulum* , *Enterococcus* , *Oenococcus* , *Tetragenococcus* , *Vagococcus* , *Weissella* , *Sporolactobacillus* , *Ligilactobacillus* , and *Bifidobacterium* . Only ASVs with RA ≥0.01% are displayed.

| Population | ASV | Taxonomy | Mean RA | Prevalence | Top BLAST hit (Per.ID%) | Source | Accession # | Closest classified BLAST hit (or additional match) |
|---|---|---|---|---|---|---|---|---|
| ARKW | ASV106 | Bifidobacteriaceae; g_ | 2.57% | 57.69% | unclassified (99.15%) | Bottlenose dolphin (rectal & gastric) | JQ201016.1 | *Bifidobacterium coryneforme* (94.44%) |
| ARKW | ASV280 | Bifidobacteriaceae; g_ | 0.30% | 11.54% | unclassified (99.57%) | Bottlenose dolphin (rectal & gastric) | JQ201016.1 | *Bifidobacterium coryneforme* (94.87%) |
| ARKW | ASV698 | Bifidobacteriaceae; g_ | 0.06% | 11.54% | unclassified (98.72%) | Bottlenose dolphin (rectal & gastric) | JQ201016.1 | *Bifidobacterium coryneforme* (94.87%) |
| ARKW | ASV232 | *Ligilactobacillus* ; s_ | 0.66% | 65.38% | unclassified (100.00%) | Bottlenose dolphin (rectal & gastric) | JQ201990.1 | |
| ARKW | ASV789 | *Ligilactobacillus* ; s_ | 0.13% | 26.92% | *Lactobacillus sp. MMP242* (100.00%) | Bottlenose dolphin (gastric) | JX142131.1 | |
| ARKW | ASV406 | *Ligilactobacillus* ; s_ | 0.10% | 34.62% | unclassified (100.00%) | Bottlenose dolphin (gastric) | JQ193526.1 | *Ligilactobacillus salivarius* (97.45%) |
| ARKW | ASV942 | *Ligilactobacillus* ; s_ | 0.07% | 15.38% | unclassified (100.00%) | Bottlenose dolphin (gastric) | JQ193526.1 | *Ligilactobacillus salivarius* (97.02%) |
| ARKW | ASV1463 | *Ligilactobacillus* ; s_ | 0.06% | 11.54% | unclassified (100.00%) | Bottlenose dolphin (rectal & gastric) | JQ194436.1 | *Lactobacillus sp. MMP242* (100.00%) |
| ARKW | ASV1260 | *Ligilactobacillus* ; s_ | 0.03% | 15.38% | *Lactobacillus sp. MMP242* (100.00%) | Bottlenose dolphin (gastric) | JX142131.1 | *Bifidobacterium longum* (100.00%) |
| ARKW | ASV1376 | *Ligilactobacillus* ; s_ | 0.02% | 11.54% | unclassified (100.00%) | Bottlenose dolphin (gastric) | JQ193526.1 | *Ligilactobacillus salivarius* (100.00%) |
| ARKW | | | 1.75% | 65.38% | | | | |
| NRKW | ASV69 | *Ligilactobacillus* ; s_ | 0.01% | 46.15% | *Lactobacillus sp. MMP242* (100.00%) | Bottlenose dolphin (rectal & gastric) | JX142131.1 | |
| SRKW | | | 0.02% | 36.36% | | | | |

**Supplementary Table 6. Adonis2 results for SRKW and AqKW study covariates.** A multivariate analysis of variance (MANOVA, 'adonis2') was performed using clr-transformed Aitchison distances between the SRKW (left) and AqKW (right) fecal microbiotas to determine their contribution to the variation observed in the KW microbiotas. For the SRKW population, the analysis was repeated after shuffling the order of the covariates, to obtain the results for each covariate while incorporating the available health metadata.

| Southern Resident killer whale population (SRKW) | | | | | |
|---|---|---|---|---|---|
| adonis2(aitchison.matrix ~ ID* Sex * Pod * Matriline * Year * Age, data = df, perm = 999, by="terms") | | | | | |
| SRKW covariate | Df | SumOfSqs | R2 | F | Pr(>F) |
| Killer whale Identity (ID) | 52 | 119490 | 24.71% | 2.03 | 0.001 |
| Year | 1 | 17462 | 3.61% | 15.44 | 0.001 |
| Age | 1 | 2207 | 0.46% | 1.95 | 0.005 |
| ID: Year | 37 | 54304 | 11.23% | 1.30 | 0.001 |
| ID: Year: Age | 30 | 40072 | 8.29% | 1.18 | 0.022 |
| ID: Age | 23 | 27814 | 5.75% | 1.07 | 0.179 |
| ID: Matriline: Year: Age | 3 | 3380 | 0.70% | 1.00 | 0.617 |
| Year: Age | 1 | 3319 | 0.69% | 2.93 | 0.001 |
| ID: Pod: Matriline: Year: Age | 1 | 2439 | 0.50% | 2.16 | 0.022 |
| ID: Sex: Pod: Year: Age | 1 | 1691 | 0.35% | 1.50 | 0.207 |
| ID: Sex: Pod: Matriline: Year: Age | 1 | 1225 | 0.25% | 1.08 | 0.365 |
| ID: Sex: Matriline: Year: Age | 1 | 1205 | 0.25% | 1.07 | 0.296 |
| Matriline: Year: Age | 1 | 1148 | 0.24% | 1.02 | 0.305 |
| ID: Sex: Year: Age | 1 | 1034 | 0.21% | 0.91 | 0.251 |
| ID: Pod: Year: Age | 1 | 926 | 0.19% | 0.82 | 0.633 |
| Residual | 182 | 205876 | 42.57% | | |
| Total | 337 | 483594 | 100.00% | | |

| Aquarium-housed killer whales | | | | | |
|---|---|---|---|---|---|
| adonis2(aitchison.matrix ~ Facility * Sex * ID * Age * Month, data = df, perm = 999, by="terms") | | | | | |
| Aquarium KW covariate | Df | SumOfSqs | R2 | F | Pr(>F) |
| ID | 10 | 100001 | 33.86% | 14.67 | 0.001 |
| Facility | 1 | 15968 | 5.41% | 23.43 | 0.001 |
| Sex | 1 | 13926 | 4.71% | 20.43 | 0.001 |
| Month | 1 | 3012 | 1.02% | 4.42 | 0.001 |
| Age | 1 | 1310 | 0.44% | 1.92 | 0.001 |
| ID: Month | 10 | 10692 | 3.62% | 1.57 | 0.001 |
| ID: Age | 4 | 4570 | 1.55% | 1.68 | 0.001 |
| Facility: Month | 1 | 1910 | 0.65% | 2.80 | 0.001 |
| Facility: Age | 1 | 1505 | 0.51% | 2.21 | 0.001 |
| Sex: Month | 1 | 1284 | 0.43% | 1.88 | 0.001 |
| ID: Age: Month | 2 | 1473 | 0.50% | 1.08 | 0.264 |
| Sex: Age | 1 | 1011 | 0.34% | 1.48 | 0.057 |
| Facility: Age: Month | 1 | 619 | 0.21% | 0.91 | 0.259 |
| Age: Month | 1 | 623 | 0.21% | 0.91 | 0.282 |
| Residual | 231 | 157461 | 53.31% | | |
| Total | 263 | 295363 | 100.00% | | |

| SRKW covariates mixed order | | | | | |
|---|---|---|---|---|---|
| adonis2(aitchison.matrix ~ Year * Sex * Pod * Matriine * ID * Age, data = df, perm = 999, by="terms") | | | | | |
| SRKW covariate | Df | SumOfSqs | R2 | F | Pr(>F) |
| Killer whale Identity (ID) | 37 | 79858 | 16.51% | 1.89 | 0.001 |
| Matriline | 12 | 26086 | 5.39% | 1.90 | 0.001 |
| Year | 1 | 20332 | 4.20% | 17.81 | 0.001 |
| Pod | 2 | 7654 | 1.58% | 3.35 | 0.001 |
| Sex | 1 | 3022 | 0.62% | 2.65 | 0.001 |
| Age | 1 | 2207 | 0.46% | 1.93 | 0.002 |
| Year: ID | 23 | 32926 | 6.81% | 1.25 | 0.002 |
| Year: ID: Age | 17 | 23049 | 4.77% | 1.19 | 0.037 |
| Year: Matriline | 11 | 16797 | 3.47% | 1.34 | 0.003 |
| ID: Age | 12 | 14836 | 3.07% | 1.08 | 0.295 |
| Year: Matriline: Age | 11 | 13789 | 2.85% | 1.10 | 0.287 |
| Matriline: Age | 8 | 9410 | 1.95% | 1.03 | 0.224 |
| Year: Pod | 2 | 3292 | 0.68% | 1.44 | 0.034 |
| Year: Pod: Age | 2 | 3029 | 0.63% | 1.33 | 0.058 |
| Year: Age | 1 | 2941 | 0.61% | 2.58 | 0.002 |
| Pod: Age | 2 | 2396 | 0.50% | 1.05 | 0.229 |
| Sex: Age | 1 | 1550 | 0.32% | 1.36 | 0.074 |
| Year: Pod: Matriline: ID: Age | 1 | 1505 | 0.31% | 1.32 | 0.319 |
| Year: Sex: Pod: Matriline: ID: Age | 1 | 1493 | 0.31% | 1.31 | 0.102 |
| Year: Sex: ID: Age | 1 | 1392 | 0.29% | 1.22 | 0.219 |
| Year: Sex | 1 | 1289 | 0.27% | 1.13 | 0.288 |
| Year: Sex: Matriline: ID: Age | 1 | 1182 | 0.24% | 1.04 | 0.473 |
| Year: Sex: Age | 1 | 1099 | 0.23% | 0.96 | 0.615 |
| Year: Matriline: ID: Age | 1 | 917 | 0.19% | 0.80 | 0.678 |
| Year: Pod: ID: Age | 1 | 835 | 0.17% | 0.73 | 0.748 |
| Year: Sex: Pod: ID: Age | 1 | 659 | 0.14% | 0.58 | 0.988 |
| Residual | 184 | 210049 | 43.43% | | |
| Total | 337 | 483594 | 100.00% | | |

| SRKW with survival data (n=274) | | | | | |
|---|---|---|---|---|---|
| onis2(aitchison.matrix ~ Survival * Matriline * Sex * Pod * ID * Year * Age, data = df, permutations = 999, by="term |  |  |  |  |  |
| SRKW covariate | Df | SumOfSqs | R2 | F | Pr(>F) |
| ID | 33 | 64591 | 16.87% | 1.79 | 0.001 |
| Matriline | 14 | 31930 | 8.34% | 2.08 | 0.001 |
| Year | 1 | 11799 | 3.08% | 10.78 | 0.001 |
| Survival | 2 | 6584 | 1.72% | 3.01 | 0.001 |
| Sex | 1 | 2677 | 0.70% | 2.45 | 0.001 |
| Age | 1 | 2090 | 0.55% | 1.91 | 0.003 |
| ID: Year | 16 | 23068 | 6.03% | 1.32 | 0.009 |
| Matriline: Year | 13 | 19070 | 4.98% | 1.34 | 0.005 |
| Matriline: Year: Age | 13 | 17188 | 4.49% | 1.21 | 0.114 |
| ID: Year: Age | 10 | 12867 | 3.36% | 1.18 | 0.003 |
| ID: Age | 8 | 11728 | 3.06% | 1.34 | 0.022 |
| Matriline: Age | 10 | 10997 | 2.87% | 1.01 | 0.332 |
| Survival: Age | 2 | 2840 | 0.74% | 1.30 | 0.08 |
| Year: Age | 1 | 2234 | 0.58% | 2.04 | 0.013 |
| Survival: Matriline | 2 | 2205 | 0.58% | 1.01 | 0.172 |
| Survival: Year | 1 | 1676 | 0.44% | 1.53 | 0.069 |
| Sex: Year | 1 | 1547 | 0.40% | 1.41 | 0.174 |
| Sex: Age | 1 | 1320 | 0.34% | 1.21 | 0.238 |
| Survival: Year: Age | 1 | 1284 | 0.34% | 1.17 | 0.221 |
| Sex: Year: Age | 1 | 847 | 0.22% | 0.77 | 0.895 |
| Residual | 141 | 154262 | 40.30% | | |
| Total | 273 | 382803 | 100.00% | | |

**Supplementary Table 7. Threshold Indicator Taxa Analysis (TITAN2).** Threshold Indicator Taxa Analysis (TITAN2, v2.4.3, nBoot = 500) was performed on the SRKW fecal microbiotas to identify taxon-specific timepoints of maximum relative abundance (RA) and prevalence and community-level change points along two temporal gradients, (top) day of study and (bottom) host age (in years). Indicator species were classified based on their directionality around the change point as either "decreasers" (negative response, highlighted in gray) or "increasers" (positive response, highlighted in red). Only taxa with TITAN2 purity ≥95.0% and reliability scores ≥97.0% are displayed. The abundant taxa (RA >0.25%) with purity and reliability scores ≥97.0% are in bold text--these are included in **Fig. 4F** and further analyzed in **Supplementary Figs. 15-17**.

### DAY OF STUDY: (2005-2019)

| Group | Taxonomy | Age indicator | zenv.cp | freq | IndVal | zscore | 5% | 50% | 95% | purity | reliability | z.median | Avg. RA |
|---|---|---|---|---|---|---|---|---|---|---|---|---|---|
| DECREASERS | ***Photobacterium; s_ ASV2*** | | **2764.5** | **319** | **74.92** | **4** | **1445** | **2759** | **2856** | **0.98** | **0.98** | **4.35** | **3.74%** |
| DECREASERS | ***Leuconostoc lactis ASV15*** | **Age decreaser** | **2205** | **256** | **90.92** | **13.21** | **1861** | **2199** | **2502** | **1** | **1** | **13.43** | **1.89%** |
| DECREASERS | ***Mycoplasma; s_ ASV16*** | | **1096.5** | **34** | **20.8** | **6.5** | **325** | **1099** | **2210** | **0.992** | **0.982** | **7.46** | **0.87%** |
| DECREASERS | ***Lactococcus lactis ASV45*** | | **777.5** | **232** | **93.6** | **7.69** | **751** | **2138** | **2505** | **1** | **1** | **8.44** | **0.59%** |
| DECREASERS | ***Leuconostoc; s_ ASV35*** | **Age decreaser** | **1676** | **84** | **65.73** | **25.24** | **1490** | **1747** | **1826** | **1** | **1** | **25.01** | **0.57%** |
| DECREASERS | ***Lactococcus piscium ASV49*** | | **2494** | **198** | **74.62** | **13.92** | **2152** | **2230** | **2521** | **1** | **1** | **13.93** | **0.42%** |
| DECREASERS | *Citrobacter freundii ASV51* | | 2159 | 184 | 77.02 | 17.38 | 1884 | 2187 | 2494 | 1 | 1 | 17.78 | 0.35% |
| DECREASERS | *Romboutsia; s_ ASV41* | | 669 | 168 | 66.16 | 5.24 | 411 | 673 | 3153 | 0.998 | 0.998 | 5.47 | 0.30% |
| DECREASERS | *Acinetobacter; s_ ASV67* | Age decreaser | 2174.5 | 185 | 75.73 | 17.14 | 2151 | 2229 | 2502 | 1 | 1 | 17.16 | 0.29% |
| DECREASERS | *Ureaplasma; s_ ASV46* | | 2145 | 122 | 35.26 | 4.56 | 755 | 1854 | 2507 | 0.98 | 0.998 | 5.16 | 0.26% |
| DECREASERS | *Lactococcus; s_ ASV76* | Age decreaser | 2497.5 | 144 | 60.44 | 15.66 | 2149 | 2230 | 2505 | 1 | 1 | 16.28 | 0.20% |
| DECREASERS | *Weissella; s_ ASV92* | | 2238.5 | 183 | 74.51 | 10.59 | 2187 | 2250 | 2506 | 1 | 1 | 11.17 | 0.20% |
| DECREASERS | Enterobacteriaceae; g_ ASV81 | | 2522.5 | 151 | 56.63 | 9.14 | 2179 | 2521 | 2551 | 1 | 1 | 9.55 | 0.18% |
| DECREASERS | *Escherichia-Shigella; s_ ASV19* | Age increaser | 324.5 | 49 | 46.53 | 8.15 | 306 | 410 | 2169 | 0.978 | 0.97 | 8.46 | 0.16% |
| DECREASERS | *Terrisporobacter; s_ ASV65* | | 324.5 | 114 | 45.57 | 3.03 | 325 | 716 | 3265 | 0.986 | 0.976 | 4.31 | 0.15% |
| DECREASERS | *Streptococcus sp. ASV102* | | 2494 | 153 | 66.39 | 11.81 | 2186 | 2238 | 2505 | 1 | 1 | 11.72 | 0.14% |
| DECREASERS | *Acinetobacter; s_ ASV9* | Age decreaser | 2523.5 | 161 | 61.78 | 5.75 | 2204 | 2520 | 2551 | 1 | 1 | 6.61 | 0.14% |
| DECREASERS | *Acinetobacter; s_ ASV118* | Age decreaser | 1828.5 | 111 | 66.08 | 20.34 | 1579 | 1828 | 2150 | 1 | 1 | 20.92 | 0.11% |
| DECREASERS | *Cetobacterium; s_ ASV94* | Age decreaser | 1292.5 | 15 | 14.99 | 9.81 | 1100 | 1372 | 1593 | 1 | 1 | 10.30 | 0.09% |
| DECREASERS | *Acinetobacter brisouii ASV171* | | 2238 | 126 | 50.8 | 11.82 | 2187 | 2250 | 2506 | 1 | 1 | 12.59 | 0.07% |
| DECREASERS | *Edwardsiella; s_ ASV180* | | 1456 | 31 | 17.39 | 8.57 | 1090 | 1456 | 1809 | 1 | 1 | 9.41 | 0.06% |
| INCREASERS | ***Paeniclostridium; s_ ASV1*** | | **1870** | **342** | **60.46** | **7.76** | **1293** | **1848** | **2229** | **0.996** | **1** | **7.70** | **32.13%** |
| INCREASERS | ***Actinobacillus delphinicola ASV3*** | | **751** | **338** | **73.79** | **5.04** | **414** | **714** | **1208** | **0.978** | **0.996** | **6.05** | **11.36%** |
| INCREASERS | ***Fusobacterium; s_ ASV10*** | | **1821.5** | **128** | **40.31** | **5.02** | **1491** | **1822** | **2199** | **1** | **0.996** | **5.47** | **1.80%** |
| INCREASERS | ***Ureaplasma; s_ ASV17*** | | **2515** | **239** | **64.83** | **5.73** | **2197** | **2508** | **2533** | **1** | **1** | **5.89** | **1.41%** |
| INCREASERS | ***Mycoplasma; s_ ASV24*** | | **2515** | **147** | **43.87** | **5.84** | **2205** | **2513** | **2533** | **0.998** | **0.998** | **5.88** | **0.60%** |
| INCREASERS | *Romboutsia; s_ ASV4* | | 5076 | 99 | 71.45 | 8.24 | 2648 | 3412 | 5073 | 0.996 | 0.998 | 9.06 | 0.39% |
| INCREASERS | *Comamonas testosteroni ASV73* | | 2513.5 | 46 | 26.85 | 10.28 | 2505 | 2516 | 2540 | 1 | 1 | 10.74 | 0.28% |
| INCREASERS | *Mycoplasma; s_ ASV103* | | 2376.5 | 165 | 50.69 | 6.82 | 2204 | 2515 | 2551 | 0.996 | 1 | 7.10 | 0.14% |
| INCREASERS | *Mycoplasma; s_ ASV132* | | 2515 | 45 | 17.35 | 5.63 | 2505 | 2515 | 3325 | 0.994 | 0.982 | 5.75 | 0.12% |

### AGE OF SOUTHERN RESIDENT KILLER WHALE

| Group | Taxonomy | Study day indicator | zenv.cp | freq | IndVal | zscore | 5% | 50% | 95% | purity | reliability | z.median | Avg. RA |
|---|---|---|---|---|---|---|---|---|---|---|---|---|---|
| DECREASERS | *Leuconostoc lactis ASV15* | Study day decreaser | 20.58 | 256 | 65.47 | 6.62 | 18 | 19 | 20 | 1 | 1 | 6.59 | 1.89% |
| DECREASERS | *Cetobacterium; s_ ASV20* | | 21.205 | 160 | 42.02 | 6.28 | 19 | 22 | 31 | 0.996 | 1 | 6.97 | 0.77% |
| DECREASERS | *Leuconostoc; s_ ASV35* | Study day decreaser | 19.17 | 84 | 30.5 | 7.81 | 13 | 19 | 20 | 1 | 1 | 8.49 | 0.57% |
| DECREASERS | *Paeniclostridium; s_ ASV40* | | 6.44 | 92 | 46.39 | 7.24 | 5 | 7 | 36 | 1 | 0.996 | 6.83 | 0.49% |
| DECREASERS | *Paeniclostridium; s_ ASV44* | | 6.44 | 99 | 50.94 | 7.69 | 6 | 7 | 21 | 1 | 0.998 | 7.52 | 0.45% |
| DECREASERS | *Acinetobacter; s_ ASV67* | Study day decreaser | 19.17 | 185 | 45.3 | 5.29 | 12 | 20 | 21 | 0.99 | 0.994 | 5.73 | 0.29% |
| DECREASERS | *Lactococcus; s_ ASV76* | Study day decreaser | 18.26 | 144 | 34.18 | 3.86 | 6 | 18 | 21 | 0.976 | 0.968 | 4.84 | 0.20% |
| DECREASERS | *Acinetobacter; s_ ASV9* | Study day decreaser | 20.915 | 160 | 42.35 | 3.24 | 4 | 18 | 35 | 0.99 | 0.986 | 4.03 | 0.14% |
| DECREASERS | *Acinetobacter; s_ ASV118* | Study day decreaser | 19.99 | 111 | 32.38 | 6.07 | 7 | 19 | 20 | 0.986 | 1 | 6.62 | 0.11% |
| DECREASERS | *Cetobacterium; s_ ASV94* | Study day decreaser | 17.11 | 15 | 9.91 | 5.72 | 11 | 17 | 18 | 0.996 | 1 | 6.90 | 0.09% |
| DECREASERS | *Actinobacillus; s_ ASV162* | | 10.1 | 12 | 10.91 | 5.96 | 5 | 10 | 23 | 0.998 | 0.974 | 7.08 | 0.08% |
| DECREASERS | *Paeniclostridium; s_ ASV181* | | 18.125 | 12 | 7.69 | 6.58 | 3 | 18 | 20 | 0.986 | 1 | 6.81 | 0.05% |
| INCREASERS | *Plesiomonas; s_ ASV13* | | 39.1 | 144 | 50.1 | 6.79 | 37 | 45 | 100 | 1 | 1 | 7.21 | 1.84% |
| INCREASERS | *Clostridium SS1; s_ ASV30* | | 8.215 | 143 | 41.89 | 2.56 | 8 | 13 | 47 | 0.98 | 0.996 | 3.69 | 0.40% |
| INCREASERS | *Paeniclostridium; s_ ASV48* | | 97.645 | 58 | 54.96 | 9.48 | 54 | 79 | 100 | 0.996 | 1 | 10.67 | 0.34% |
| INCREASERS | *Ligilactobacillus; s_ ASV69* | | 97.645 | 168 | 87.73 | 6.81 | 52 | 74 | 98 | 1 | 1 | 7.14 | 0.16% |
| INCREASERS | *Escherichia-Shigella; s_ ASV19* | Study day decreaser | 22.64 | 48 | 18.97 | 3.7 | 20 | 22 | 33 | 0.98 | 0.984 | 4.25 | 0.16% |
| INCREASERS | *Fusobacterium; s_ ASV134* | | 77.61 | 20 | 29.61 | 10.1 | 21 | 77 | 79 | 0.984 | 0.988 | 10.00 | 0.12% |
| INCREASERS | *Paeniclostridium; s_ ASV183* | | 24.21 | 7 | 5 | 5.07 | 22 | 31 | 39 | 1 | 0.984 | 5.61 | 0.11% |

**Supplementary Table 8. The temporal core microbiota.** The Southern Resident killer whale (SRKW) dataset was divided into four time periods, each spanning 3-4 consecutive years, and the prevalence and relative abundance (RA) of each ASV was calculated. ASVs that were prevalent (≥50%) and abundant (mean RA ≥1%) in a given time period were considered "core" microbiota and are highlighted in pink. Those that were consistently classified as core throughout the study are outlined and labeled as "SRKW Temporal Core". For ASVs that were not consistently detected throughout the 2005—2019 study period, the first and last year during which they were detected (RA >0) are shown. (2005 and 2019 are indicated in grey font to highlight whether taxa were emerging, blooming or vanishing.) For reference, the right column shows whether each ASV was previously classified as core for a wild KW population(s) in **Supplementary Table 2** (using just the SRKW data from 2017–2019).

| | | Emerging & vanishing ASVs (study duration: 2005-2019) | | | | | 2005-2008 (n=86) | | | 2009-2011 (n=119) | | | 2012-2015 (n=115) | | | 2017-2019 (n=22) | | | | |
|---|---|---|---|---|---|---|---|---|---|---|---|---|---|---|---|---|---|---|---|---|
| ASV | ASV taxonomy | Trend | First detected | Last detected | Mean RA (full study) | Mean Prev (full study) | RA | Prevalence | SRKW core? | RA | Prevalence | SRKW core? | RA | Prevalence | SRKW core? | RA | Prevalence | SRKW core? | | Wild Core 2017-2019 (**Table S2**)? |
| ASV1 | Paeniclostridium; s_ASV1 | CORE | | | 30.95% | 100.00% | 23.16% | 100.00% | Core | 30.96% | 100.00% | Core | 39.69% | 100.00% | Core | 29.98% | 100.00% | Core | **SRKW temporal core** | Universal wild KW core |
| ASV3 | Actinobacillus delphinicola ASV3 | CORE | | | 11.44% | 98.15% | 8.73% | 98.84% | Core | 12.77% | 99.16% | Core | 11.34% | 99.13% | Core | 12.90% | 95.45% | Core | | Resident KW core |
| ASV6 | Cetobacterium; s_ASV6 | CORE | | | 5.46% | 94.47% | 5.22% | 91.86% | Core | 5.05% | 95.80% | Core | 2.79% | 94.78% | Core | 8.78% | 95.45% | Core | | Resident KW core |
| ASV7 | Clostridium_SS_1; s_ASV7 | CORE | | | 4.31% | 91.83% | 4.71% | 97.67% | Core | 4.93% | 94.12% | Core | 3.59% | 98.26% | Core | 4.02% | 77.27% | Core | | Resident KW core |
| ASV5 | Edwardsiella tarda ASV5 | CORE | | | 8.67% | 97.52% | 12.17% | 98.84% | Core | 7.23% | 96.64% | Core | 5.73% | 99.13% | Core | 9.56% | 95.45% | Core | | SRKW & ARKW core |
| ASV8 | Fusobacterium; s_ASV8 | CORE | | | 4.16% | 70.49% | 3.97% | 73.26% | Core | 3.62% | 73.95% | Core | 3.62% | 75.65% | Core | 5.41% | 59.09% | Core | | SRKW-specific core |
| ASV2 | Photobacterium; s_ASV2 | CORE | | | 3.40% | 94.85% | 4.91% | 96.51% | Core | 4.05% | 92.44% | Core | 2.82% | 90.43% | Core | 1.83% | 100.00% | Core | | SRKW & TKW core |
| ASV15 | Leuconostoc lactis ASV15 | Vanish | 2005 | 2014 | 2.23% | 80.70% | 4.89% | 90.70% | Core | 1.73% | 97.48% | Core | 0.08% | 53.91% | | | | | | |
| ASV35 | Leuconostoc; s_ASV35 | Bloom | 2006 | 2013 | 0.69% | 29.34% | 1.67% | 62.79% | Core | 0.40% | 24.37% | | 0.00% | 0.87% | | | | | | |
| ASV13 | Plesiomonas; s_ASV13 | Bloom | 2006 | 2017 | 1.48% | 36.11% | 2.23% | 52.33% | Core | 2.72% | 37.82% | | 0.96% | 45.22% | | 0.03% | 9.09% | | | |
| ASV45 | Lactococcus lactis ASV45 | Vanish | 2005 | 2018 | 0.52% | 56.63% | 1.67% | 96.51% | Core | 0.37% | 93.28% | | 0.03% | 32.17% | | 0.00% | 4.55% | | | |
| ASV49 | Lactococcus piscium ASV49 | Vanish | 2005 | 2014 | 0.46% | 62.43% | 0.70% | 72.09% | | 0.64% | 88.24% | | 0.04% | 26.96% | | | | | | |
| ASV20 | Cetobacterium; s_ASV20 | | | | 0.86% | 51.33% | 0.80% | 51.16% | | 1.00% | 52.94% | Core | 0.40% | 33.04% | | 1.24% | 68.18% | Core | | Resident KW core |
| ASV17 | Ureaplasma; s_ASV17 | | | | 1.40% | 67.65% | 0.63% | 68.60% | | 0.43% | 66.39% | | 2.93% | 76.52% | Core | 1.61% | 59.09% | Core | | Resident KW core |
| ASV33 | Ureaplasma; s_ASV33 | | | | 0.54% | 70.12% | 0.42% | 79.07% | | 0.29% | 71.43% | | 1.16% | 80.00% | Core | 0.29% | 50.00% | | | ARKW-specific core |
| ASV25 | Actinobacillus; s_ASV25 | | | | 0.62% | 54.32% | 0.19% | 48.84% | | 0.99% | 57.98% | | 0.47% | 69.57% | | 0.80% | 40.91% | | | ARKW-specific core |
| ASV66 | Actinobacillus; s_ASV66 | | | | 0.24% | 51.78% | 0.17% | 55.81% | | 0.21% | 57.14% | | 0.22% | 48.70% | | 0.38% | 45.45% | | | NRKW-specific core |
| ASV24 | Mycoplasma; s_ASV24 | | | | 0.63% | 43.62% | 0.22% | 44.19% | | 0.22% | 35.29% | | 1.21% | 49.57% | | 0.89% | 45.45% | | | NRKW & ARKW core |
| ASV16 | Mycoplasma; s_ASV16 | Vanish | 2005 | 2014 | 1.02% | 11.61% | 2.03% | 22.09% | | 0.79% | 8.40% | | 0.24% | 4.35% | | | | | | |
| ASV46 | Ureaplasma; s_ASV46 | Emerge | 2006 | 2019 | 0.24% | 35.20% | 0.31% | 40.70% | | 0.38% | 36.97% | | 0.09% | 31.30% | | 0.20% | 31.82% | | | |
| ASV10 | Fusobacterium; s_ASV10 | | | | 2.60% | 37.53% | 0.12% | 27.91% | | 2.37% | 36.97% | | 1.54% | 44.35% | | 6.38% | 40.91% | | | |
| ASV11 | Fusobacterium necrogenes ASV11 | Emerge | 2006 | 2019 | 1.64% | 39.17% | 1.51% | 45.35% | | 2.33% | 38.66% | | 1.20% | 40.87% | | 1.52% | 31.82% | | | |
| ASV28 | Clostridium_SS_1; s_ASV28 | | | | 0.67% | 71.58% | 0.80% | 75.58% | | 0.71% | 68.91% | | 0.90% | 60.00% | | 0.26% | 81.82% | | | |
| ASV22 | Tyzzerella; s_ASV22 | | | | 0.34% | 26.12% | 0.70% | 32.56% | | 0.13% | 24.37% | | 0.53% | 33.91% | | 0.02% | 13.64% | | | |
| ASV51 | Citrobacter freundii ASV51 | Vanish | 2005 | 2014 | 0.39% | 58.86% | 0.64% | 77.91% | | 0.48% | 88.24% | | 0.05% | 10.43% | | | | | | |
| ASV67 | Acinetobacter; s_ASV67 | Vanish | 2005 | 2014 | 0.32% | 58.83% | 0.59% | 74.42% | | 0.35% | 90.76% | | 0.03% | 11.30% | | | | | | |
| ASV34 | Clostridium_SS_1; s_ASV34 | Emerge | 2006 | 2019 | 0.71% | 37.42% | 0.56% | 38.37% | | 0.40% | 31.93% | | 0.37% | 33.91% | | 1.52% | 45.45% | | | |
| ASV41 | Romboutsia; s_ASV41 | | | | 0.26% | 43.34% | 0.56% | 56.98% | | 0.21% | 51.26% | | 0.23% | 46.96% | | 0.02% | 18.18% | | | |
| ASV40 | Paeniclostridium; s_ASV40 | | | | 0.48% | 27.49% | 0.45% | 34.88% | | 0.39% | 26.05% | | 0.63% | 21.74% | | 0.44% | 27.27% | | | |
| ASV165 | Limosilactobacillus; s_ASV165 | Vanish | 2005 | 2014 | 0.15% | 35.97% | 0.44% | 58.14% | | 0.01% | 43.70% | | 0.00% | 6.09% | | | | | | |
| ASV76 | Lactococcus; s_ASV76 | Vanish | 2005 | 2014 | 0.25% | 45.70% | 0.44% | 56.98% | | 0.29% | 71.43% | | 0.03% | 8.70% | | | | | | |
| ASV44 | Paeniclostridium; s_ASV44 | | | | 0.46% | 30.06% | 0.41% | 37.21% | | 0.36% | 27.73% | | 0.54% | 23.48% | | 0.52% | 31.82% | | | |
| ASV19 | Escherichia-Shigella; s_ASV19 | Emerge | 2006 | 2019 | 0.13% | 14.56% | 0.39% | 19.77% | | 0.04% | 10.92% | | 0.11% | 13.91% | | 0.00% | 13.64% | | | |
| ASV14 | Fusobacterium; s_ASV14 | | | | 1.06% | 35.52% | 0.37% | 36.05% | | 0.79% | 36.13% | | 1.73% | 42.61% | | 1.36% | 27.27% | | | |
| ASV50 | Cetobacterium; s_ASV50 | Bloom | 2007 | 2011 | 0.24% | 2.42% | 0.37% | 2.33% | | 0.11% | 2.52% | | | | | | | | | |
| ASV39 | Fusobacterium; s_ASV39 | | 2006 | 2019 | 0.35% | 15.28% | 0.36% | 11.63% | | 0.54% | 19.33% | | 0.28% | 16.52% | | 0.23% | 13.64% | | | |
| ASV94 | Cetobacterium; s_ASV94 | Bloom | 2006 | 2009 | 0.17% | 8.40% | 0.33% | 15.12% | | 0.01% | 1.68% | | | | | | | | | |
| ASV120 | Bacteroides; s_ASV120 | Bloom | 2007 | 2013 | 0.11% | 1.92% | 0.32% | 2.33% | | 0.00% | 0.84% | | 0.00% | 2.61% | | | | | | |
| ASV81 | Enterobacteriaceae; g_ASV81 | | 2005 | 2019 | 0.15% | 36.89% | 0.30% | 59.30% | | 0.26% | 68.07% | | 0.03% | 15.65% | | 0.00% | 4.55% | | | |
| ASV118 | Acinetobacter; s_ASV118 | Bloom | 2006 | 2014 | 0.14% | 37.12% | 0.29% | 65.12% | | 0.11% | 45.38% | | 0.00% | 0.87% | | | | | | |
| ASV54 | Photobacterium damselae ASV54 | Emerge | 2006 | 2019 | 0.18% | 12.64% | 0.24% | 11.63% | | 0.07% | 9.24% | | 0.14% | 6.96% | | 0.26% | 22.73% | | | |
| ASV92 | Weissella; s_ASV92 | Vanish | 2005 | 2014 | 0.20% | 58.17% | 0.23% | 73.26% | | 0.34% | 89.08% | | 0.03% | 12.17% | | | | | | |
| ASV4 | Romboutsia; s_ASV4 | | 2006 | 2019 | 0.44% | 42.31% | 0.22% | 26.74% | | 0.24% | 22.69% | | 0.61% | 24.35% | | 0.69% | 95.45% | | | |
| ASV48 | Paeniclostridium; s_ASV48 | Emerge | 2006 | 2019 | 0.30% | 15.00% | 0.18% | 10.47% | | 0.17% | 12.61% | | 0.65% | 27.83% | | 0.20% | 9.09% | | | |
| ASV30 | Clostridium_SS_1; s_ASV30 | | | | 0.32% | 37.55% | 0.18% | 50.00% | | 0.43% | 42.02% | | 0.57% | 40.00% | | 0.10% | 18.18% | | | |
| ASV80 | Mycoplasma; s_ASV80 | | | | 0.18% | 39.34% | 0.11% | 40.70% | | 0.14% | 35.29% | | 0.17% | 49.57% | | 0.32% | 31.82% | | | |
| ASV57 | Tyzzerella; s_ASV57 | Bloom | 2007 | 2014 | 0.22% | 9.20% | 0.10% | 13.95% | | 0.54% | 7.56% | | 0.02% | 6.09% | | | | | | |
| ASV27 | Bacteroides; s_ASV27 | Emerge | 2006 | 2019 | 0.19% | 25.66% | 0.09% | 13.95% | | 0.07% | 21.01% | | 0.20% | 31.30% | | 0.39% | 36.36% | | | |
| ASV162 | Actinobacillus; s_ASV162 | Emerge | 2006 | 2019 | 0.18% | 7.74% | 0.09% | 6.98% | | | | | 0.11% | 2.61% | | 0.34% | 13.64% | | | |
| ASV61 | Tyzzerella; s_ASV61 | Bloom | 2007 | 2014 | 0.14% | 6.11% | 0.08% | 8.14% | | 0.32% | 6.72% | | 0.01% | 3.48% | | | | | | |
| ASV86 | Actinobacillus; s_ASV86 | | | | 0.13% | 23.31% | 0.08% | 37.21% | | 0.34% | 26.05% | | 0.08% | 20.87% | | 0.02% | 9.09% | | | |
| ASV103 | Mycoplasma; s_ASV103 | | | | 0.14% | 47.51% | 0.06% | 44.19% | | 0.04% | 38.66% | | 0.29% | 61.74% | | 0.18% | 45.45% | | | |
| ASV71 | Bacteroides; s_ASV71 | Emerge | 2006 | 2019 | 0.44% | 9.95% | 0.04% | 9.30% | | 0.07% | 9.24% | | 0.59% | 12.17% | | 1.06% | 9.09% | | | |
| ASV115 | Mycoplasma; s_ASV115 | Emerge | 2006 | 2019 | 0.11% | 9.14% | 0.02% | 5.81% | | 0.03% | 7.56% | | 0.35% | 9.57% | | 0.03% | 13.64% | | | |
| ASV73 | Comamonas testosteroni ASV73 | Bloom | 2007 | 2015 | 0.28% | 13.76% | 0.00% | 5.81% | | 0.04% | 5.04% | | 0.79% | 30.43% | | | | | | |
| ASV89 | Aliivibrio; s_ASV89 | Bloom | 2006 | 2012 | 0.14% | 6.63% | 0.00% | 10.47% | | 0.00% | 4.20% | | 0.43% | 5.22% | | | | | | |
| ASV170 | Candidatus_Arthromitus; s_ASV170 | Emerge | 2008 | 2019 | 0.07% | 4.01% | 0.00% | 1.16% | | 0.01% | 2.52% | | 0.27% | 7.83% | | 0.00% | 4.55% | | | |
| ASV183 | Paeniclostridium; s_ASV183 | Emerge | 2011 | 2019 | 0.25% | 4.44% | | | | 0.19% | 3.36% | | 0.02% | 0.87% | | 0.55% | 9.09% | | | |
| ASV436 | Bacteroides; s_ASV436 | Emerge | 2019 | 2019 | 0.30% | 4.55% | | | | | | | | | | 0.30% | 4.55% | | | |
| ASV249 | Fusobacterium; s_ASV249 | Emerge | 2009 | 2019 | 0.16% | 2.69% | | | | 0.00% | 0.84% | | | | | 0.32% | 4.55% | | | |
| ASV21 | Cetobacterium; s_ASV21 | Bloom | 2011 | 2012 | 0.38% | 4.65% | | | | 0.62% | 7.56% | | 0.13% | 1.74% | | | | | | |

**Supplementary Table 9. Differential abundance analysis of fecal microbiotas from pregnant Southern Resident killer whales (SRKW) with poor outcomes and from nonpregnant females.** Amplicon sequence variant (ASV) differential relative abundance (DA) was analyzed in the 11 SRKW females that were sampled during a reproductive event (**Fig. S18**). Eight of these females were also sampled when presumably nonpregnant and nonlactating (“nonpregnant”, n=31) — these fecal microbiotas were compared to those from 6 pregnant SRKWs that had poor pregnancy outcomes (n=15). Only taxa with ≥1000 reads in ≥1 sample were considered. ASV DA was determined using coda4microbiome and ALDEx2. Asterisks indicate robust indicator taxa found to have DA via both methods. Samples with low confidence assignments to a reproductive event (i.e., “Possibly” pregnant) were excluded.

| **Pregnant** (poor outcome, n=15) vs **nonpregnant, nonlactating** (n=31) females | | **ALDEx2** (denom="mean") | | | **ALDEx2** (denom="zero") | | | **CODA** | | **Relative abundance** | | **Prevalence** | |
|---|---|---|---|---|---|---|---|---|---|---|---|---|---|
| **Differentially abundant taxon** | **Enriched group (>1 DA method)** | **Enriched** | **abs(effect)** | **p-adj** | **Enriched** | **abs(effect)** | **p-adj** | **Enriched** | **coeff** | **Poor outcome** | Nonpregnant | **Poor outcome** | Nonpregnant |
| *Fusobacterium necrogenes* ASV11 * | **Pregnant SRKWs with poor outcome** | **Poor outcome** | 1.14 | 9.86E-03 | **Poor outcome** | 1.06 | 5.86E-03 | **Poor outcome** | 1.00 | 7.36% | 0.44% | 80.00% | 22.58% |
| *Romboutsia* ; s_ASV41 | | | | | | | | Nonpregnant | 0.47 | 0.02% | 0.40% | 26.67% | 64.52% |
| *Clostridium SS1* ; s_ASV7 | | | | | | | | Nonpregnant | 0.38 | 1.16% | 5.31% | 93.33% | 100.00% |
| *Actinobacillus* ; s_ASV87 | | | | | | | | Nonpregnant | 0.15 | 0.00% | 0.04% | 6.67% | 6.45% |

**Supplementary Table 10. Average nucleotide identity (ANI) of the two novel *Fusobacterium* MAGs recovered from a SRKW L26 fecal sample versus NCBI reference genomes.**

| ***Fusobacterium*** | | | | | | | | | | | | | | | |
|---|---|---|---|---|---|---|---|---|---|---|---|---|---|---|---|
| SRKW (L26) MAGs | | NCBI Reference Genomes | | | | | | | | | | | | | |
| MAG | L26_86 | *F. hominis* | *F. mortiferum* | *F. necrogenes* | *F. varium* | *F. ulcerans* | *F. perfoetens* | *F. canifelinum* | *F. periodonticum* | *F. polymorphum* | *F. nucleatum* | *F. animalis* | *F. russi* | *F. gonidiaform* | *F. necrophorum* |
| L26_30: *F.mortiferum* -like | 72.44% | **78.85%** | 77.16% | 76.57% | 74.87% | 74.43% | 72.65% | 72.56% | 72.40% | 72.31% | 72.27% | 72.21% | 71.80% | 71.61% | 70.75% |
| L26_86: *Fusobacterium* | | 72.50% | 73.34% | 72.59% | 72.56% | 72.77% | **75.34%** | 70.98% | 71.06% | 71.26% | 71.07% | 71.14% | 70.08% | 69.75% | 69.16% |

**Supplementary Table 11. An extended set of putative virulence factor (VF) genes identified in the two novel *Fusobacterium* MAGs recovered from a SRKW L26 fecal sample.** The two SRKW *Fusobacterium* metagenomically-assembled genomes (MAGs) were annotated using the Prokka and Bakta databases in the Proksee online interface. Gene annotations were compared to a custom VF database from VFDB[1] and relevant literature. Putative VF genes were analyzed using BLASTN ("B") to identify their most closely related homologues. Sequences were also analyzed using FusoPortal[2] ("FP") and bold text indicates VFs previously identified in *Fusobacterium* reference genomes[3]. Query sequences for each gene are provided in **Supplementary Data 10.**

| | | | **L26 F. mortiferum-like MAG** (L26_F_mort) | | | | **L26 unclassified *Fusobacterium* MAG** (L26_Fuso) | | | |
|---|---|---|---|---|---|---|---|---|---|---|
| **Virulence factor (VF) class** | **VF gene** | **Documented role in pathogenesis** (reference) | **Closest hit** (database) | **identity** | **score** | **E-value** | **Closest hit** (database) | **identity** | **score** | **E-value** |
| Adherence & biofilm | *ompA* | Host cell-binding porin (1,2,7) | *F. hominis* (B) | 77.40% | 398 | 6.90E-106 | | | | |
| Stress survival | *dnaK* | Stress tolerance (heat, oxidative & dessication); porin (1-4) | *F. mortiferum* (FP) | 90.30% | 2490 | 0.00E+00 | *F. sphaericum* (B) | 92.23% | 2655 | 0.00E+00 |
| Antiphagocytosis | *galE* | Capsule biosynthesis (1,2) | *F. hominis* (B) | 85.10% | 1119 | 0.00E+00 | *F. sphaericum* (B) | 87.12% | 1140 | 0.00E+00 |
| VF regulation | *cvfB* | Conserved virulence factor; hemolysin production (16) | *F. hominis* (B) | 84.80% | 948 | 0.00E+00 | *F. sphaericum* (B) | 87.10% | 1036 | 0.00E+00 |
| Biofilm | *dnaJ* | Biofilm, dessication tolerance (heat shock protein) (1-3) | *F. mortiferum* (FP) | 82.60% | 815 | 0.00E+00 | *F. sphaericum* (B) | 86.89% | 1471 | 0.00E+00 |
| Adherence & biofilm | *yiaD (OmpA)* | Host cell-binding porin (1,2,7) | *F. mortiferum* (FP) | 82.60% | 652 | 0.00E+00 | *F. sphaericum* (B) | 84.50% | 713 | 0.00E+00 |
| Metabolism | *pfo* | Pyruvate metabolism (oxidoreductase), biofilm (2,5) | *F. periodonticum* (B) | 81.10% | 6330 | 0.00E+00 | *F. sphaericum* (B) | 93.70% | 5400 | 0.00E+00 |
| Immune evasion | *kpsF* | Capsule biosynthesis, antiphagocytosis (1,2) | *F. hominis* (B) | 80.90% | 912 | 0.00E+00 | *F. varium* (B) | 76.77% | 696 | 0.00E+00 |
| Pore-forming toxins | *hlyC/corC* | Hemolysin; host cell lysis (activates HlyA toxin) (1,2) | *F. mortiferum* (FP) | 80.30% | 1139 | 0.00E+00 | *F. sphaericum* (B) | 83.78% | 1394 | 0.00E+00 |
| Pore-forming toxins | *hemolysin* | Hemolysin; host cell lysis (1,2) | *F. hominis* (B) | 78.90% | 532 | 5.23E-146 | *F. sphaericum* (B) | 77.29% | 353 | 2.00E-92 |
| Antiphagocytosis | *murJ* | Capsule/peptidoglycan (PG) biosynthesis (1,2) | *F. hominis* (B) | 77.10% | 1104 | 0.00E+00 | *F. varium* (FP) | 72.92% | 703 | 0.00E+00 |
| Pore-forming toxins | *tlyA* | Hemolysin; host cell lysis (RNA methyltransferase) (1,2) | *F. hominis* (B) | 75.70% | 573 | 2.24E-158 | *F. sphaericum* (B) | 84.93% | 918 | 0.00E+00 |
| Pore-forming toxins | *hlyD* | Hemolysin; host cell lysis (membrane fusion protein) (1,2) | *F. varium* (FP) | 73.60% | 531 | 2.17E-150 | *F. varium* (B) | 73.42% | 663 | 0.00E+00 |
| Pore-forming toxins | *cirA* | Cytotoxic activity (Colicin I receptor) (1,8) | *F. varium* (FP) | 68.80% | 551 | 2.95E-156 | *F. ulcerans* (FP) | 78.03% | 389 | 6.47E-108 |
| Exotoxins | *epsF* | T2SS toxin secretion (protein F) (1) | *F. mortiferum* (FP) | 64.79% | 61 | 1.00E-08 | *F. sphaericum* (B) | 84.54% | 1181 | 0.00E+00 |
| Motility | *fliY* | Flagellum-mediated motility (1) | *F. mortiferum* (FP) | 79.70% | 613 | 5.08E-175 | *F. sphaericum* (B) | 82.36% | 778 | 0.00E+00 |
| Stress survival | *coa_dr* | ROS detoxification (CoA-disulfide reductase) (2,6) | *F. mortiferum* (B) | 77.88% | 1315 | 0.00E+00 | *F. mortiferum* (FP) | 81.85% | 1325 | 0.00E+00 |
| T3SS | *ctpA* | Cytotoxicity (S41 serine protease) (2,9) | *F. hominis* (B) | 83.50% | 1270 | 0.00E+00 | | | | |
| Adherence & biofilm | *upaG (YadA)* | Fibronectin and laminin-binding adhesin (1,2,12) | *F. necrophorum* (FP) | 77.20% | 59 | 3.64E-08 | | | | |
| Adherence | *ompH* | Host cell-binding periplasmic chaperone (11) | *F. mortiferum* (FP) | 75.00% | 263 | 5.40E-70 | | | | |
| Immune evasion | *brkB* | Resists complement-mediated killing in serum (1,2) | *F. hominis* (B) | 73.00% | 681 | 0.00E+00 | | | | |
| T4SS | *virB4* | T4SS protein; replication within host (1,2) | *F. varium* (B) | 70.80% | 650 | 7.00E-158 | | | | |
| Invasion | *MORN repeat* | Outer membrane transport protein (2,10) | *F. varium* (FP) | 70.40% | 324 | 4.32E-88 | | | | |
| T4SS | *virB11* | T4SS ATPase; replication within host (1,2) | *F. varium* (B) | 69.20% | 333 | 4.00E-86 | | | | |
| Pore-forming toxins | *shlB* | Hemolysin transporter (1,2) | *F. ulcerans* (FP) | 69.00% | 54 | 1.67E-06 | | | | |
| Pore-forming toxins | Septicolysin | Cytolysin (cholesterol-dependent) (15) | | | | | *F. mortiferum* (FP) | 82.57% | 467 | 2.76E-131 |
| Stress survival | *creD* | Cell envelope integrity (1,13) | | | | | *F. mortiferum* (FP) | 82.68% | 1318 | 0.00E+00 |
| VF regulation | *TldD* | Regulates hemolysis (TldD/PmbA family) (14) | | | | | *F. sphaericum* (B) | 80.19% | 1217 | 0.00E+00 |

Database: (B)= BLAST, (FP)= FusoPortal
