## Supplementary Text for "Fifteen-year microbiome survey of endangered killer whales (*Orcinus orca*) reveals declining diversity and population differences"

### Supplementary Methods

#### Amplicon sequence variant inference and taxonomic assignment

Demultiplexed reads were processed using the open-source package DADA2 (v1.34.0) and the parameters recommended in the "Big Data: Paired-end" workflow[1] in R (v4.4.3, **Supplementary Code 1**). Read quality was assessed for each lane individually: truncLen$_{Pool1}$ = c(245, 200) and truncLen$_{Pool2}$ = c(245, 220). Sequences from both lanes were merged prior to chimera removal. Taxonomy was assigned using a naïve Bayesian classifier (‘assignTaxonomy’, DADA2) and the SILVA v138 reference database (formatted for DADA2)[2]. The ‘addSpecies’ DADA2 function was used to make species-level assignments for amplicon sequence variants (ASVs) with exact matches to sequences in the reference database. A rooted phylogenetic tree was then created in QIIME2[3] by inserting the ASVs into the SILVA v138 reference alignment (‘qiime fragment-insertion sepp’) using SATé-enabled phylogenetic placement[4].

#### Quality filtering

Two additional filtering approaches were used to remove extremely rare taxa that may reflect sequencing artifacts[1] and to create ps objects for downstream analyses. For alpha diversity analyses, a permutation test was used to identify and remove potentially spurious ASVs while preserving richness (PERFect). This yielded the “ps_alpha” object with 9,813 ASVs (72,639,633 reads) in 671 samples. For all other analyses, only ASVs present in ≥2 samples with ≥100 reads/sample were retained. This yielded “ps_filt” with 780 ASVs (71,869,905 reads) in 669 samples. The rationale behind this more stringent filtering approach was to emphasize biologically relevant taxa and to reduce computation time.

#### Overall 16S rRNA gene amplicon sequence dataset

The wild KW dataset (considering ps_alpha) consisted of 384 fecal samples (resident KWs, n = 381; TKWs, n = 3) with 6,460 ASVs (**Supplementary Code 2**), and 48,700,808 reads (mean = 126,825 reads per sample; range = 4,177–649,264 reads per sample), and 10 seawater samples with 863 ASVs and 575,578 reads (mean = 57,555; range = 37,259–85,688). The AqKW dataset consisted of 275 fecal samples with 3,309 ASVs and 23,249,677 reads (mean = 81,320; range = 6,253–205,417), and two pool water samples with 1,028 ASVs and 113,601 reads (mean = 56,800; range = 34,538–79,063).

#### Statistical analyses and data normalization

Microbial community profiles were primarily analyzed using phyloseq[5] (**Supplementary Code 2**). To account for differences in sample size between KW subgroups (e.g., populations), normalization was performed by random subsampling ('sample_n', dplyr v1.1.4). To control for time, only timeframes with samples available for all subgroups were considered. To account for differences in sequencing depth, rarefaction was performed ('rarefy_even_depth', phyloseq) prior to analysis of richness, dominance, ASV turnover, and Unifrac distances. For Aitchison-distance based analyses, raw read counts were center log ratio-transformed (clr, 'ord_calc', microViz v0.12.4). For analysis of taxonomic relative abundance (RA), data were proportion-transformed (‘transform’, microbiome v1.28.0). For all other analysis, raw read counts were used.

#### Alpha diversity and relative abundance

Alpha diversity metrics were examined using the ‘plot-richness’ (phyloseq) and ‘alpha’ (microbiome) functions. All alpha diversity analyses were performed using both Shannon index (using raw read counts) and observed species counts (using rarefied data) to capture both richness and evenness. To compare alpha diversity between KW groups (ecotypes, populations, or husbandry groups), data were normalized to ≤3 samples/individual KW to control for repeated measures (while retaining all samples from the smallest subgroup) and then normalized to match the size of the smallest comparison group. After examining comparison group normality ('shapiro.test', stats v4.4.3) and variance ('bartlett.test', stats), Dunn tests (for pairwise comparison of heterogeneous groups) or t-tests (for homogeneous groups or comparison to one reference group) were performed with Benjamini-Hochberg (BH) p-value adjustment ('geom_pwc', ggpubr v0.6.0). For RA analyses, ASVs were glommed to the taxonomic level of interest (‘tax_glom’, phyloseq) and the ‘preDA’ function (DAtest 2.8.0) was used to merge rare taxa before compositional transformation of the data. Alluvial RA plots were created using ‘ggalluvial’ (v0.12.5). Dominance was examined by identifying the most abundant ASV in each sample and calculating the relative frequency at which each ASV was dominant across the dataset (‘dominant_taxa’, microbiomeutilities v1.00.17). The RA of lactic acid producing bacteria was examined by filtering the data to the following genera: *Lactobacillus, Lactococcus, Leuconostoc, Pediococcus, Streptococcus, Aerococcus, Alloiococcus, Carnobacterium,*

*Dolosigranulum, Enterococcus, Oenococcus, Tetragenococcus, Vagococcus, Weissella, Sporolactobacillus, Ligilactobacillus and Bifidobacterium*[6,7].

#### Beta diversity and analysis of variance

Bray-Curtis, weighted Unifrac, unweighted Unifrac (UWU) and Aitchison dissimilarity distance metrics were used for unsupervised analyses of the data. Aitchison dissimilarity, designed for compositional data and more robust to subsetting[8,9], revealed the clearest partitioning of the data and was used moving forward. However, the main analyses were repeated in the supplemental material using UWU to evaluate whether Aitchison results were primarily driven by abundance. Aitchison distance matrices were created using 'ord_calc' (microViz v0.12.4) and visualized using an unconstrained principal component analysis ('ord_plot'). The relative contribution of study covariates to community variation was determined using permutational MANOVA (PERMANOVA, 'adonis2', vegan v2.6-10). KW ID was used in the permutation block (nperm=999) to account for repeated measures. Group dispersions (average Aitchison distance to the centroid) were determined using ‘betadisper’ (vegan) and homogeneity was assessed with ANOVA. Univariate distribution side panel boxplots were created using ggside (v0.3.1) and significant differences were identified along each axis using a Dunn test (‘geom_pwc’).

#### Differential abundance analyses

A consensus approach utilizing four differential abundance (DA) methods was used to identify differentially abundant amplicon sequence variants (ASVs) in the killer whale fecal microbiotas that were robust indicators of husbandry group (i.e., aquarium-house vs wild residents, **Supplementary Fig. 9**) or specific wild resident population, i.e., the Southern (SRKW), Northern (NRKW), and Alaska (ARKW) Residents (**Fig. 2b**, **Supplementary Fig. 7**). Each wild resident population was compared to both others (using samples collected 2017—2019) to identify differentially abundant ASVs that best distinguished the reference population (effect>0). For the endangered SRKWs, differential ASVs for both relatively fit NRKW and ARKW populations (effect<0, “Fit Pops”) were also considered relevant as possible indicators of population health.

The primary analysis utilized the MaAsLin2[10] R package (v1.20.0), which is designed to handle both count and compositional data. Multiple MaAsLin2 workflows were explored (**Supplementary Code 2**) to account for structural differences between datasets (skewed sample size, library size, and/or zero-inflation between comparison groups). Read counts and relative abundances (RA) were analyzed using the developer's recommendations[11]. Normalization techniques were chosen based on input type: Trimmed Mean of M-values (TMM) for counts and RA, Cumulative Sum Scaling (CSS) for counts, Center Log Ratio (CLR) for RA, and Total Sum Scaling (TSS) for RA. For counts, Tweedie compound Poisson linear models (NEGBIN) were run without data transformation. For RA, traditional (LM) and Compound Poisson (CPLM) linear models were run using log-transformed and untransformed data. All approaches used Benjamini-Hochberg (BH) p-value adjustment. RA, prevalence, and sparsity (N.not.zero) were examined across comparison groups in each dataset to identify clear misclassifications and tailor the workflow.

To ensure robust interpretation of the results, three additional DA analysis packages (also designed for compositional data) were utilized. Sparse linear phylogenetic tree-based discriminant analysis[12] (treeDA v0.0.5) was used to identify differential taxonomic clades or individual ASVs. For each analysis, four-fold cross validation was performed, and the model was fitted with the number of predictors that corresponded with the minimum cross validation error. Only ASVs with |beta coefficient| ≥0.02 were classified as differential. The ALDEx2 package (v1.38.0, 'aldex.clr') was used to estimate taxonomic variation between groups using CLR-transformed data and a Dirichlet distribution (mc.samples=1000). This analysis was run using denom="median" and denom="zero" since the latter approach was more appropriate for sparse datasets (i.e., ARKW vs "others"). Welch's t-tests were used to assess significance. A *coda-lasso* approach was also used for variable selection through penalized regression ('coda_glmnet', coda4microbiome v0.2.4) using five-fold cross validation and filtering to |beta coefficient| ≥0.05.

The results from all DA methods were then merged and examined to confirm that there were no conflicting classifications (**Supplementary Table 4**). ASVs that were identified as differential for NRKW or ARKW and "Fit Pops" were examined to determine the most appropriate classification. For ASVs designated "Fit Pops", the population with higher RA was used to color the tree and effect size in the Figures. Differential ASVs were ranked by test consensus (from 4 to 1) and MaAsLin2 effect size to identify the most important indicators for each subgroup.

#### Longitudinal diversity analyses

Diversity measures were regressed against time or age using autoregressive (AR1) general mixed-effect linear models fit by restricted maximum likelihood with 'ID' treated as a random effect (‘lme’, nlme v3.1-167); marginal $R^2$ values (the variance explained by the fixed factors) are shown. For each analysis, models with and without the autocorrelation factor (corAR1) were compared ('anova', stats) and corAR1 was retained if it significantly lowered the AIC statistic. Compositional turnover between consecutive time points ($t_1$, $t_2$…) was calculated ('turnover', codyn v2.0.5) as total turnover = ($ASVs_{Gained}$ $t_2$ + $ASVs_{Lost}$ $t_2$)/Richness $t_1$ + $t_2$[13]. Changes in mean compositional turnover were assessed using simple linear regression models ('lm', stats) to identify non-zero temporal trends. Time-lag graphs were created using Aitchison distance matrices and then by plotting pairwise compositional dissimilarities between samples collected at increasing temporal distances (“time-lags”). The SplinectomeR[14] (v0.1.0) ‘trendyspliner’ function was used to identify non-zero temporal diversity trends in one group; ‘sliding_spliner’ was used to perform sequential permutation tests and identify timeframes of significant differences between groups.

#### Identification of temporal indicator taxa

Taxa Indicator Threshold ANalysis[15] (TITAN2, v2.4.3) was performed to identify community-level change points in the SRKW microbiotas over time. TITAN2 detects changes in taxonomic abundance and occurrence along a temporal gradient and uses bootstrapping to identify indicator species. Taxon abundances are weighted by prevalence and standardized as z-scores across permutations. TITAN2 assesses taxon “purity” (the cutoff of bootstrapped runs with the same response direction, default=95%) and “reliability” (the cutoff of bootstrapped runs with p-value <0.05, default = 95%) as indicators of the gradient and classifies them as "increasers" or "decreasers". The analysis was performed using 250 permutations and 500 bootstrap replicates. Only taxa with purity and reliability scores ≥99.6% were considered robust indicator species. For population-level analyses, TITAN2 was also performed using age as the temporal gradient to distinguish age- versus time-related trends.

#### Taxonomic volatility and host survival outcome

Longitudinal taxonomic RA was examined in the microbiotas of the most frequently sampled SRKWs (n≥9) to identify “volatile” (temporally unstable with regards to baseline abundance) taxa in individuals with different health outcomes. For each SRKW, an individualized baseline of ASV abundance was created using all but their final two samples. For each ASV, RA was regressed against time ('predict', stats) and a posterior predictive distribution was used to calculate a conservative baseline (95% prediction interval * 2.5) expected to capture all new observations and identify RA outliers indicative of taxonomic volatility. ASV RA in the final two samples was then examined to identify “blooming” taxa with RA that exceeded their baseline’s upper limit. Volatility was also examined using TITAN2 and QIIME2’s ‘longitudinal feature-volatility' (QLVF) function. QLVF performs random forest regression to characterize the temporal structure of the data and identify volatile taxa that are predictive of a specific time point(s)[16]. ASVs with a QLVF importance score >5% and a net average change >1% were considered blooming.

#### Identification of fecal samples potentially collected during pregnancy or lactation

To identify SRKW females that were known to be pregnant or lactating during the study period, we leveraged records of SRKW reproductive events compiled by the Center for Whale Research (CWR). Calf birth dates were estimated by CWR researchers based on serial observations of the pregnant female and age-dependent morphological features of the calf as described in Wasser et al., 2017. Calf death dates were estimated based on necropsies or the first observation of a dam without her nursing calf. If a pregnant female (identified by direct observation or drone photogrammetry) was later observed to be thin and without a calf, it was assumed that a spontaneous abortion or perinatal mortality occurred in the interim. If the exact date of a reproductive event was unknown, the mid-point of the estimated time range of occurrence was used.

To identify samples potentially collected during a reproductive event, sequential time ranges from the estimated birth or death dates were defined to delineate, with high to low confidence, when study SRKWs were pregnant or lactating (**Supplementary Fig. 18**). These time ranges were inferred based on the duration of pregnancies (468—554 days) and lactation periods (1—3 years) previously described for aquarium-housed killer whales[17–19]. Fecal samples were then classified as “pregnant” (≤468 days before estimated calf birthdate), “likely pregnant” (467–520 days), “possibly pregnant” (521–554 days), “lactating” (≤1 year after estimated calf birthdate), or “possibly lactating” (1–3 years). These samples were excluded from most other (non-reproduction) analyses. Pregnancies were considered successful if the calf survived 1 year (the minimum age of weaning observed in the aquarium setting)[19].

#### Metagenomic library preparation

Metagenomic sequence data were generated from fecal samples that had sufficient remaining DNA, that were collected from individuals within 6 months of death, and based on the results of a preliminary 16S rRNA gene sequence analysis. Libraries were prepared for metagenomic sequencing using the Nextera DNA Flex Library Prep kit (Illumina 20018705) and sequenced to a target depth of 10Gbp per sample on one lane of an Illumina NovaSeq S4 at the DNA Services Lab, Roy J. Carver Biotechnology Center, University of Illinois at Urbana-Champaign.

#### Metagenomic assembly, binning, and bin curation

Sequencing reads were filtered for host contamination using bbduk (*qhdist=1*) with the Oorc_1.1 *Orcinus orca* genome (GCA_000331955.2) as a reference. To reduce computational complexity, each set of sequencing reads was randomly down-sampled to 50 million read pairs using seqtk (seed=7) and shuffled using fq2fa. Before sequence assembly, Illumina adapters were removed from sequencing reads using BBTools and reads were subsequently trimmed using Sickle (default thresholds)[20]. Quality-filtered reads were then assembled using IDBA-UD (--*pre_correction*)[21]. For each sample, reads were mapped back to the assembled scaffolds using bowtie2[22] to compute scaffold coverage values. Assembled scaffolds were restricted to those ≥1000 bp in length for gene prediction and binning. Genes were predicted using Prodigal (*meta* mode)[23] and predicted proteins were annotated using USEARCH and the KEGG, UniRef, and UniProt databases. 16S rRNA genes were identified using a custom script employing a hidden Markov model[24,25].

Genome binning was performed using both a manual and an automated approach. For the manual approach, scaffold information was loaded into ggKbase (ggkbase.berkeley.edu) and scaffolds were binned based on coverage, GC content, taxonomic affiliation, and inventories of bacterial ‘single copy’ genes. For the automated approach, reads were cross-mapped against every other assembly using bowtie2. Coverage tables were generated using the jgi_summarize_bam_contig_depths script (bitbucket.org/berkeleylab/metabat/src/master) and passed to MetaBAT2 (minimum contig size 1500 bp)[26] for automated bin generation. Bins derived from the manual and automated approaches were reconciled with DAS Tool[27] to create the best merged set of bins for downstream analysis.

To prepare for the bin refinement step, preliminary taxonomic classifications were assigned using GTDB-Tk[28]. Bins associated with the Candidate Phyla Radiation bacteria were separated and profiled for reduced sets of marker genes sensitive to lineage-specific losses in these groups[29]. Completeness and redundancy were calculated as the percentage of marker genes present and duplicated, respectively. These quality metrics were combined with those for all other, non-CPR bins estimated by CheckM[30]. Bins were then restricted to those with ≥25% completeness and de-replicated at 99% average nucleotide identity (ANI) using dRep to create a secondary set for manual curation. All quality-filtered bins were loaded into Anvi'o[31] and visualized individually using the *anvi-refine* command. Bins were refined by removing sets of scaffolds with aberrant coverage profiles across all cross-mapped samples. Completeness and redundancy metrics were also considered. Refined bins were then re-assessed for quality as described above.

##### Shotgun metagenomic sequence-based functional analysis

Based on the results of the 16S rRNA gene sequence analysis, two SRKW *Fusobacterium* metagenomically assembled genomes (MAGs) were selected for analysis. Comparison of ANI values was performed using the Proksee online interface[32] between both MAGs and relevant NCBI reference genomes; those with ANI >97% were considered the same species. Both MAGs were annotated with Prokka[33] (v1.14.6) and Bakta[34] (v1.8.2) using Proksee[32]. A custom virulence factor database was constructed using the Virulence Factor Database[35] (VFDB) and relevant literature[36]. Potential virulence factors were identified and sequences were extracted using Proksee, then analyzed with BLASTN and the FusoPortal[37] online database to find their closest homologues in *Fusobacterium* reference genomes.

#### Supplementary Results & Discussion:

##### Wild killer whale 16S rRNA sequence dataset

We examined taxonomic composition in the fecal microbiotas of 56 SRKW individuals (**Supplementary Table 1**) sampled from 08/12/2005 through 08/09/2019 (**Fig. 1a, b**), representing approximately 77% of the living population[38]. Of these, 42 SRKW individuals were sampled longitudinally over a duration ranging from 5 days to 13 years (**Supplementary Fig. 1**; n = 328, mean samples per individual = 8, range = 2–17). A total of 342 Southern Resident killer whale (SRKW) fecal samples were analyzed. After amplicon sequencing and quality filtering, 5,655 unique amplicon sequence variants (ASVs, mean = 85, range = 14–704) were identified in the SRKW dataset (considering 'ps_alpha' in **Supplementary Code 2**; total = 44,619,137 reads, mean = 130,465 per sample, sd = 60,664).

To provide context for interpreting patterns of the fecal microbial communities within the SRKW population, 39 fecal samples were collected from the geographically adjacent and partially sympatric NRKW and ARKW populations, from 5/30/2017 through 8/18/2019. The 13 samples collected from 9 NRKWs (mean = 1 sample per individual, range = 1-3; duration of sampling per individual = 9–363 days) yielded 161 unique ASVs (mean = 31 ASVs per sample, range = 15–67; total reads = 1,085,391; mean = 83,492; sd = 13,826). The 26 samples collected from 21 ARKWs (mean = 1 sample per individual, range = 1–3; duration = 10–364 days) yielded 1,133 ASVs (mean unique ASVs per sample = 91, range ASVs per sample = 16–477; total reads = 2,513,385, mean = 96,669; sd = 25,127). Three opportunistic samples were also collected from three marine mammal-eating transient killer whales (TKWs) that were encountered during the study; these yielded 303 ASVs (total = 482,895 reads, mean = 160,965; sd = 36,836). Combined, the wild killer whale dataset (n = 384 from 89 individuals) yielded 6,460 unique ASVs.

To examine temporal microbial stability in the killer whale host under relatively controlled conditions, we characterized the fecal microbiotas of 12 aquarium-housed killer whales (AqKWs) at two U.S. aquariums (**Supplementary Fig. 2**) over a four-month period. Fifteen individuals were initially enrolled but received antibiotics during the sample collection period and were excluded from the study. The remaining 12 individuals were clinically healthy throughout the sampling period and did not receive antibiotics. From these individuals, a total of 275 fecal samples were collected (3309 ASVs; 23,249,677 reads; mean = 66 $\pm$ 48 ASVs per sample, mean = 84,544 reads per sample; sd = 25,241; **Supplementary Code 2**).

#### The killer whale fecal microbiotas had distinct compositions from those of their surrounding water

To address the potential for bacterial cross-contamination, seawater was collected adjacent to SRKW and NRKW fecal samples during sample collection in 2018 and 2019 and analyzed in parallel. During sampling of the SRKW population, 7 surface seawater samples were collected from the Juan de Fuca Strait (JFS) (545 unique ASVs considering ps_alpha; 409,424 reads; mean = 58,489; range=37,259–85,688; sd = 21,254). During sampling of the NRKW population, 3 seawater samples were collected from Queen Charlotte Strait (QCS) (632

ASVs; 166,123 reads; mean = 55,374; range=49,715–61,406; sd = 5,854). Ordination of the data using Aitchison distances showed that the SRKW and NRKW fecal microbiotas clustered together, separate from the seawater communities (**Supplementary Fig. 3a**) (ADONIS2 $R^2$ = 55.0%, p<0.001), which clustered by sampling location. The dissimilarity between the fecal and seawater microbiotas was evident even at the phylum-level (**Supplementary Fig. 3b**); KW fecal microbiotas were dominated by Firmicutes ($RA_{SRKW}$ = 43.08%, $RA_{NRKW}$ = 92.80%) whereas seawater was dominated by Proteobacteria ($RA_{JFS}$ = 57.18%, $RA_{QCS}$ = 57.42%). There was no overlap between the most abundant ASVs in the fecal microbiotas (**Supplementary Fig. 3c, d**) and those most abundant in seawater (**Supplementary Data 3, 4**).

For the AqKWs, one pool water sample was collected at each facility (Facility A, considering ps_alpha, 455 ASVs, 79,023 reads; Facility B, 659 ASVs, 34,538 reads). The AqKW fecal microbiotas (**Supplementary Data 7a**) were also distinct from those of their pool water (**Supplementary Data 7b**) and had more similar structure to those of the wild killer whales (**Supplementary Fig. 3e, f**).

These results are consistent with those of previous studies that have also demonstrated sharp distinctions between the microbiotas of other marine mammals (e.g., harbor seals[39], sea otters[40], belugas[41], dolphins[42,43], and sea lions[42]) and their in-contact water. Together, these findings imply that the acquisition and maintenance of marine mammal-associated microbiotas largely depends on factors other than environmental filtering from water.

#### The transient killer whale fecal microbiotas were characterized by high abundance of *Paeniclostridium* ASV1 and *Clostridium SS1* spp.

Ordination of between-sample Aitchison dissimilarities of the wild KW ecotypes and populations showed that the three fecal samples collected from transient KWs (TKWs) overlapped with those of the SRKWs and separated according to the collection year with the 2010 samples clustering closer to the NRKW and ARKW populations (sampled 2017–2019) and the 2008 sample clustering with SRKW samples collected in earlier years (**Supplementary Fig. 5a**). The TKW samples had high richness, although the small TKW sample size (n = 3) reduced the power to detect statistical significance (**Supplementary Fig. 5b**).

The TKW fecal microbiotas were characterized by high abundance of *Paeniclostridium* (vs ARKW: $p$ = 0.012, **Supplementary Fig. 5c**) and universal wild killer whale taxon *Paeniclostridium* ASV1 (mean RA = 41.13%, prev = 100.0%) (**Supplementary Fig. 3f**). TKW core taxa also included *Clostridium SS1 chauvoei* ASV68 (mean RA = 11.82%), *Clostridium SS1* *septicum* ASV72 (10.72%), *Photobacterium* ASV2 (5.43%), *Mycobacterium* ASV256 (3.80%), *Tyzzerella* ASV61 (2.88%), *Paeniclostridium* ASV153 (1.31%), and ASV349 (1.08%) (**Supplementary Table 2**).

#### Wild and aquarium-housed killer whale fecal microbiotas were distinct due to the differential abundance of phylogenetically related taxa

The compositions of the AqKW fecal microbiotas were largely distinct from those of the wild resident killer whales when all samples collected between 2017–2019 were considered (**Supplementary Fig. 8a-d**) and when the data was normalized to n = 13 per subgroup (adonis2: Pr(>F) = 0.001, $R^2_{Husbandry}$ = 12.4%) (**Supplementary Fig. 8b**). The AqKW and SRKW microbiotas exhibited similar richness (**Supplementary Fig. 8c**), but distinct compositions when comparing Aitchison dissimilarity (Pr(>F) = 0.001, $R^2$ = 17.1%) (**Fig. 8e**), Bray-Curtis distance (Pr(>F) = 0.001, $R^2$ = 20.0%) (**Supplementary Fig. 8f**), and unweighted (Pr(>F) = 0.001, $R^2$ = 18.6%) (**Supplementary Fig. 8g**), and weighted Unifrac distances (Pr(>F) = 0.030, $R^2$ = 10.2%) (**Supplementary Fig. 8h**). However, no husbandry-associated clustering was observed along the primary axes using weighted Unifrac. This suggested that differences were driven by less abundant taxa, and that dominant taxa had similar abundances between groups but were represented by different, closely related ASVs. The AqKW fecal microbiotas had higher abundance of *Photobacterium* (p = 0.042) (**Supplementary Fig. 8d**) ASV2 (also classified as core in the SRKW microbiotas in **Supplementary Table 2**), ASV23, and ASV37 (**Supplementary Fig. 9, Supplementary Table 4**). *Romboutsia* (p = 0.004) (**Supplementary Fig. 8d**) also distinguished the AqKW microbiotas, but the most abundant ASV (*Romboutsia* ASV4, $rank_{DA}$ = 2, mean RA = 26.69%, prev = 100.00%) (**Supplementary Table 2**) was also highly prevalent (RA = 0.69%, prev = 95.45%) in the SRKW microbiotas. Similarly, all three wild resident populations had higher abundance of *Actinobacillus* ($p \leq 0.043$), primarily *Actinobacillus delphinicola* ASV3 ($rank_{DA}$ = 1), which was prevalent (62.91%) at low abundance in the AqKW microbiotas. Among the Enterobacteriaceae, *Escherichia-Shigella* ASV19 best differentiated the AqKW fecal microbiotas ($rank_{DA}$ = 1) (**Supplementary Fig. 9**), whereas *Edwardsiella tarda* was differential for the wild populations ($rank_{DA}$ = 3) (**Supplementary Table 4**). AqKW ASV19 sequences were identical to *Escherichia coli* (**Supplementary Data 8**).

Taxa that were differentially abundant between husbandry groups but prevalent in both could indicate host-associated taxa or taxa that impart essential, but redundant functionality to the host. *Romboutsia* and *Photobacterium* have both been described as dominant symbionts in guts of marine fish[44] and shrimp[45]; *Romboutsia* aids in amino acid and vitamin metabolism and *Photobacterium* produces fatty acids, antimicrobials[46], and chitinases to aid the digestion of crustaceans[47]. In mammals, such as cattle[48] and goats[49], *Romboutsia* is important for vitamin synthesis, carbohydrate utilization and immunomodulation. Whereas *Romboutsia* and *Paeniclostridium* often co-occur at high abundance in ruminants, here, only one was dominant in each killer whale husbandry group. This may indicate shared traits between these closely related Peptostreptococcaceae since both taxa have been shown to promote immune functions and modify bile acids in the host gut[50,51]. Such functional redundancy within the killer whale microbiotas may help maintain gut homeostasis even when environmental conditions select for different dominant taxa.

#### Temporal changes in the Southern Resident killer whale microbiotas over time

Vanishing taxa within the SRKW population could also reflect the end of transient colonization events, such as infections, should the affected individuals recover or die. For example, *Plesiomonas* ASV13 was an early core taxon (2005-2008: RA = 2.23%, Prev = 52.33%) that declined to RA = 0.03% and prevalence = 9.09% in 2017–2019. This taxon had an inverse association with host age (TITAN2) (**Supplementary Table 7**). *Plesiomonas* ASV13 was most persistent in two adult females, including J2 (10/13 samples), the oldest individual in the study, and it became undetectable in the SKRW microbiotas after her death in 2017. *Plesiomonas* ASV13 was present in 10/16 of J28's fecal samples, 8 of which were collected when she was pregnant (n = 6) or lactating (**Supplementary Data 2**). An association with advanced age or pregnancy could indicate that ASV13 opportunistically colonized immunocompromised individuals. ASV13 is identical to the corresponding gene sequence of *Plesiomonas shigelloides* (**Supplementary Data 8**), a cause of disease in fish and foodborne illness[52]. *P. shigelloides* has been identified in multiple marine mammal species and is thought to act as an opportunistic pathogen in stressed hosts[53].

#### The fecal microbiotas of Southern Residents during poor outcome pregnancies were enriched for *Fusobacterium* compared to the microbiotas of nonpregnant females.

Eleven female SRKWs were identified as having been potentially sampled during pregnancy (n = 25 samples) and/or lactation (n = 25 samples) (**Supplementary Fig. 18b**). Of these, 4 were each sampled during two or three pregnancies and 8 were also sampled at times when they were assumed to be nonpregnant and nonlactating (“nonpregnant”, n = 31 samples). *Paeniclostridium* was the dominant genus overall in the fecal microbiotas of pregnant or lactating SRKWs with good pregnancy outcomes (**Supplementary Fig. 20a-c**). However, *Fusobacterium* and *Photobacterium* were the dominant genera overall among pregnant SRKWs with poor outcomes. These results were similar to those observed in the survival group analysis (**Fig. 5a-d**) where *Paeniclostridium* and *Fusobacterium* were enriched in the “Survived” and “Dying” groups, respectively. Taken together, these findings suggest that both genera are linked to both survival and reproductive health in SRKWs. However, our reproductive dataset was limited in sample size, and the reproductive phase sampled (pregnancy versus lactation) was confounded with pregnancy outcome (“good” outcome, lactation n = 25, pregnancy n = 4; “bad” outcome, lactation n = 0, pregnancy n = 21). Since the normal microbiome of pregnant killer whales has not yet been described, it was unclear whether the observed differences in genus-level dominance reflected reproductive phase or pregnancy outcome.

At the ASV level, *Fusobacterium necrogenes* ASV11 was the only taxon indicative of SRKW fecal microbiotas collected during poor outcome pregnancies (n = 15, considering only “Pregnant” and “Likely pregnant” samples) when compared to those from “nonpregnant” females (n = 31) (**Supplementary Fig. 20d, Supplementary Table 11**). The same taxon was previously shown to have bloomed in the final samples collected from “Dying” SRKW L26 and “Died later” SRKWs K21 and J2 (**Fig. 6e**). ASV11 may therefore be influential on both reproductive and systemic SRKW health or represent a sequelum of failing health (e.g., opportunistic infection). Little information is available regarding the clinical implications of *Fusobacterium necrogenes*—although it has been associated with infections in humans, documented cases are rare[54]. ASV11 matched sequences from healthy, managed dolphins and pinniped rectal swabs[42] (**Supplementary Data 8**), which suggests it may serve as a marine mammal commensal. However, it also matched *Fusobacterium mortiferum*, which has been more commonly implicated in human disease[55,56]. A larger, more balanced sampling effort to characterize normal microbiota dynamics during killer whale reproduction could help elucidate the role of *Fusobacterium necrogenes* ASV11 in pregnancy and help identify other potential indicators of SRKW reproductive health.

#### Volatile *Fusobacterium* abundance dynamics in fecal samples collected prior to death

Of the six “Dying” SRKWs sampled within 6 months of death, longitudinal samples were available for three individuals: J8 (n = 15), L2 (n = 10), and L26 (n = 9) (**Fig. 6a**). For each SRKW, taxonomic abundance in their early fecal microbiotas (i.e., all samples excluding the last two) was used to create an individualized baseline using a linear model to predict ASV RA over time. A 95% prediction interval (PI) was calculated for each ASV and multiplied to create a conservative prediction threshold (PT = PI*2.5) for new observations. Taxa with RA>PT in the final two samples were classified as volatile. This analysis identified 5 volatile ASVs in the final samples collected from J8 and L26, and 4 volatile ASVs in the final samples from L2 (**Fig. 6b-d**). To determine whether taxonomic volatility was unique to the Dying SRKW cohort, we repeated these analyses using all other SRKWs with n≥10 (**Fig. 6e**, **Supplementary Code 2**). Three of these SRKWs (J17, K21 and J2) died during the study period but their final sample was collected >6 months (1.2-3.3 years) before death (“Died later”). A total of five volatile taxa were identified in the “Died later” cohort. Considering both “Dying” and “Died later”, *Fusobacterium* ASVs were the only volatile taxa to be identified in more than one SRKW; *Fusobacterium* ASV14 was most frequently volatile, followed by ASV11, then ASV8 and ASV10. Q2 longitudinal and TITAN2 analyses performed on all SRKW individuals except L26 confirmed the volatility of *Fusobacterium* ASV8 and ASV14 in J8’s final samples, and *Fusobacterium* ASV11 in L26’s final samples. No volatile taxa were identified in the Survived cohort (J26, J31 and J27) using all analytical methods.

The fact that more taxonomic volatility was identified in the “Dying” than in the “Died later” cohort, and volatile taxa were not identified in the surviving SRKW suggests that volatility itself is an indicator of ill health. Volatility may imply an unstable, less resilient gut microbiome that is more prone to disease-associated states. In humans and mice, volatility of the gut microbiota has been associated with chronic stress[57] and gut inflammation[58]. Here, the most frequently volatile taxa associated with dying SRKW all belonged to *Fusobacterium*. This is consistent with our earlier findings that *Fusobacterium* differentiated the endangered SRKWs from healthier populations, late from early years in the declining SRKW population, and dying from surviving SRKW health groups. Therefore, we sought to better understand the role of *Fusobacterium* in SRKW health by examining its functional potential.

#### Metagenomic characterization and virulence profiles of novel fusobacteria in the fecal microbiota of a dying Southern Resident killer whale

We recovered two metagenome-assembled genomes (MAGs) affiliated with the genus *Fusobacterium* in the final fecal sample collected from L26 (5.5 months prior to death) corresponding to a *F. mortiferum*-like ("F_mort") strain and an unclassified *Fusobacterium* ("Fuso_unclassified") strain. The two L26 MAGs were distinct at the species level (ANI = 72.44%) (**Supplementary Table 9**) and distinct from all 14 NCBI *Fusobacterium* reference genomes analyzed (ANI = 69.2–77.2%). The closest matches for F_mort were *F. hominis* (ANI = 78.9%) and *F. mortiferum* (ANI = 77.2%). The closest matches for Fuso_unclassified were *F. perfoetens* (ANI = 75.3%) and *F. mortiferum* (ANI = 73.3%). To determine whether these genomes had pathogenic potential that could have been associated with L26's declining health, we screened both against two virulence factor databases (VFDB and FusoPortal). The majority of the 77 potential VFs identified in F_mort (**Supplementary Table 10; Supplementary Data 9**) were most similar to sequences from *F. hominis* (42 VFs) and veterinary pathogen *F. mortiferum* (19 VFs). Of the 55 potential VFs identified in Fuso_unclassified, 28 were most similar to those from *F. sphaericum* in human colorectal adenocarcinoma biopsies (BioSample SAMN15202580).

Several established virulence factors were identified in both SRKW *Fusobacterium* MAGs, including those that were previously identified in other *Fusobacterium* species[36], such as genes associated with hemolysis (*hlyC/corC, tlyA, hlyD, cvfB*), biofilm formation (*pfo, dnaJ*), and capsule biosynthesis (*galE, murJ, kpsF*). Thirty-five established VFs were only identified in the F_mort MAG, including adhesins (*ompH* and *upaG*), a hemolysin transporter (*shlB*) and components of type III (*ctpA*) and IV (*virB4* and *virB11*) secretion systems. Thirteen established VFs were only identified in the Fuso_unclassified MAG, including septicolysin and hemolysis regulator *tldD*. The detection of numerous established virulence factors in both L26 *Fusobacterium* MAGs suggests there is potential for these taxa to act as primary or opportunistic pathogens in the SRKW hosts. Therefore, further exploration of their function within the SRKW gut over time using complementary methods such as metabolomics and metatranscriptomics is warranted and might help elucidate their association with host health.
